# Ultrahigh-throughput directed evolution of a metal-free α/β-hydrolase with a Cys-His-Asp triad into an efficient phosphotriesterase

**DOI:** 10.1101/2022.02.14.480337

**Authors:** David Schnettler Fernández, Oskar James Klein, Tomasz S. Kaminski, Pierre-Yves Colin, Florian Hollfelder

**Affiliations:** Department of Biochemistry, University of Cambridge, 80 Tennis Court Road, Cambridge, CB2 1GA, United Kingdom

**Author notes:** Institute of Integrative Biology, Department of Environmental Systems Science, ETH Zurich, Universitätsstrasse 16, 8092 Zürich, Switzerland. Department of Chemistry, University of Cambridge, Lensfield Road, Cambridge, CB2 1EW, United Kingdom (Oskar James Klein). Department of Environmental Microbiology and Biotechnology, Institute of Microbiology, Faculty of Biology, University of Warsaw, Miecznikowa 1, 02-096 Warsaw, Poland (Tomasz S. Kaminski). LabGenius, G01-03 Cocoa Studios, Biscuit Factory, 100 Drummond Road, London, SE16 4DG, United Kingdom (Pierre-Yves Colin).

**Keywords:** protein engineering, directed evolution, microfluidics, ultrahigh throughput screening, phosphotriesterase, catalytic mechanism, catalytic triad

## Abstract

The recent massive release of new, man-made substances into the environment requires bioremediation, but a very limited number of enzymes evolved in response are available. When environments have not encountered the potentially hazardous materials in their evolutionary history, existing enzymes have to be repurposed. The recruitment of accidental, typically low-level promiscuous activities provides a head start that, after gene duplication, can adapt and provide a selectable advantage. This evolutionary scenario raises the question whether it is possible to adaptively improve the low-level activity of enzymes recruited from non- (or only recently) contaminated environments quickly to the level of evolved bioremediators.

Here we address the evolution of phosphotriesterases (enzymes for hydrolysis of organophosphate pesticides or chemical warfare agents) in such a scenario: In a previous functional metagenomics screening we had identified a promiscuous phosphotriesterase activity of the α/β-hydrolase P91, with an unexpected Cys-His-Asp catalytic triad as the active site motif. We now probe evolvability of P91 using ultrahigh-throughput screening in microfluidic droplets, and test for the first time whether the unique catalytic motif of a cysteine-containing triad can adapt to achieve rates that rival existing phosphotriesterases. These mechanistically distinct enzymes achieve their high rates based on catalysis involving a metal-ion cofactor. A focussed, combinatorial library of P91 (> 10^5^ members) was screened on-chip in microfluidic droplets by quantification of the reaction product, fluorescein. Within only two rounds of evolution P91’s phosphotriesterase activity was increased ≈ 400-fold to a *k*_*cat*_/*K*_*M*_ of ≈ 10^6^ M^−1^s^−1^, matching the catalytic efficiencies of naturally evolved metal-dependent phosphotriesterases. In contrast to its homologue acetylcholinesterase that suffers suicide inhibition, P91 shows fast de-phosphorylation rates and is rate-limited by the formation of the covalent adduct rather than by its hydrolysis. Our analysis highlights how the combination of focussed, combinatorial libraries with the ultrahigh throughput of droplet microfluidics can be leveraged to identify and enhance mechanistic strategies that have not reached high efficiency in Nature, resulting in alternative reagents with a novel catalytic machinery.

**GRAPHICAL ABSTRACT:** 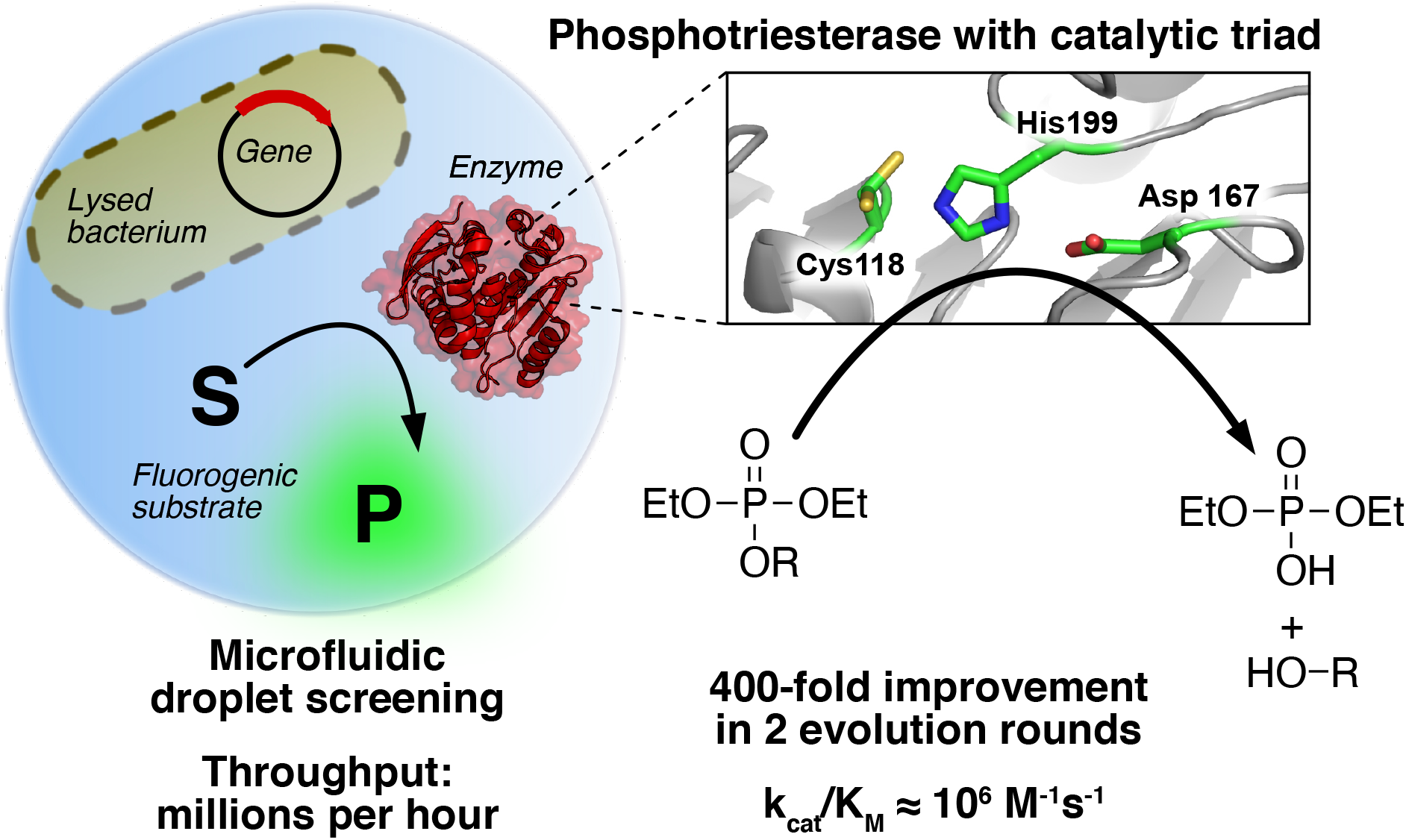

## INTRODUCTION

Organophosphate nerve agents range among the most toxic synthetic substances known. They act as potent covalent inhibitors of the enzyme acetylcholinesterase in human synapses, derailing synaptic transmission of nerve signals and leading to paralysis by seizures and subsequent death by respiratory arrest. This toxic potential has been exploited by their use as chemical warfare agents (e.g. sarin, VX, or novichok). In addition, the global use of phosphotriesters as pesticides in agriculture leads to thousands of hospital admission with organophosphate poisonings. However, the current medical treatment of organophosphate damage is limited to mitigating primary symptoms.^1–3^ While existing organophosphate-degrading enzymes hold prophylactic and therapeutic promise as catalytic bioscavengers,^4–6^ further organophosphate-degrading enzymes are needed as options for the detoxification of the large variety of organophosphates that require therapeutic and bioremediation interventions.^7^

From an evolutionary point of view, the massive release of this xenobiotic substance class into the environment (only starting in 1947)^1^ provides a unique opportunity to observe the emergence of new biocatalytic solutions to ecological challenges. It is well established that new enzymatic activities can arise from pre-existing promiscuous activities that give a head start to adaptive evolution.^2^ Here too, in an astonishing showcase of rapid convergent evolution, existing enzymes have adapted to the efficient hydrolysis of organophosphates within just decades, arising from diverse protein superfamilies, such as the amidohydrolases,^3^ the pita-bread fold,^4^ the β-propellers,^5,6^ and the metallo-β-lactamases.^7^ Remarkably, despite their diverse evolutionary origins, all these enzymes have independently come to the same mechanistic solution, requiring bivalent cations (Zn^2+^, Ca^2+^, Mn^2+^, Co^2+^) as a cofactor for metal-ion catalysis.^8^ Other promiscuous enzymes with this metal-dependent catalytic motif have also been evolved or engineered to high phosphotriesterase activities (mirroring the scaffolds which adapted naturally: β-propellers,^9,10^ amidohydrolases^11^ and metallo-β-lactamases^12^). In each case the emerging new activity relied on the intrinsic promiscuous reactivity of a metal ion that provides Lewis acid catalysis, transition state charge compensation and coordination of a water molecule with reduced p*K*a to facilitate its use as a nucleophile in hydrolysis.

Our previous work^20^ had identified a second, metal cofactor-free catalytic motif by ultrahigh throughput screening of a >10^6^-membered naïve metagenomic library generated from soil, marine sludge and cow rumen samples that had not been exposed to phosphotriester contamination. This functional metagenomic screen elicited P91, a member of the α/β hydrolase superfamily and distant homologue of acetylcholinesterase, with a a Cys-His-Asp triad as the catalytic motif (**Figures 1 and 3**).^20^ P91’s ability to hydrolyse phosphotriesters is surprising, given that phosphotriesters have been specifically designed to covalently inhibit the active-site nucleophile of similar catalytic triads, like the Ser-His-Glu triad of synaptic acetylcholinesterase. The flipside of the suitability of triesters as inhibitors is that they will make difficult substrates for triad-bearing enzymes. Indeed, no metal-independent, triad-bearing phophotriesterase has achieved comparably high efficiency, raising the question whether (and if yes, how) a catalytic motif without a metal can be improved. To take up this challenge we subjected P91 to directed evolution – making use of the ultrahigh throughput of microfluidic droplet screening (>10^6^/h) – and identified a variant able to hydrolyse phosphotriesters at catalytic efficiencies rivalling those of many metal-dependent phosphotriesterases. The success of this adaptation by directed evolution suggests that a Cys-containing catalytic triad is an evolvable motif for this new function, in principle set-up for efficient hydrolysis of organophosphates, even though such adaptation has never been observed in Nature.

**Figure 1:**
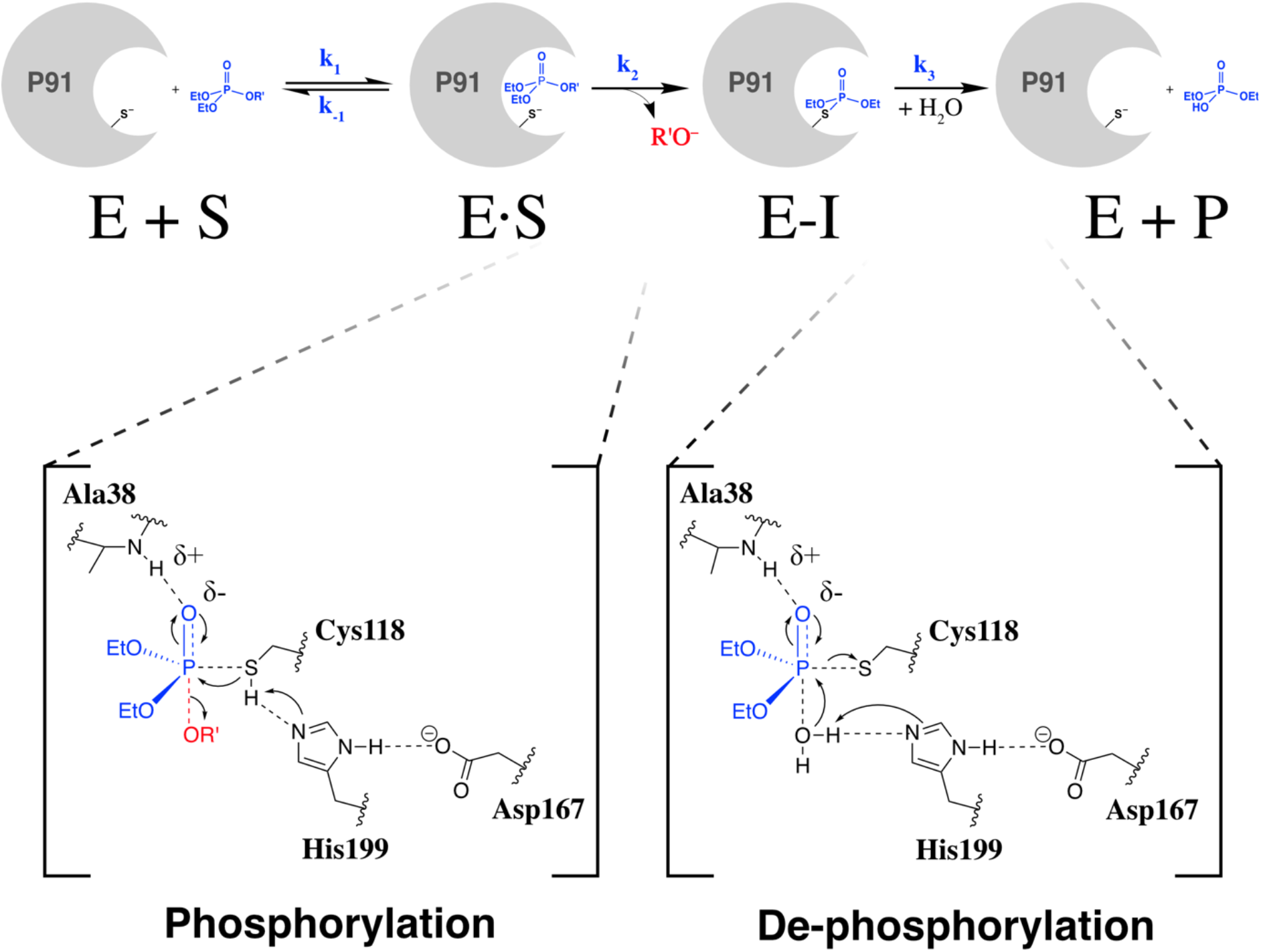
Overview of the assumed reaction scheme and the transition states. P91 hydrolyses phosphotriesters via a mechanism involving the formation (transition state 1, *k*_*2*_) and breakdown (transition state 2, *k*_*3*_) of a covalent intermediate (E-I). Both transition states have a trigonal-bipyramidal geometry around the phosphorus centre during the formation of the covalent adduct and the breakdown of the covalent intermediate. His199 and Asp167 form a charge relay system, with His199 acting as a general acid/base catalyst. Ala38 contributes with its backbone nitrogen to stabilisation of the oxyanion and forms the so-called ‘oxyanion hole’. Note that this reaction scheme assumes fast product release (EP **→** E+P).

**Figure 2:**
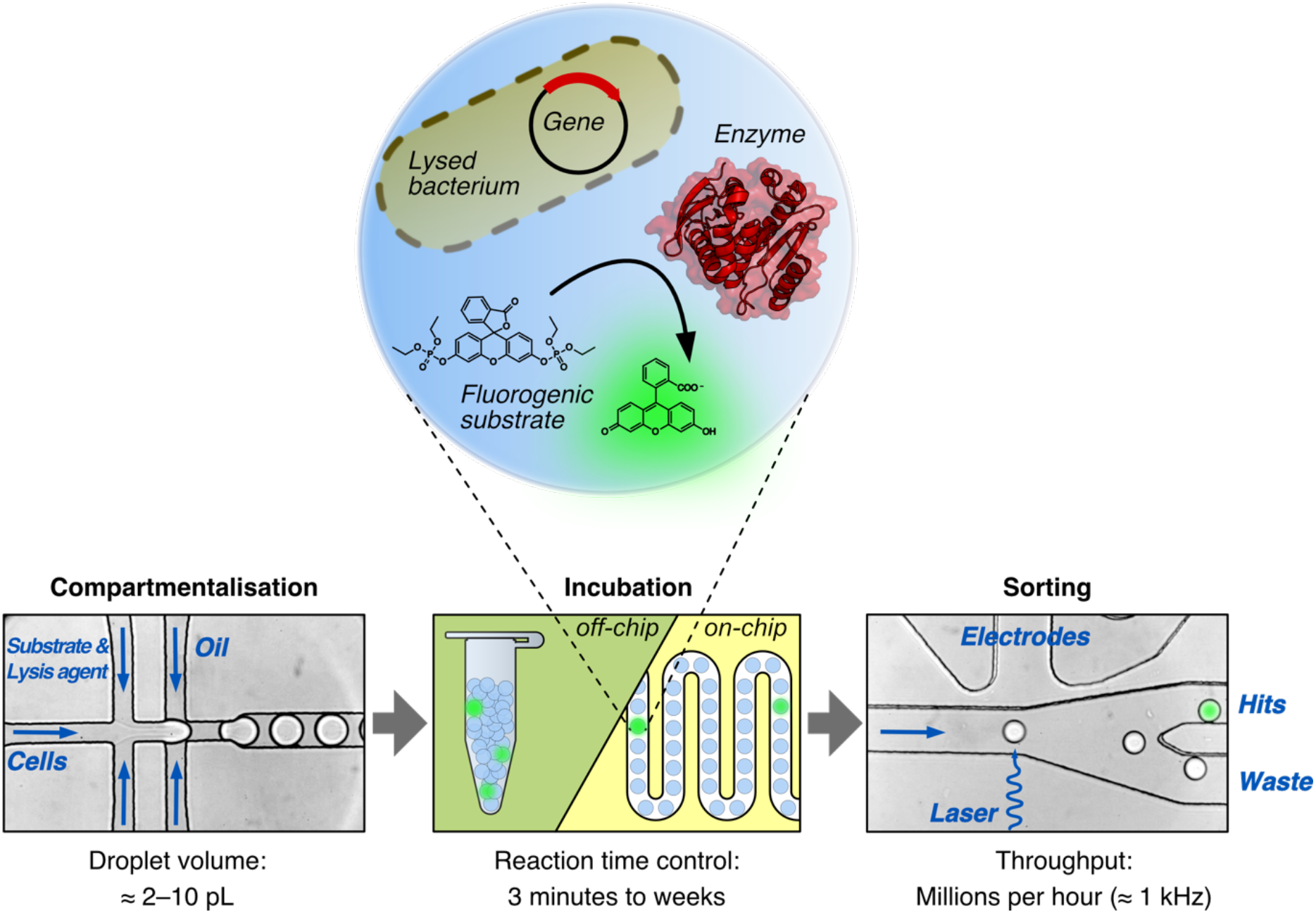
Microfluidic droplet screening assay. The microfluidic screening assay consists of three steps: (1) Encapsulation of bacterial cells together with a fluorogenic substrate and lysis agent into picolitre-sized aqueous droplets that are separated with fluorinated oil. The droplets serve as miniaturised reaction vessels that link phenotype (catalytic activity indicated by fluorescence), to genotype (gene sequence encoded on a plasmid). (2) Droplet incubation can be carried out in a delay line on-chip (for the range of minutes) or off-chip (for hours to weeks). (3) The droplets can then be sorted according to their fluorescence with an excitation laser that is focussed on the droplet flow along a Y-shaped junction. When surpassing a pre-set fluorescence threshold, a single droplet can be electrophoretically deviated by the electrodes away from the waste channel into the hit collection channel. Additional chip details are given in **Figures S2 and S3**.

**Figure 3:**
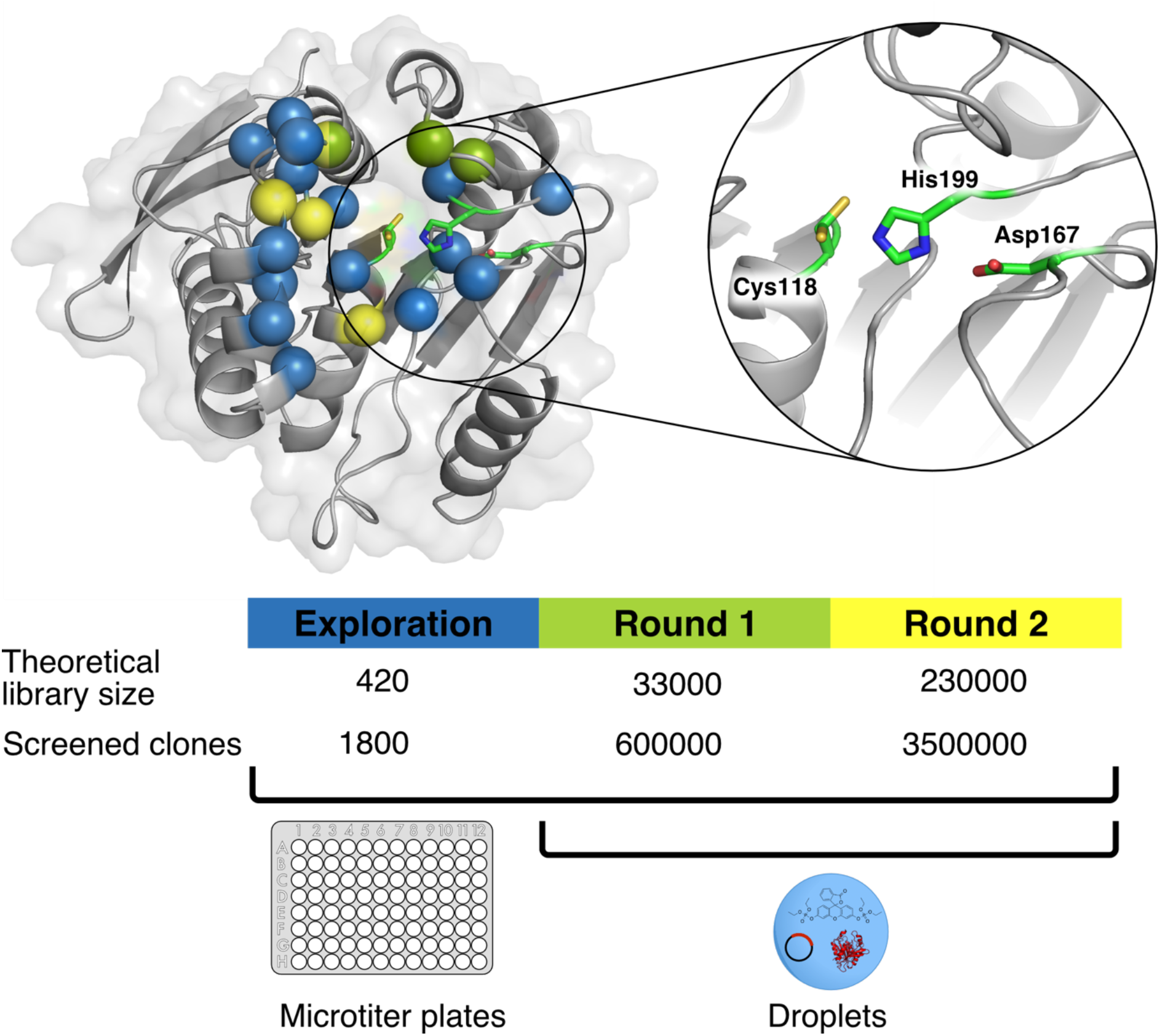
Library strategy for the directed evolution of P91. The active site of P91 was first mutationally explored (all spheres) by screening of single-site saturation libraries for phosphotriesterase activity in multititer plates (see Figure S2). A subset of these residues was then combined in round 1 (green spheres) and round 2 (yellow spheres) into combinatorial multiple-site saturation mutagenesis libraries which were screened in microfluidic droplets. In round 1, droplets were incubated off-chip whereas in round 2, droplets were incubated in a delay line on-chip in order to maintain selection stringency by shortening the reaction time. **Blowout**: Catalytic triad of P91, consisting of Cys118, His199, and Asp167. Note the two conformations of Cys118 in the structure, an inwards-pointing protected and an outwards-pointing active conformation.

## RESULTS AND DISCUSSION

### Mutational scanning identifies the catalytic potential of residues near P91’s active site

In order to remodel P91’s active site to proficiently turn over phosphotriesters, we investigated how substitutions in P91’s active site would affect activity in a mutational scanning process. The protein mutability landscape concept^21^ and our previous work^22^ suggest that an exploration of catalytic potential can help to design a relatively small library that is amenable to plate screening. All 23 residues that comprise the first shell around the catalytic triad, lining the catalytic site within up to 12 Å of the active-site nucleophile Cys118, were individually completely randomised. We determined the maximal increase in lysate activity upon mutation and ranked the residues according to the observed effects (**Figure S2**). Based on screening for hydrolytic activity towards the fluorogenic model phosphotriester substrate **1** (fluorescein di(diethylphosphate), FDDEP; **Figures 2 and S1**) we identified three positions most amenable to significant rate enhancements: Ala73, Ile211 and Leu214 (each increasing activity up to ≈ 8–10-fold in cell lysate). These three residues are in direct vicinity with each other and line the upper edge of the active site (**Figure 3**, green spheres). They are situated in loops which are partly covering the active site, highly variable in sequence and length within the dienelactone hydrolase-like protein family, to which P91 is attributed by sequence homology. Based on the recognition of their roles as crucial for substrate specificity and substrate-induced activation,^23–26^ we selected these residues for simultaneous randomisation and constructed a combinatorial library P91-A with a theoretical size of ≈ 33,000 members (NNK codes for 32 codons, three positions randomised: 32^3^ = 32 768) using the degenerate codon NNK and transformed it into *Escherichia coli*.

### Droplet screening of P91 variants leads to a 400-fold improvement in two rounds

We screened the resulting library for phosphotriesterase activity using ultrahigh-throughput droplet sorting. To this end, the library was expressed in *E. coli* cells which were encapsulated together with substrate and lysis agent in picolitre-sized microdroplet compartments on a microfluidic chip. After 2–3 h of incubation, the droplets were re-injected into a separate sorting chip where they were screened and sorted according to their fluorescence at kHz frequencies. Due to the high throughput, the entire library could be confidently sampled (≈ 3.6-fold oversampling at 20 % droplet occupancy) by screening ≈ 600 000 droplets in less than one hour. Of these, ≈ 10 000 droplets were selected, corresponding to the top ≈ 1.7 % brightest droplets. These relatively permissive sorting conditions were chosen to avoid losing false negative clones due to phenotypic variation in enzyme expression on the single cell level, allowing cumulative enrichment of improved clones over several rounds of sorting. This enrichment process by droplet sorting was carried out three times in total in order to gradually narrow down the diversity of the library before proceeding to secondary screening in microtiter plates. To exert selective pressure on *k*_*cat*_/*K*_*M*_, we chose a low substrate concentration of 3 μM (≈ 1/10 *K*_*M*_).

Following droplet screening, ≈ 90 randomly picked individual clones from each sorting (≈ 350 clones for the third, final sorting) were analysed in a lysate-based microtiter plate screening, revealing a successive enrichment of active clones along the course of sorting. The most active clones were sequenced, revealing that all sequenced clones had a tryptophan in position 211 and a valine in position 214. Position 73, in contrast, was more varied among enriched variants. Therefore, we constructed a library (P91-B) for the next round of evolution based on the consensus sequence Trp211, Val214 (dubbed P91-R1) and kept position 73, that had shown no clear consensus, randomised. We additionally randomised three further residues ranking next in terms of activity change upon individual mutation (Ala38, Leu76, and Ala122, **Figure S2**). Ala38 and Leu76 are in direct vicinity to the residues already randomised in the first library, seaming the active site and therefore have a high potential of interacting with these. In the canonical esterase mechanism, Ala38 contributes with its backbone nitrogen to stabilisation of the oxyanion and forms the so-called ‘oxyanion hole’ (**Figure 1**). Ala122 is directly below the active site nucleophile and was hypothesised to be involved in positioning the hydrolytic water molecule for the breakdown of the covalent intermediate. We randomised these positions using a mixture of the degenerate codons NDT, VHG, and TGG, which code for all canonical amino acids at approximately even representation and no stop codon (known as 22-codon trick^27^), yielding a theoretical diversity of 160 000 (= 20^4^) on the amino acid level and ≈ 234 000 (22^4^ = 234 256) on the nucleotide level.

To adjust for the shorter reaction time required for the improvement of mutants of P91-R1, we designed an integrated device on which droplet generation, incubation, and sorting were combined on a single chip^28,29^ (**Figure S4**) so that stringent sorting was possible in the second round. On this chip, the incubation time of the droplets was precisely controlled by the length of the delay line.^30^ To optimise the length of the delay line for stringent sorting, we measured the reaction progress of the starting variant P91-R1 in a chip with an elongated delay line (**Figures S4a and S5a**). The required delay line length for the library sorting device was then chosen such that sorting would occur in the early linear phase of the reaction, corresponding to a reaction time of only 5 minutes (**Figures S4b and S5b**).

As in the first round, droplet sorting was again repeated three times in total to gradually enrich active clones. In each sorting step, ≈ 1.5–3.5 million droplets were screened (≈ 5-fold oversampling of the library) and ≈ 10 000–70 000 droplets were sorted. To balance throughput with accuracy, the average droplet occupancy was continuously reduced from 35 % in the first sorting, to 20 % and then 10 % in the last sorting. Droplet sorting was again followed by a secondary microtiter plate screening (≈ 350 randomly picked clones) and sequencing of the most active clones.

### Evolved P91 variants rival the efficiencies of metal-dependent phosphotriesterases

The most improved variant, P91-R2, has five mutations compared to wild-type P91: Ala38Leu, Ala73Glu, Leu76Val, Ile211Trp, Leu214Val. A comparison of the structure of P91-WT and a structural model of P91-R2 (in an AlphaFold2/ColabFold rendition^14,15^) suggests a reshaping of the active site volume by an increase of bulk by Trp211 and slight enlargement near the catalytic triad by the Leu214Val mutation. A new hydrogen bond between Glu73 and Trp211 may rigidify this arrangement, amounting to a tighter fit^16^ and the catalytic triad’s His199 is subtly repositioned with respect to the reshaped active site.

The kinetic characterisation of purified P91-R2 revealed a ≈ 400-fold increase in *k*_*cat*_/*K*_*M*_ over wild-type (**Figures 4a, 4b** (**bars a & b**), **S6, and Table 1**), with *k*_*cat*_/*K*_*M*_ ≈ 8 × 10^5^ M^−1^s^−1^, achieved in just two rounds of evolution (**Table 1**). P91-R2 shows a greater propensity for substrate inhibition, possibly a consequence of adaptation to the low substrate concentrations (3 μM) used during screening. Despite only two rounds of evolution departing from a weakly promiscuous starting point, P91-R2’s catalytic parameters fall in a range shared with many physiological enzymes shaped by long-term natural Darwinian evolution.^17^ When compared to metal-dependent organophosphate-degrading enzymes that have either naturally evolved (isolated from organophosphate-contaminated environments) or were engineered in the laboratory (by directed evolution or rational choice) from promiscuous enzymes, the rates of P91-R2 matches most of them and falls within two orders of magnitude of the best. The fastest known metal-dependent phosphotriesterase was isolated from *Brevundimonas diminuta* (*Bd*PTE), which achieves catalytic efficiencies approaching the diffusion limit (**Figure 4b, bar l**).^18^ *Bd*PTE has been the target of numerous directed evolution campaigns, mainly towards chemical warfare agents, but has also been further evolved towards paraoxon hydrolysis, reaching an even higher *k*_*cat*_, however with low further increases in overall *k*_*cat*_/*K*_*M*_^19^. It should be noted that the catalytic efficiency of *Bd*PTE is exceptional while most naturally evolved or artificially engineered metal-containing phosphotriesterases have efficiencies in the range of 10^4^–10^6^ M^−1^s^−1^ (**Table S2**).^13^ The *k*_*cat*_/*K*_*M*_ of P91-R2 is similar to or even surpasses the efficiencies of these metal-dependent phosphotriesterases (including their evolved mutants obtained after comparatively more rounds of evolution than shown here for P91-R2; **Figure 4b**).

**Table 1:** Steady-state catalytic parameters for phosphotriester. hydrolysis by His_6_-tagged P91-WT, P91-R1, and P91-R2, measured with the substrate FDDEP in 50 mM HEPES-NaOH, 150 mM NaCl, 1 mM TCEP, pH 8.0 at 25 °C. Enzyme concentrations were: 0.2 μM for P91-WT, 1 nM for P91-R1, and 0.2 nM for P91-R2.

| Enzyme variant | Mutations | $k_{cat}$<br>(s<sup>-1</sup>)<sup>a</sup> | $K_M$<br>(μM)<sup>a</sup> | $K_i$<br>(μM) | $k_{cat}/K_M$<br>(M<sup>-1</sup>s<sup>-1</sup>)<sup>b</sup> |
| --- | --- | --- | --- | --- | --- |
| P91-WT | -- | 0.054 | 27 | 470 | $2.0 \times 10^3$ |
| P91-R1 | Ile211Trp, Leu214Val | 37 | 120 | 7.9 | $3.0 \times 10^5$ |
| P91-R2 | Ala38Leu, Ala73Glu,<br>Leu76Val, Ile211Trp,<br>Leu214Val | 15 | 20 | 14 | $7.8 \times 10^5$ |

<sup>a</sup> Due to strong substrate inhibition, estimates of $k_{cat}$ and $K_M$ for P91-R1 and P91-R2 are extrapolations and only $k_{cat}/K_M$ can be regarded as precise (see Michaelis-Menten plot, **Figure S6a**).
<sup>b</sup> The values measured for P91-WT differ from previously published values ( $< 5$ -fold),<sup>13</sup> which may be ascribed to a different affinity tag (here a N-terminal His<sub>6</sub>-tag had replaced an N-terminal StrepII-tag in reference<sup>13</sup>), and differences in purification procedure and buffer conditions.

**Figure 4:**
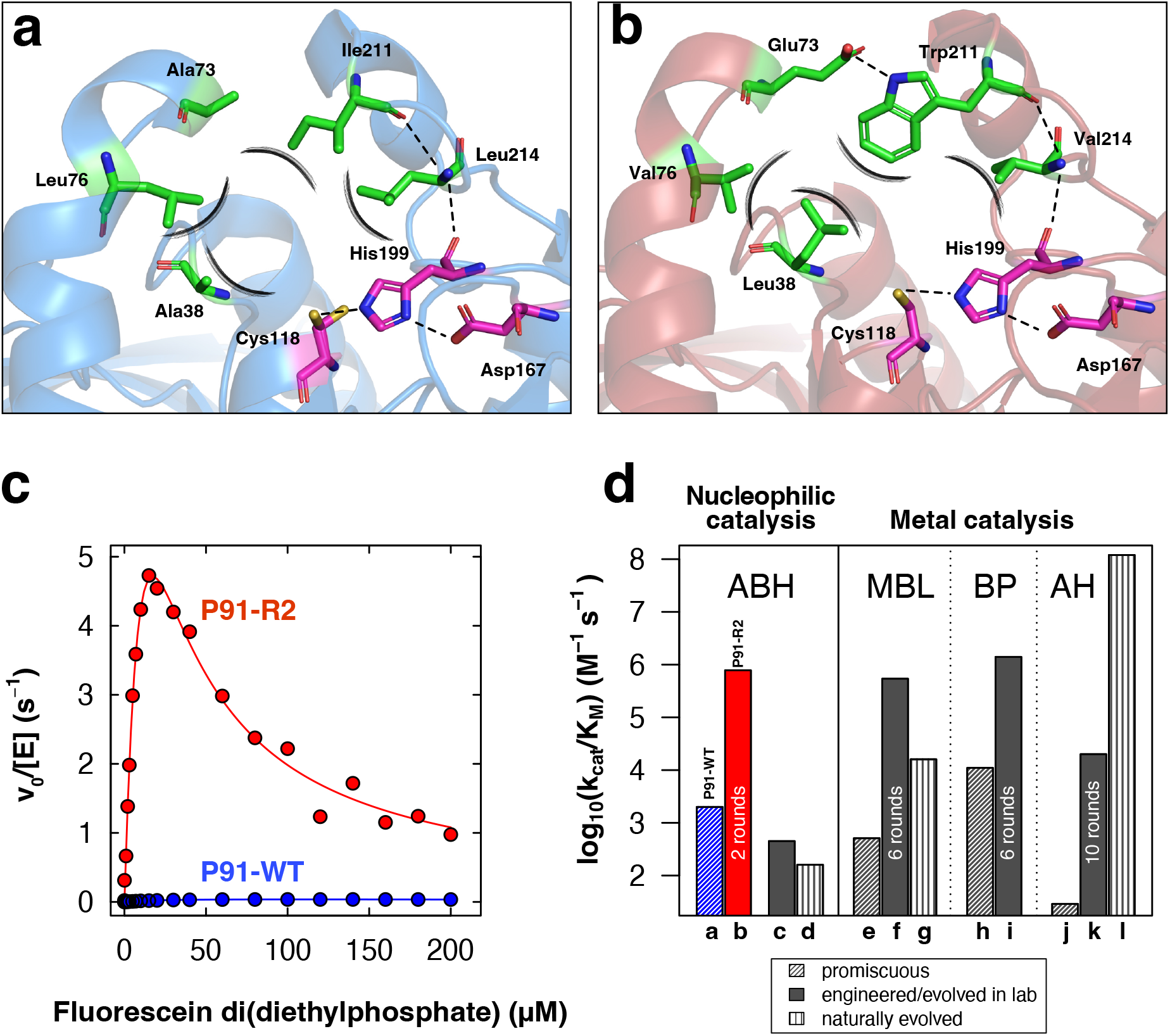
Directed evolution of P91 brings about the mutant P91-R2 that rivals the efficiencies of engineered and naturally evolved metal-dependent phosphotriesterases. **(a)** Positions identified in the preliminary mutational scanning shown in the structure of P91-WT (blue). The catalytic triad is shown in magenta and positions chosen for mutation in the library in green. Hydrogen bonds are highlighted as dotted lines, bulky protrusions into the active site as brushy lines. **(b)** The mutations in P91-R2 (red) are shown in a structure created by AlphaFold2/ColabFold.^14,15^ **(c)** Michaelis-Menten plots for P91-WT (blue) and the evolved variant P91-R2 (red) for the hydrolysis of fluorescein di(diethylphosphate) (FDDEP, 1), measured in 50 mM HEPES-NaOH, 150 mM NaCl, 1 mM TCEP, pH 8.0 at 25 °C. v_0_, initial reaction velocity; [E], initial enzyme concentration. Enzyme concentration was 0.2 μM for P91-WT and 0.2 nM for P91-R2. **(d)** Comparison of catalytic efficiencies of promiscuous (hashed), engineered (filled), naturally evolved (lined) phosphotriesterases from different protein superfamilies: ABH, α/β-hydrolases; BP, β-propellers; MBL, metallo-β-lactamases; AH, amidohydrolases. P91-WT is shown in blue, the evolved variant P91-R2 in red. For enzymes that were evolved by directed evolution, the number of rounds is indicated in the bar. Annotation for the bar labels a–l and the respective references are detailed in **Table S2**. Substrates between enzymes differ in leaving group (*p*-nitrophenol, fluorescein, and umbelliferone) but are all diethyl-substituted phosphotriesters as these are among the most common organophosphate insecticides in agricultural use and, due to their high accessibility compared to highly regulated warfare agents, are used in most enzyme characterisations and directed evolution studies in the published literature. When evolved variants showed higher activity towards a different organophosphate substrate used in the respective study, this is additionally noted in **Table S2**.

Organophosphate hydrolase activity has been identified in a range of enzymes comprising diverse protein superfamilies, such as the amidohydrolases^10^, the pita-bread fold^11^, the β-propellers^12,13^, and the metallo-β-lactamases.^14^ Despite their different fold architectures, a metal cofactor-dependent mechanism of degradation is their common feature, typically involving Zn^2+^, Ca^2+^ or other bivalent cations. Directed evolution of these enzymes from promiscuous precursors has been successful: a promiscuous member of the amidohydrolase superfamily (with lactones as the best hydrolytic substrate) has been improved in 10 rounds towards higher organophosphate hydrolase activity, reaching an efficiency 2 × 10^4^ M^−1^s^−1^ of for paraoxon-ethyl and 1.1 × 10^6^ M^−1^s^−1^ for a methyl phosphonate (**Figure 4b, bars j & k, and Table S2**).^18^ Promiscuous phosphotriesterases from the β-propeller and the metallo-β-lactamase superfamilies have also been subjected to extensive laboratory evolution, reaching catalytic efficiencies in the range of 10^4^–10^6^ M^−1^s^−1^ within 6 rounds (**Figure 4b, bars e, f & i and Table S2**).^17,19^ For diethyl-substituted phosphotriester substrates, improvements between 690-fold for an amidohydrolase,^18^ 1100-fold for a metallo-β-lactamase^19^ and 130-fold for a β-propeller^17^ were achieved in 10, 6, or 6 rounds, respectively (**Table S2**). Thus, more rounds of evolution were necessary in these previous campaigns to achieve similar improvements compared to only two here, due to the ultrahigh throughput of droplet microfluidics, and similar absolute activities were reached.

### The effects of directed evolution on a two-step mechanism

Given the rapid attainment of efficient phosphotriesterase activity of P91 within only two rounds of directed evolution, we investigated the molecular basis of its unprecedented mechanism. In analogy to the textbook case of serine proteases,^40^ a mechanism that involves a covalent intermediate had been postulated for P91 (**Figure 1** and comment in the **Supplementary Information** on evidence for the presence of an intermediate).^20^ A quantification of the rates of formation (*k*_*2*_) and breakdown (*k*_*3*_) of the intermediate will allow a comparison with the catalytic triads of other enzymes (e.g. the Ser-His-Glu triad in acetylcholinesterase and many targets beyond^20^), where a fast and near-irreversible formation of a covalent adduct leads to their inactivation in single-turnover fashion, depriving these enzymes of their physiological function. In the case of this fatal suicide reaction the phosphorylation rate *k*_*2*_ is much larger than the de-phosphorylation rate *k*_*3*_ and *k*_*3*_ ≈ 0. For a multiple-turnover enzyme we expect a larger *k*_*3*_. However, time courses for P91-WT and its mutants showed monophasic kinetics that could be fitted to a single exponential increase in reaction product (initially linear saturation curve), even when measured in a stopped-flow apparatus (**Figure S7**). Thus, the rates of phosphorylation and de-phosphorylation, *k*_*2*_ and *k*_*3*_ were not directly measurable due to lack of an observable burst.^41^ This means that P91, in contrast to acetylcholinesterase, is not rate-limited by the breakdown of its intermediate, but the formation of the covalent adduct.

A nucleophile exchange from cysteine to serine was then made to alter the rate-determining step by offering an alternative nucleophile with different reactivity. This mutant (Cys118Ser) had 10^2^–10^3^-fold lower second order rates, consistent with the idea that better availability of a deprotonated nucleophile (due to the lower p*K*_*a*_ of cysteine *vs.* serine) is important for catalysis. Cys118Ser mutants revealed biphasic kinetics, suggesting that now intermediate formation is rate-limiting and offering the possibility to quantify the relative effects of the directed evolution campaign on *k*_*2*_ and *k*_*3*_.^21^ Pre-steady state burst kinetics could be observed for nucleophile mutants of the wild-type (P91-WT Cys118Ser) and of the evolved variant (P91-R2 Cys118Ser) (**Figures 5a and S8**) and fitted to a two-step model, describing a fast intermediate formation followed by its breakdown with quantification of *k*_*2*_ and *k*_*3*_ (**Figures S8–S10**). Notably, the equipment required to measure rates of the initial burst phase (*k*_*2*_) reflected the large differences between the respective wild-type-based (microtiter plate reader) and the evolved variants (stopped-flow apparatus; **Figure 5a**). The rate of intermediate formation *k*_*2*_ increased by ≈ 900-fold from the P91-WT to the evolved P91-R2 background (**Figure 5b**). *k*_*2*_ was rate-limiting in the original cysteine-bearing wild-type enzyme and the similar increase in *k*_*cat*_/*K*_*M*_ of 400-fold in *k*_*cat*_/*K*_*M*_ in the evolved mutants suggests that adaptive evolution manifested itself predominantly in faster intermediate formation *k*_*2*_. This provides a molecular explanation for the observed improvements in the directed evolution experiment and implies that the primary adaptive correction for the promiscuously catalysed reaction was the encounter of the enzyme’s nucleophile and the substrate, while the reactivity of the cysteine-bound intermediate was already sufficient to break down relatively rapidly to avoid its build-up.

**Figure 5:**
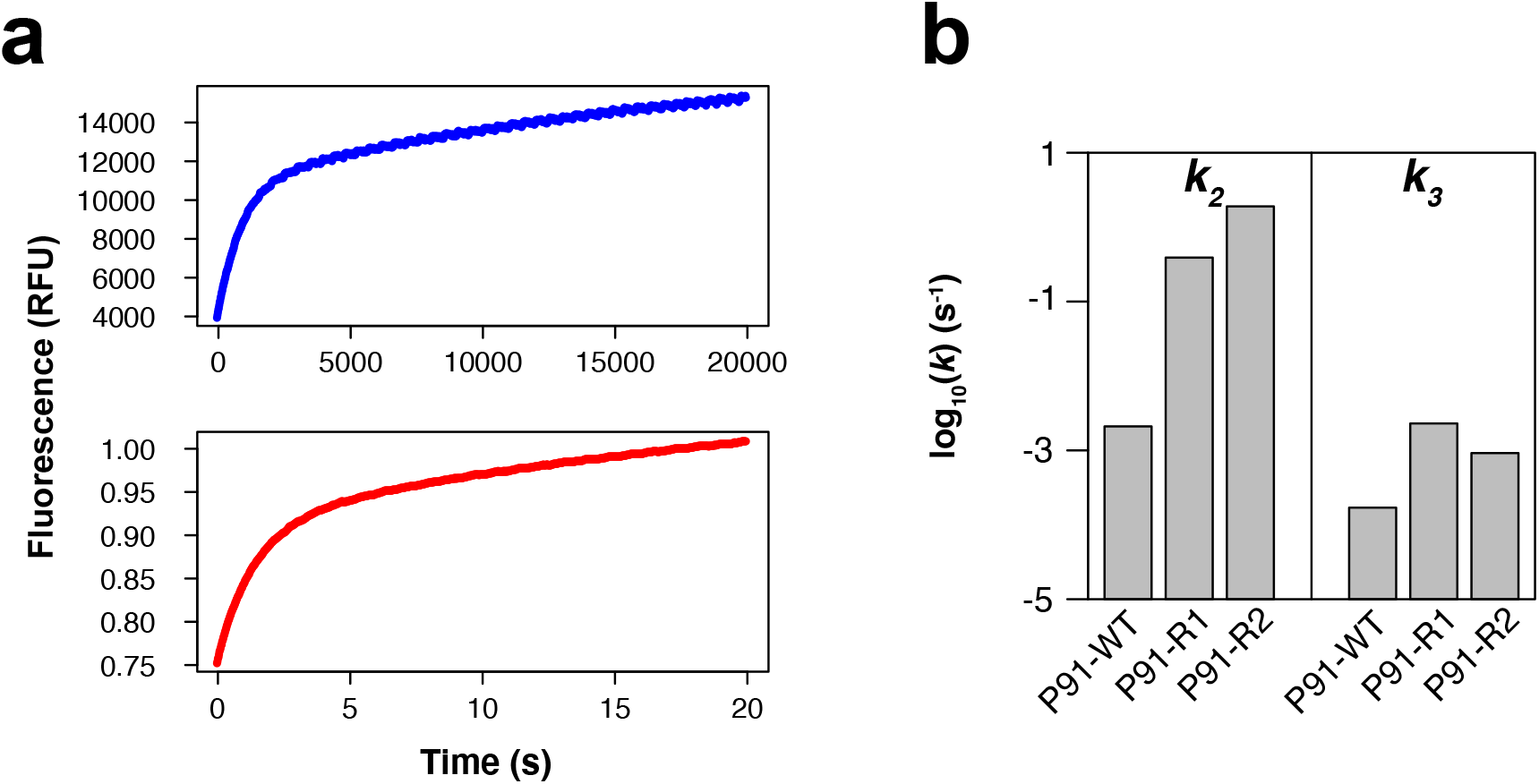
The largest effect of evolution is on intermediate formation. (**a)** Examples of traces of kinetic bursts with nucleophile-exchanged P91 variants (Cys118Ser). Reaction time course of P91-WT Cys118Ser (blue, enzyme concentration 10 μM) and P91-R2 Cys118Ser (red, enzyme concentration 1 μM) with 100 μM FDDEP in 50 mM HEPES-NaOH, 150 mM NaCl, pH 8.0 at 25 °C. Note the different time scales for the different variants, allowing the use of a spectrophotometric microplate reader for the wild-type-derived enzyme while requiring the use of a stopped-flow instrument for the evolved variant (with each instrument using different relative fluorescence units, RFU). **(b)** Individual rate constants of nucleophile-exchanged P91 variants (Cys118Ser) were determined from burst kinetics. The phosphorylation rate (*k*_*2*_) was measured with FDDEP (1) and de-phosphorylation rate (*k*_*3*_) was measured with paraoxon-ethyl (2) (see explanation in Supplementary Information). While *k*_*2*_ changed by almost three orders of magnitude over evolution, *k*_*3*_ remained approximately within the same order of magnitude.

### Specificity analysis of P91 with the native cysteine triad suggests a large increase in the rate of intermediate formation

When phosphotriester and acyl ester reactions are compared, changes in transition state geometry or in the leaving group will affect *k*_*2*_ and *k*_*3*_ differently. While reaction-type specificity is determined by the enzyme’s adaptation to the transition state geometry of a reaction (*k*_*2*_ and *k*_*3*_), leaving-group preference is only determined by the rates of formation of the Michaelis complex (*k*_−*1*_/*k*_*1*_) and nucleophilic attack on the substrate (*k*_*2*_) (**Figure 1**). Therefore, we expect that major changes in intermediate hydrolysis (*k*_*3*_) would only be reflected in reaction type specificity and not in leaving-group preference. In contrast, changes in the rate of intermediate formation (*k*_*2*_) would affect both reaction type specificity and leaving group preference. Thus, we determined arylbutylesterase and phosphotriesterase activity for two different leaving groups, *p*-nitrophenol and fluorescein, respectively (**Figures S1 and S6, Table S1**). This was possible because P91-WT has comparable catalytic parameters for the hydrolysis of carboxyesters (with a tetrahedral transition state) in addition to its phosphotriesterase activity (trigonal-bipyramidal transition state). Wild-type P91 is a slightly better carboxyesterase than phosphotriesterase for both leaving groups. It also has a slight preference for fluorescein over *p*-nitrophenol as a leaving group.

We observe that over the course of the directed evolution campaign, P91 specialises for both the fluorescein leaving group *and* for phosphotriester hydrolysis (**Figure 6**). Remarkably, this specialisation towards phosphotriesterase function at the cost of carboxyesterase activity is strongly leaving-group dependent: the increase in phosphotriesterase activity is much more pronounced with fluorescein (≈ 400-fold) than with *p*-nitrophenol (≈ 4-fold) as a leaving group. Carboxyesterase activity, in contrast, decreased roughly equally (≈ 3.6 and ≈ 7.6-fold, respectively; **Figure S6, Table S1**) with both leaving groups. The preference for fluorescein as a leaving group is therefore linked to the phosphotriester transition state (affecting *k*_2_), and thus rules out a simple ground state binding effect (i.e. a decrease of *K*_*d*_ as the consequence of changing the ratio of *k*_*1*_ and *k*_−*1*_). This finding is consistent with a significant increase in *k*_*2*_ rather than in *k*_*3*_ and confirms that the previous rate measurements in the nucleophile-exchanged variants were representative of a mechanism in which directed evolution strongly accelerated intermediate formation in P91. It also validates that the effects observed in the nucleophile-exchanged enzyme variants can be applied to P91 with a cysteine nucleophile. Apparently breakdown of the intermediate (*k*_*3*_) is so fast it does not need improvement, implying that this step is effectively optimised (compared to an alkoxy leaving group in serine-containing hydrolases).

**Figure 6:**
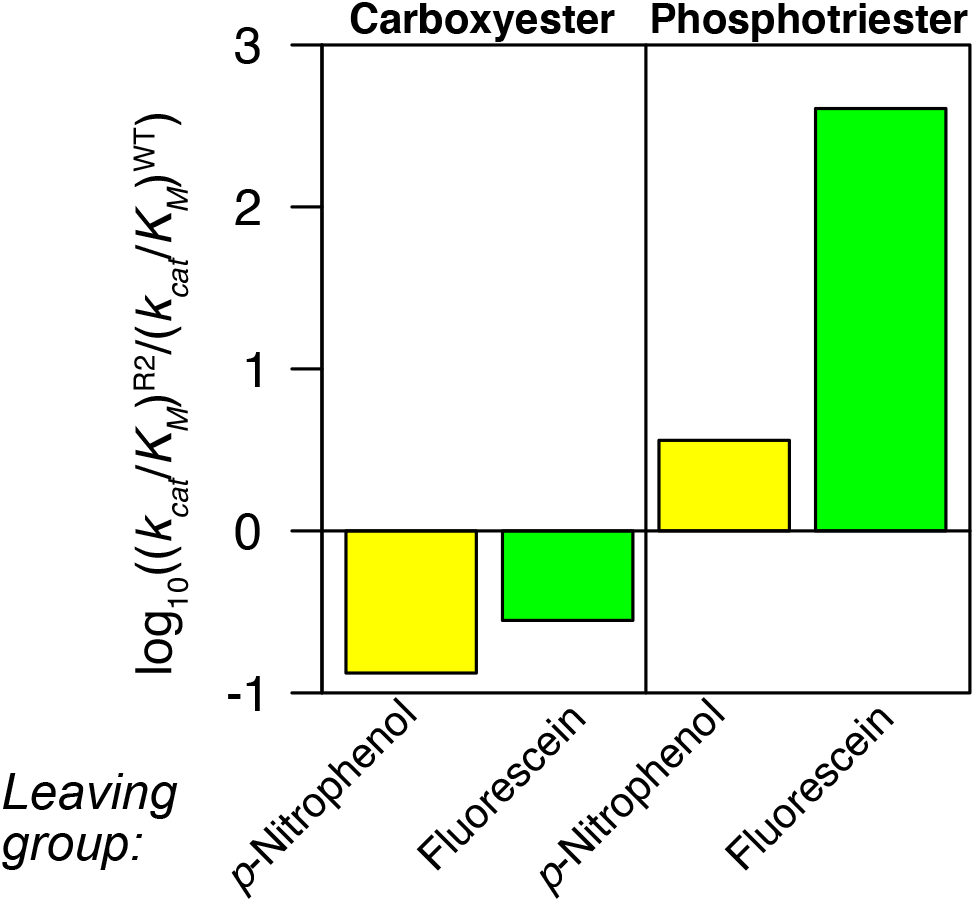
Comparison of reaction type specificity and leaving group preference for P91-WT and P91-R2 indicates that the main difference is intermediate formation. The relative change in catalytic efficiency (*k*_*cat*_/*K*_*M*_) from P91-WT to P91-R2 was measured for carboxyesterase and phosphotriesterase activity with two different leaving groups, *p*-nitrophenol (yellow) and fluorescein (green). Over the course of the directed evolution campaign, P91 specialises for both phosphotriester substrates as well as towards the fluorescein leaving group. The degree of specialisation for phosphotriesterase activity over carboxyesterase activity strongly depends on the identity of the leaving-group, suggesting that the major adaptation must affect the transition state stabilisation during the formation of the intermediate (transition state 1, *k*_*2*_). The relative reduction in carboxyesterase activity is not commensurately leaving group-specific, excluding a simple ground state binding effect.

### Brønsted analysis is consistent with rate-limiting intermediate formation

A linear free-energy relationship (LFER) was constructed to quantify the relationship of leaving group ability (p*K*_*a*_, covering the range from 5.9 to 9.1) on the catalytic parameters *k*_*cat*_ and *k*_*cat*_/*K*_*M*_ for a series of paraoxon derivatives (**Figures S1 and S12 and Tables S4–S6**).

The Brønsted coefficients β_LG_ were almost identical between both Michaelis-Menten parameters, *k*_*cat*_ and *k*_*cat*_/*K*_*M*_, for both enzymes. Any significant rate-determining influence of the intermediate hydrolysis (*k*_*3*_) is expected to be independent of leaving group p*K*_*a*_ and would result in diverging effects on *k*_*cat*_ and *k*_*cat*_/*K*_*M*_ (**equations 4 and 5**), as they represent different elemental processes.^49^ Adding to the observation of monophasic burst kinetics, the identical effects of leaving group p*K*_*a*_ on each parameter are consistent with formation of the covalent intermediate as the rate-limiting step of the reaction.

For *k*_*cat*_/*K*_*M*_ this treatment reports on the transition state in the first irreversible step of the reaction. For P91-WT we observed a linear relationship, with a slope (β_LG_ ≈ −0.95 and −1.1, respectively; **Figure 7a**) that was as steep as the uncatalysed reaction (β_LG_ ≈ −1.0 for spontaneous hydrolysis)^22^ or steeper (0.3 to −0.6 for the hydroxide-catalysed reaction),^22^ indicating a substantial charge accumulation on the leaving group during the transition state. The charge compensation by active site groups has been shown to address the challenge of leaving group departure by developing the enzyme’s ability to offset the charge. This charge compensation in the active site leads to shallower Brønsted plots, as previously observed in the evolution^23^ (and also retrospectively by alanine scanning mutations^24^) in the active site of a sulfatase member of the alkaline phosphatase superfamily. The 400-fold efficiency increase in the evolved variant P91-R2 (with six substrates; 2, 5–10, **Figure S1**) is likewise accompanied by a shallower Brønsted slope (β_LG_ for *k*_*cat*_/*K*_*M*_ ≈ −0.55, for *k*_*cat*_ ≈ 0.68, **Figure 7b**). This difference suggests that the leaving group charge offset is at least partially responsible for the acceleration of intermediate formation in P91-R2. This leaving group stabilisation leads to a smaller effective change in the transition state charge at the leaving group oxygen in the enzyme active site as compared to aqueous solution, as the result of improvements after adaptive evolution.

**Figure 7:**
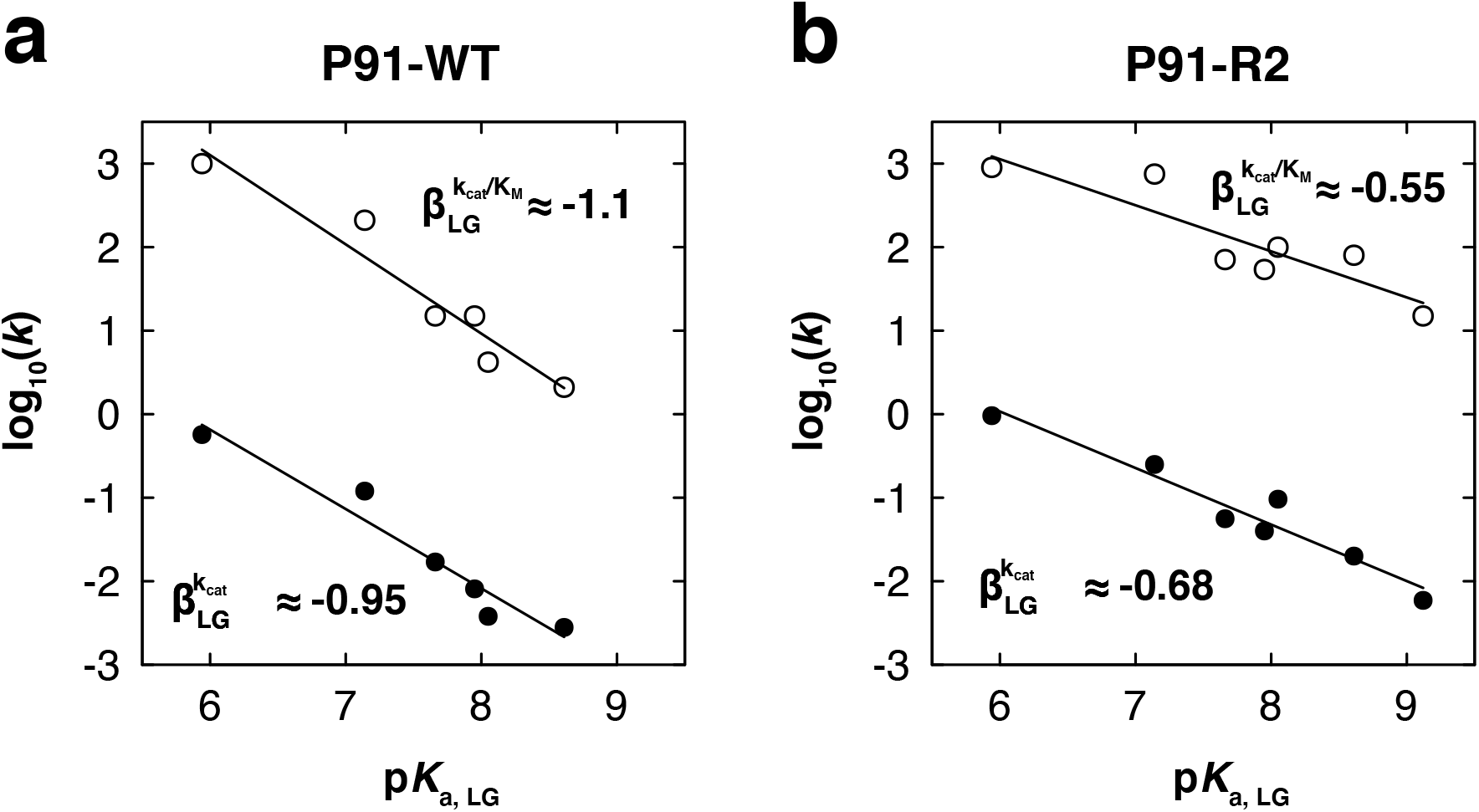
Brønsted analysis shows that the evolved variant P91-R2 accelerates intermediate formation by improved leaving group stabilisation. The Brønsted plots show the linear free-energy relationship between the rate of hydrolysis of paraoxon derivatives and the p*K*_*a*_ of the leaving group for P91-WT and P91-R2. Filled dots: *k*_*cat*_ in s^−1^. Open circles: *k*_*cat*_/*K*_*M*_ in M^−1^s^−1^. **(a)** Brønsted plot for P91-WT: *k*_*cat*_/*K*_*M*_: βLG = −1.07, R^2^ = 0.93; *k*_*cat*_: βLG = −0.95, R^2^ = 0.94. **(b)** Brønsted plot for P91-R2: *k*_*cat*_/*K*_*M*_: βLG = −0.55, R^2^ = 0.80; *k*_*cat*_: βLG = −0.68, R^2^ = 0.94. As the slope of the linear fits (β_LG_) is very similar for both kinetic parameters (*k*_*cat*_ and *k*_*cat*_/*K*_*M*_) intermediate formation (*k*_*2*_) must be the rate-limiting step. The lower β_LG_ of P91-R2 (≈ −0.6) compared to P91-WT (≈ −1) indicates that the evolved variant has adapted to offsetting the charge which accumulates on the leaving group during the transition state.

We note that the metal-catalysed enzymatic reactions in PON1^46^ and *Bd*PTE^47^ were associated with much steeper relationships (β_LG_ ≈ −1.6 and −1.84, respectively), suggesting an entirely different scenario, possibly with a different, larger β_EQ_ (as in non-protein metal complexes^48^) or more nucleophilic involvement (compared to P91’s thiolate) with little charge compensation on the leaving group.

## IMPLICATIONS AND CONCLUSIONS

### Droplet microfluidics allow exploration of new mechanistic terrain

Five mutations were sufficient to increase P91’s phosphotriesterase activity by a factor of ≈ 400, leveraging its catalytic efficiency close to 10^6^ M^−1^s^−1^ and its turnover rate into the range of > 10 s^−1^, thus approaching the catalytic efficiencies of many metal-dependent, ‘conventional’ phosphotriesterases after only two rounds of evolution. This is the first instance of a catalytic triad turning over organophosphates at efficiencies comparable to naturally evolved, metal-dependent enzymes and defines a cysteine triad as an evolvable motif for phosphotriesterase activity. As this catalytic motif has no precedent for this reaction and no database sequences with identical functionality exist, prediction of point mutants based on phylogeny is impossible in this case. In the absence of other cysteine-containing catalytic triads in evolved metal-free phosphotriesterases, no sequence comparisons or mechanistic information were available – a knowledge deficit that could be overcome with the high throughput of our approach. The focussed combinatorial library design combined with the high throughput of microfluidic droplet sorting allowed leaps in sequence space, giving access to and establishing new mechanistic territory that, despite strong selective pressure, has not been exploited in Nature. Indeed, this work has achieved one of the highest single-round improvements in catalytic efficiency obtained by directed evolution in microfluidic droplets (only surpassed by a single droplet evolution campaign of an oxidase^25^), underlining the utility of droplet screening. The precision of control over reaction time obtained by switching between off-chip and on-chip droplet incubation allowed flexible fine-tuning of selection stringency over a large range of catalytic efficiencies. Interestingly, the mutations in variant P91-R2 do not correspond to the individually *best* mutations at the respective positions (i.e. their catalytic contributions are not additive). Thus, the combination of mutations in P91-R2 would not have emerged from iterative saturation mutagenesis of single residues (see Figure S13). The ultrahigh throughput allowed combinations to be screened in one experiment, highlighting how droplet screening enables navigating sequence space without having to take shortcuts that may be at the peril of ignoring epistatic interactions.

### A plausible mechanism for P91 and its improved mutants

Our pre-steady state measurements are consistent with a covalent mechanism of phosphotriester hydrolysis reminiscent of serine proteases, involving formation of an intermediate via the cysteine of the catalytic Cys-His-Asp triad. However, in contrast to serine-triad enzymes, we find that P91 is already pre-disposed for fast de-phosphylation (i.e. breakdown of the covalent intermediate) and rather rate-limited by the initial nucleophilic attack on the substrate. This is consistent with a more labile cysteine-connected thio-phosphate intermediate (compared to a covalent adduct via serine), which leads to formation of a lower energy P-O bond instead by subsequent hydrolysis. In energetic terms, introducing a cysteine as a nucleophile instead of a serine increases the ground state energy of the intermediate thiophosphate and thus lowers the barrier to hydrolysing it. This increased propensity of the P-S bond for hydrolysis (compared to the P-O bond) in phosphate triesters is well established.^26^

The residue changes identified in this directed evolution campaign address the formation of the intermediate and consequently, the evolved variant P91-R2 achieves its higher efficiency over wild-type by accelerating the initial formation of the intermediate (*k*_2_). The residues with the highest individual and combined effects upon mutation are Ile211Trp and Leu214Val, located in a loop that is partly covering the active site entrance. This loop is anchored by a new hydrogen bond to Glu73, a round 2 mutation (from Ala), consistent with a better fit for the promiscuous substrate and possibly allowing π-π-interactions between the aromatic leaving group fluorescein and the new aromatic tryptophan in position 211. For the formation of the covalent adduct (transition state 1, TS1) the cysteine thiol is polarised and deprotonated while the charge developing on the leaving group needs to be stabilised as it departs. During the hydrolysis of the covalent adduct (transition state 2, TS2) a water molecule needs to be positioned, polarised, and deprotonated (**Figure 1**). The active site loop and the catalytic triad’s His199 are interconnected (through polar contacts with Leu214’s backbone amide, which in turn interacts with the backbone of Ile211). These extended interactions may subtly tweak the positioning of His199 and thus influence both the deprotonation of Cys118 (in TS1) and the deprotonation of water (in TS2), better accommodating the geometry of the phosphate transfer.The flattening of the Brønsted slopes (**Figure 7**) is consistent with acceleration of the initial phosphotriester binding by removing a charge incompatibility in the transition state of the phosphorylation step, e.g. at the leaving group. In analogy to the canonical mechanism for ester hydrolysis in the same fold, the oxyanion formed during the transition state is stabilised by backbone nitrogen of Ala38. His199 would be in a position to facilitate leaving group charge offset at the apical position, so re-positioning of His199 could offset leaving group charge development (affecting *k*_*2*_ and β_LG_) in P91-R2.^27^

### An α/β-hydrolase with a cysteine-containing triad is predisposed for promiscuous head start activity and evolvable for phosphotriester hydrolysis

While serine-containing catalytic triads (as in acetylcholinesterase) are irreversibly inhibited by organophosphates, the pre-steady state analysis presented here suggests why a protein with a cysteine-containing catalytic triad was selected in metagenomic screening,^13^ despite the preponderance of serine-containing triads in Nature. The quick and irreversible formation of a stable serine-intermediate would create a single turnover catalyst, so serine enzymes with multiple turnovers may not be easily accessible: the breakdown of this phosphorylated adduct is intrinsically difficult for carboxyesterases as the ground state of the phosphotriester intermediate geometrically resembles the tetrahedral transition state of esters which esterases have evolved to bind tightly. In contrast, a thiolate (as in the intermediate formed with a cysteine enzyme) would be a better leaving group adduct than alkoxide, making it easier to resolve a covalent adduct (Δp*K*_a_ ≈ 13 (Ser) - 8.5 (Cys) ≈ 4.5).^28^ Thus, P91’s pre-disposition for high de-phosphorylation rates can be explained by the intrinsic reactivity of its cysteine triad. In addition to intrinsic chemical promiscuity, substrates need to fit in the active site, requiring binding promiscuity in order to promote the breakdown of a covalent adduct via a trigonal-bipyramidal (instead of a tetrahedral) transition state. However, for a head start activity^29^ leaving group stabilisation would not be required (or at least would not have to be as efficient for a departing sulfur as for oxygen). P91-WT is already pre-disposed with a high de-phosphorylation rate, explaining how it could be found in a naïve library in a phosphotriesterase screen.^13^ Directed evolution found a solution for the remaining catalytic problem, the rate-limiting nucleophilic attack on the substrate.

The difference in p*K*_*a*_ between serine and cysteine in different catalytic triads should also have consequences on the susceptibility for a (fatal) side reaction, hydrolytic de-alkylation of the phosphotriester adduct, dubbed ageing: upon water/hydroxide attack at the phosphorylated intermediate, the transition state collapses with preferential loss of the best leaving group. In the case of a serine triad, one of the two ethoxy side groups of the phosphotriester is prone to leave instead of the serine, resulting in a negatively charged adduct which is resistant to further hydrolysis. As the cysteine thiol group is a much better leaving group than any of the phosphate’s ethoxy side groups, P91 should be much more resistant to ageing than serine triad enzymes.

Prior to this work, metal catalysis was considered the only efficient mechanistic solution for organophosphate triester hydrolysis identified in Nature, having convergently evolved in different protein folds on a very short evolutionary timescale. Based on their postulated origin in lactonases,^30^ it is possible to speculate why they emerged first: the transition state of lactone hydrolysis (an imperfect tetrahedron due to ring strain), is somewhat more similar to a phosphotriester transition state than that of esterases evolved for open chain substrates. It is also possible to argue that a subtle rearrangement of active site metal ion coordination is a simpler solution to accommodating a geometrically different transition state, with minimal constraints compared to an active site amino acid arrangement. Furthermore, the lack of an intermediate in metal hydrolases avoids a potentially rate-limiting additional step.

Nevertheless, this work describes a second line of versatility for adaptive phosphotriesterase evolution in hydrolases with cysteine-containing catalytic triads. While almost all serine-containing catalytic triads (as in acetylcholinesterase) are irreversibly inhibited by organophosphates, there is also evidence that adaptive evolution can activate these enzymes for multiple turnover.

The only known naturally evolved α/β hydrolase that can escape organophosphate inhibition at significant rates is an insect esterase with a Ser-His-Glu catalytic triad that is distantly related to P91 (sequence similarity to P91 ≈ 14 %, structural similarity: RMSD ≈ 4.3 Å). This Gly137Asp mutant of α-carboxylesterase αE7 (*Lc*αE7 Gly137Asp) has evolved very low phosphotriesterase activity in the blow fly *Lucilia cuprina* in response to insecticide exposure (*k*_*cat*_/*K*_*M*_ ≈ 10^2^ M^−1^s^−1^).^31,32^ However, this enzyme is mainly characterised by high affinity to pesticides and a very slow reactivation rate (*k*_*cat*_ ≈ 10^−3^ s^−1^), its dephosphorylation rate being roughly four orders of magnitude lower than of P91-R2.^33^ It is thus fundamentally still a stoichiometric scavenger of pesticides rather than as fast-turnover enzyme. Even wild-type P91 is already 100-fold faster as a phosphotriesterase (*k*_*cat*_ ≈ 0.1 s^−1^).^13^

Similarly, human butyrylcholinesterase, a close homologue of synaptic acetylcholinesterase, has been engineered at the corresponding site (Gly117His) into a slow turnover enzyme, but also shows similar, slow turnover (*k*_*cat*_ ≈ 0.09 s^−1^).^34,35^ A bacterial homologue, *p*-nitrobenzyl esterase from *Bacillus subtilis*, bearing the homologous mutation to BChE Gly117His (*Bs*pNBE Ala107His) showed very slow reactivation and could be further evolved to reach turnover rates of ≈ 1 h^−1^ (≈ 0.0003 s^−1^) after treatment with paraoxon.^36^ A triple mutant of a snake acetylcholinesterase (from *Bungarus fasciatus*) has been reported to display slow promiscuous organophosphate hydrolysis, albeit with very low catalytic efficiencies in the range of 10^−1^ to 10^1^ M^−1^s^−1^, depending on the substrate.^37^

In each case the slow de-phosphorylation rate of all those serine triad hydrolases (≈ 10^−4^ to 10^−2^ s^−1^) is in good agreement with the rates observed for the nucleophile-exchanged P91 variants (which bear a serine instead of the cysteine). The turnover rate of the evolved P91-R2 variant (≈ 15 s^−1^) is two to four orders of magnitude higher than of the serine enzymes. Conversely, the rate of intermediate formation *k*_*2*_ is reported to be 1.3 s^−1^ for the serine enzyme *Lc*αE7 mutant^33^ and, inferring from *k*_*cat*_, about two orders of magnitude lower in the cysteine enzyme P91-WT. The limited catalysis by these serine enzymes stands in stark contrast to the ready evolvability of P91 and the activity of the improved mutant P91-R2 observed in this work.

This finding also raises the question whether acetylcholinesterase and its homologues would become proficient phosphotriesterases when exchanging their catalytic serine for a cysteine. Nucleophile exchange in a triad is usually deleterious to the native activity but has been shown to be rescuable by directed evolution.^38,39^ Given that such nucleophile exchanges have a low probability in natural non-targeted randomisation events, and the known catalytically phosphotriesterase-activating mutations (reviewed above) have modest effects, it can be explained that metal-dependent hydrolases won out in the evolutionary experiment played out over the last decades, in response to phosphotriester contamination.

The untapped catalytic versatility of ecological consortia is underlined by empirical identification of P91 with its alternative active site functionality in metagenomic libraries.^13^ Using ultrahigh-throughput methods will be crucial not only to explore catalytic solutions that are dominant amongst extant enzymes, but also to uncover additional reactivities that form second lines of evolutionary contingency, even against anthropogenic chemical compounds that environments have never ‘seen’. Here we show that this contingency is not limited to a transient, low efficiency catalyst, but that initial catalytic solutions are ‘evolvable’, at least when large, smart libraries in microfluidic droplets with ultrahigh throughput are used. The evolutionary improvements demonstrated here make them as efficient as the previously known metal-containing phosphotriesterases, suggesting that not only accidental low-efficieny promiscuous catalyst exist, but instead alternative enzymatic strategies are available in a given metagenome, with the potential to be truly proficient and ready to be recruited for a survival advantage or for their biocatalytic application. Natural repertoires thus hold a variety of solutions for catalytic challenges: even if not explored thus far, they are available to contribute new-to-nature reactions^42^ or mechanisms.

## METHODS

### Cloning and library construction

Libraries of the *p91* gene were constructed on the high-copy number (≈ 800 copies/cell), anhydrotetracyclin-inducible plasmid pASK-IBA5plus (IBA Life Sciences, Germany) bearing an N-terminal StrepII-tag. Single-site saturation libraries were constructed by Golden Gate Assembly^40^ with partly degenerate primers according to the ‘22-codon trick’.^41^ Multiple site saturation libraries were constructed by assembly of gene fragments into the full-length gene by assembly PCR (library P91-A) or Golden Gate Assembly (library P91-B). The fragments were created by PCR with primers bearing the degenerate codons NNK (library P91-A) or NDT/VHG/TGG (library P91-B). Further details on cloning and library construction can be found in the Supplementary Information.

### Library screening in microfluidic droplets

Monodisperse water-in-oil microdroplets were generated with a microfluidic flow-focusing device (**Figure S3a**). Fluorocarbon oil (Novec HFE-7500, 3M, USA) containing 0.5 % (w/w) surfactant (008-FluoroSurfactant; RAN Biotechnologies, USA) served as oil phase. The two aqueous streams were supplied with the cell solution and with a 3 μM substrate solution containing lysis agents (0.7× BugBuster protein extraction reagent, Merck Millipore; 60 kU/mL rLysozyme, Novagen) in droplet assay buffer, respectively. For long incubation times in evolution round 1, requiring off-chip incubation, droplets were collected into a long polyethylene tubing (0.38 mm ID, 1.09 mm OD; Portex Smiths Medical, USA) which was closed with a syringe needle after collection. For short incubation times in evolution round 2, requiring on-chip incubation, an integrated chip was used, combining a flow-focussing module, a delay line, and a sorting module on a single device (**Figure S4**). On the sorting chip (round 1) or the sorting module of the integrated chip (round 2), droplets were sorted according to their green fluorescence (excitation wavelength: 488 nm) at a rate of ≈ 300–1000 Hz. Plasmids from sorted droplets were recovered by de-emulsification with 1H,1H,2H,2H-perfluorooctanol (Alfa Aesar, USA) and subsequent column purification and electroporation into highly electrocompetent *E. coli* cells (E. cloni 10G Elite, Lucigen, USA). Further details on the microfluidic screening can be found in the Supplementary Information.

### Kinetic measurements and data analysis

#### Steady-state kinetics

For steady-state kinetic measurements, His6-tagged P91 variants (expressed and purified as detailed in the Supplementary Information) were used. The progress of the reaction was monitored by absorbance or fluorescence in a spectrophotometric microplate reader (Tecan Infinite 200PRO, Tecan, Switzerland) at 25 °C. In order to determine the Michaelis-Menten parameters *k*_*cat*_, *K*_*M*_, and (in case of substrate inhibition) *K*_*i*_, the initial rates were fitted to the Michaelis-Menten equation:

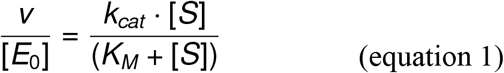

or, in the case of substrate inhibition,

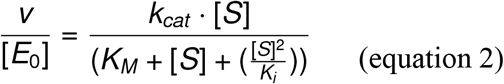

where v is the initial rate of the reaction, [E_0_] is the initial enzyme concentration, and [S] is the substrate concentration.

#### Pre-steady state kinetics

Fast transient-state kinetics for the hydrolysis of phosphotriesters FDDEP and paraoxon-ethyl were measured with a SX20 stopped-flow spectrophotometer (Applied Photophysics, UK) at the same temperature, in the same buffer and with the same substrate concentrations as the steady-state kinetics. Measurement traces were fitted to the following exponential burst equation:

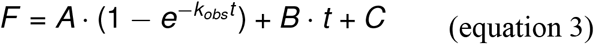

where *F* is the measured absorbance or fluorescence, *t* is the time, *A* is the amplitude of the burst, *B* is the slope of the second phase of the reaction and *C* is the offset.

The observed rate *k*_*obs*_ was then fitted to the following equation to determine *k*_*2*_ (**Figure S9, Table S3**).

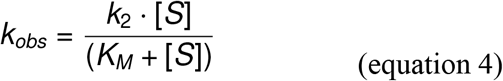

The de-phosphorylation rate *k*_*3*_ was separately determined with the mono-substituted substrate paraoxon-ethyl to exclude potential obfuscating effects due to the complex downstream kinetics of the double substituted substrate FDDEP (where the original double-substituted substrate FDDEP and the initial reaction product, the mono-substituted FMDEP, compete for turnover). As reaction with paraoxon-ethyl forms the same diethyl-phosphate covalent intermediate, *k*_*3*_ with FDDEP is identical to *k*_*3*_ with paraoxon-ethyl. Assuming a reaction model with one reversible binding step and two irreversible steps (**Figure 1**), the turnover number *k*_*ca*t_ and the catalytic efficiency *k*_*cat*_/*K*_*M*_ can be described as:

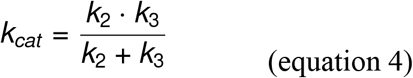

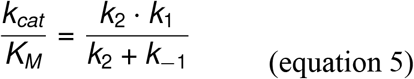

Given that in this case *k*_*2*_ ≫ *k*_*3*_, the term for *k*_*cat*_ can be simplified to:

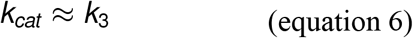

Thus, the initial slope of the second phase of the burst reaction with paraoxon-ethyl was plotted against substrate concentration and fitted to the Michaelis-Menten equation to determine *k*_*3*_ (**Figure S10, Table S3**). In cases where the burst is less pronounced (as in P91-R1 Cys118Ser) the *k*_*cat*_ is a lower estimate of *k*_*3*_ while the real value of *k3* might be higher. Further details on kinetic measurements can be found in the Supplementary Information.

## Supporting information

Microfluidic chip designs

## ASSOCIATED CONTENT

Additional experimental procedures (chip design, microfluidic device operation, droplet sorting, DNA recovery, microtiter plate screening, protein purification, substrate synthesis, NMR data, gene sequences), figures (substrate structures, mutational scanning, kinetic data, NMR spectra), and tables (kinetic parameters, comparison of phosphotriesterase literature values, primer sequences) as well as microfluidic chip designs are available in the Supplementary Information (SI).

## Notes

The authors declare no competing interests.

