## Supplementary Information for "Ultrahigh-throughput directed evolution of a metal-free α/β-hydrolase with a Cys-His-Asp triad into an efficient phosphotriesterase"

David Schnettler Fernández, Oskar James Klein, Tomasz S. Kaminski, Pierre-Yves Colin, Florian Hollfelder*

Department of Biochemistry, University of Cambridge, 80 Tennis Court Road, Cambridge, CB2 1GA, United Kingdom

### 1. Supplementary Methods

#### Materials

All chemicals were purchased from Sigma-Aldrich and all biological reagents from New England Biolabs, unless otherwise stated. The fluorogenic model substrate fluorescein di(diethylphosphate) (FDDEP, 1) was synthesized as previously described.[1] The phosphotriesters 5–10 for linear free energy relationship measurements were synthesised as detailed in the Supplementary Methods.

#### Cloning and library construction

***Single-site saturation libraries*** were constructed using the 22-codon trick:[2] primers bearing the degenerate codons NDT, VHG and TGG were mixed in the ratio 12:9:1, in order to achieve balanced amino acid representation while avoiding stop codons, allowing to sufficiently oversample the diversity of a single randomised position with a single 96-well microtiter plate. The gene was amplified from the randomised position in a whole-plasmid PCR with Q5 DNA Polymerase and digested and re-circularised in a single step of Golden Gate Assembly using BsaI-HFv2 and T4 DNA ligase.[3] Single-site mutagenesis was carried out according to the same principle.

***Multiple-site saturation library (P91-A, round 1):*** To construct multiple site-saturation libraries, the P91 gene was sub-divided into suitable fragments such that each randomised position was covered by a primer with the degenerate codon NNK. The fragments were created individually by PCR and then assembled by a further assembly PCR. Forward primers for fragment creation contained degenerate codons and ≈ 20 base pair homology arms to the neighbouring fragments with an annealing temperature of 55 ˚C. For assembly PCR, the individual fragment PCR products were pooled and amplified with outer primers by PCR with Phusion DNA Polymerase at the annealing temperature of the homology arms. The resulting PCR product was column-purified (DNA Clean & Concentrator-5 Kit; Zymo Research) and digested with NheI and HindIII. After gel purification, the library insert was ligated into accordingly digested pASK-IBA5plus vector backbone with T4 DNA Ligase at 16 ˚C for 14 h. For plasmid amplification, the column-purified ligation product was electroporated into highly electrocompetent *E. coli* (E. cloni 10G Elite, Lucigen). After overnight incubation, the bacterial carpet was scratched off the agar plate with 6 ml of LB medium and the DNA was extracted with a plasmid isolation kit (GeneJET Plasmid Miniprep Kit; Thermo Fisher). The quality of the library was assessed by measurement of the peak heights in the sequencing chromatogram of a pooled sample as previously described.[4]

***Multiple-site saturation library (P91-B, round 2):*** For the second round, the library was created by whole-plasmid PCR with degenerate primers containing a type-II restriction site overhang (BsaI: GGTCTCN) to create sticky ends for ligation. At the desired position, forward primers contained the mutagenic trinucleotides NDT, VHG and TGG and were used as a 12:9:1 mixture, previously described as the '22-codon trick'.[2] After whole-plasmid PCR with Phusion DNA Polymerase (ThermoFisher) or Q5 DNA Polymerase the PCR mix was digested with DpnI to selectively remove wild-type template. Subsequently, the amplicons were column-purified (DNA Clean & Concentrator-5 Kit; Zymo Research) and digested with BsaI to create sticky ends. After a further column purification, amplicons were self-ligated with T4 DNA Ligase. The ligation product was amplified and purified in the same way as the round 1 library.

##### Chip design and preparation of microfluidic devices

The channel layout for the microfluidic chips was designed using AutoCAD (Autodesk, USA) and printed out on a high resolution film photomask (Micro Lithography Services, UK). The designs are shown in **Figures S3 and S4** and are deposited as DXF files on DropBase (https://openwetware.org/wiki/DropBase:Devices):

https://openwetware.org/wiki/Dropbase:_3-pL-droplets-01
https://openwetware.org/wiki/DropBase:droplet_electrosorting_3
https://openwetware.org/wiki/DropBase:droplet_electrosorting_4
https://openwetware.org/wiki/DropBase:droplet_electrosorting_5

##### Photolithographic fabrication of wafer master moulds for microfluidic devices

The microfluidic devices were fabricated following standard photolithography and soft lithography protocols.[5] In brief, a silicon wafer (Prime Grade, 3 inch diameter, Czochralski Silicon (CZ-Si) wafer, thickness = 381 ± 20 µm, one-side polished; purchased from Microchemicals, Germany) was covered with a thin layer of SU-8 2000 series photoresist material using a spin coater (SPIN150i spin coater, Polos by SPS, Germany). Depending on a channel height, SU-8 2010 or SU-8 2025 photoresists were used (Kayaku Advanced Materials, Japan). Next, the wafer was soft-baked on a hot plate and the channel pattern was subsequently patterned into the master by photolithography using a mask aligner (MJB4 mask aligner; Süss MicroTec, Germany). In subsequent steps, the wafer was post-baked and, in the case of two-layer devices, the next layer of SU-8 resist was spincoated, followed by a second round of soft-baking, exposure and post-baking. After the post-baking step, the single- or double-layer chip, was developed in propylene glycol monomethyl ether acetate (Sigma Aldrich). Next, the wafer was hard-baked for 10 min at 200 ˚C and the hights of the structures were measured using a profilometer (Veeco Dektak 6M Stylus Surface Profilometer; Bruker, USA). Finally, the chip was silanised by deposition of pure trichloro(1H,1H,2H,2H- perfluorooctyl)silane (2 µL) in close proximity to the wafer placed in a Petri dish which was kept for 30 min in the vacuum chamber at 20 mbar to generate vapours of silane. A detailed overview of the steps of the protocol for the different devices is shown in **Table S0**.

**Table S0.** Protocols of photolithography of master mould of microfluidic devices used in this study.

| | **Device type** | | | | |
|---|---|---|---|---|---|
| | Flow focusing droplet generation device | FADS device | FF devices with delay lines and FADS module | | |
| | | | Layer A | Layer B1 (20 loops device) | Layer B2 (5 loops device) |
| Nominal thickness (in µm) | 12 | 20 | 15 | 30 | 15 |
| Photoresist used | SU8-2010 | SU8-2025 | SU8-2010 | SU8-2025 | SU8-2010 |
| Spin coating speed | 1st step: 10 s, 500 rpm<br>2nd step: 30 s, 2000 rpm | 1st step: 10 s, 500 rpm<br>2nd step: 30 s, 4000 rpm | 1st step: 10 s, 500 rpm<br>2nd step: 30 s, 1600 rpm | 1st step: 10 s, 500 rpm<br>2nd step: 30 s, 3000 rpm | 1st step: 10 s, 500 rpm<br>2nd step: 30 s, 3400 rpm |
| Soft baking | 3 min at 95 ºC | 1 min at 65 ºC<br>5 min at 95 ºC | 3 min at 95 ºC | 1 min at 65 ºC<br>5 min at 95 ºC | 3 min at 95 ºC |
| Exposure (at ~10 mW $cm^{-2}$) | 2 x 6 s, 2 s waiting time | 2 x 7 s, 2 s waiting time | 2 x 7 sec, 2 s waiting time | 2 x 7.5 s, 2 s waiting time | 2 x 7 s, 2 s waiting time |
| Post baking | 3 min at 95 ºC | 1 min at 65 ºC<br>5 min at 95 ºC | 3 min at 95 ºC | 1 min at 65 ºC<br>5 min at 95 ºC | 3 min at 95 ºC |
| Development in the beaker filled with 30-50ml of PGMEA (Propylene glycol methyl ether acetate, Sigma Aldrich) | Approx. 5 min until all uncured SU-8 is removed from the wafer.<br>The development time depends on the intensity of manual agitation (for 2-layer chips, implemented only after deposition of the layer B) | | | | |
| Hard baking | 10 min at 200 ºC<br>(for 2-layer chips a hard baking step was applied only after development of the final device) | | | | |
| Measured range of thicknesses (in µm) | 11.8–11.9 | 21.5–23 | 14.7–15.0 | *Total thickness of layer A and B1:*<br>46–48 | *Total thickness of layer A and B2:*<br>28–29 |

#### Soft lithography protocol for preparation of PDMS chips

For the fabrication of microfluidic PDMS chips, the master was covered with a mixture of poly(dimethyl)siloxane (PDMS) and curing agent (Sylgard 184 Silicone Elastomer Kit, Dow Chemical Company, USA) in a 10:1 ratio (w/w). After degassing and curing at 65 ˚C for approximately 3 h, the PDMS device was removed from the master and holes for tubing connections were punched using a 1 mm biopsy punch with a plunger (Kai Medical, Japan). The device was then attached to a 1 mm thick microscope glass slide (flow-focusing device) or a 0.13 mm thin glass cover slip (sorting and delay line devices) by treatment with oxygen plasma (Femto plasma system; Diener Electronic, Germany) for 30 s, followed by a baking step of ≈ 20 min incubation at 65 ˚C. For hydrophobic treatment of the channel surface, a freshly prepared and filtered solution of 1 % (v/v) trichloro(1H,1H,2H,2H-perfluorooctyl)silane in fluorinated oil (Novec HFE-7500, 3M, USA) was injected into the channels, followed by approximately 45 min of incubation on a hot plate at 65 ˚C. For small devices, the silane-containing oil was slowly injected by manual operation of a syringe. In contrast, for devices with a delay line, manual injection could lead to damage on the chip (due to high resistance and build-up of back-pressure) and therefore a syringe pump was used at at rate of 200 μL/h to inject the silane-containing oil.

**Preparation of cells for compartmentalisation**

*Escherichia coli* cells (E. cloni 10G Elite; Lucigen, USA) were transformed with 2.5 μL of library plasmids, yielding $10^6$–$10^7$ colonies after overnight incubation on agar plates, as determined by serial dilution. Transformed cells were induced with anhydrotetracycline (final concentration 200 ng/mL; IBA Life Sciences, Germany) and incubated for expression for 14 h in 20 mL LB medium at 20 ˚C and 220 rpm shaking. After expression, the cells were washed five times with buffer (100 mM MOPS-NaOH, 150 mM NaCl, pH 8.0) and diluted to $OD_{600}$ = 1.0. A 200 μL aliquot of the suspension was diluted 1:2 with 100 μL droplet assay buffer (100 mM MOPS-NaOH, 150 mM NaCl, pH 8.0, cOmplete EDTA-free protease inhibitor (one tablet per 50 mL; Roche, Switzerland) and 100 μL Percoll (a silica nanoparticle solution to reduce cell-cell adhesion and prevent sedimentation of cells in the syringe; Cytiva, USA) to a bacterial density of $OD_{600}$ = 0.5. This bacterial suspension was diluted in order to match the desired final average bacterial droplet occupancy (which determines the specific occupancy of individual droplets according to the Poisson distribution). For instance, in a droplet volume of ≈ 3 pL a final bacterial density of $OD_{600}$ = 0.25 resulted in 16 % of droplets with a single bacterium and 3 % of droplets containing two or more bacteria, as determined by microscopic imaging. All solutions (except for cell suspensions) were previously filtered with 0.2 μm PTFE syringe filters (Acrodisc CR 13 mm syringe filters, PALL Life Sciences, USA) to avoid clogging of microfluidic channels.

**Compartmentalisation of cells into microdroplets**

Monodisperse water-in-oil microdroplets were generated with a microfluidic flow-focusing device (design and fabrication, of microfluidic devices are described in the Supplementary Information). The device was connected via PE tubing (0.38 mm inner diameter, 1.09 mm outer diameter; Portex Smiths Medical, USA) to glass syringes (100 μL and 1 mL; SGE Analytics, Australia), which were driven by syringe pumps (neMESYS, Cetoni, Germany). Fluorocarbon oil (Novec HFE-7500, 3M, USA) containing 0.5 % (w/w) surfactant (008-FluoroSurfactant; RAN Biotechnologies, USA) served as oil phase. The two aqueous streams were supplied with the cell suspension and with a 3 μM substrate solution containing lysis agents (0.7× BugBuster protein extraction reagent, Merck Millipore; 60 kU/mL rLysozyme, Novagen) in droplet assay buffer, respectively. The enzymatic reaction was initiated by cell lysis upon droplet formation from the three supply streams. Droplet formation was monitored using an inverted microscope (SP98I, Brunel Microscopes, UK) with a high-speed camera (Phantom Miro eX4, Vision Research, USA). For long incubation times in evolution round 1, requiring off-chip incubation, flow rates of 50 μL/h for the aqueous phases and 500 μL/h for the oil phase were used to generate droplets with a volume of 3 pL at rates of 0.5–3 kHz. Droplets were collected into a long PE tubing (0.38 mm ID, 1.09 mm OD; Portex Smiths Medical, USA) which was closed with a syringe needle after collection. For short incubation times in evolution round 2, requiring on-chip incubation, an integrated chip was used, combining a flow-focussing module, a delay line, and a sorting module on a single device (**Figure S4**). For tight spacing of the droplets in the delay line, required for even mixing and homogenous incubation times, oil was removed through an oil extractor. At the end of the delay line, droplets were injected into the sorting module. On this chip, flow rates were 7.5 µL/h for the aqueous phases, 25 µL/h for the oil phase, ≈ -10 µL/h for the oil extractor, and ≈ 300 µL/h for the spacing oil, resulting in a droplet volume of ≈ 11 pL.

##### Fluorescence-assisted droplet sorting (FADS)

Optics and electronics of the microfluidic on-chip sorting device were set-up as previously described[6,7]. After incubation at room temperature, droplets were reinjected from the collection tubing into the sorting device at a rate of 10–25 µL/h. To enable precise sorting of single droplets, the distance between the droplets was increased by injection of spacing oil (Novec HFE-7500, 3M, USA) into the device at a flow rate of 100–300 µL/h. The asymmetric Y-shaped junction in the device ensures that all droplets automatically flow into the waste channel, unless deviated by an electrical pulse into the sorting channel. A 488-nm laser was focused 100 µm upstream of the sorting junction through a 40× microscope objective (UPlanFLN, Olympus, Japan) for fluorophore excitation and the emitted fluorescent light was collected and amplified using photomultiplier tubes (PMM02, Thorlabs, USA). Whenever the fluorescence peak reached a user-defined threshold, an electric field was applied by the two electrodes on the sorting device, attracting the highly fluorescent droplet towards the narrower sorting channel (**Figure 1**). Droplets were sorted into a collection tube pre-filled with 100 µL nuclease-free water.

##### DNA recovery from microdroplets

Plasmids from sorted droplets were recovered by de-emulsification with 1H,1H,2H,2H-perfluorooctanol (Alfa Aesar, USA) and subsequent column purification and electroporation into highly electrocompetent E. coli cells (E. cloni 10G Elite, Lucigen, USA) in a modification of the previously described protocol.[6] In brief, after droplet sorting, 400 ng of salmon sperm DNA (Invitrogen, USA) were added to the collection tube. Subsequently, PFO (approx. half the volume of the collected oil volume; e.g. 400 µL PFO for 800 µL oil in the collection tube) was added. The tube was vortexed for ≈ 60 s and centrifuged for 1 min at 14000 g. The top aqueous layer was completely removed and transferred into a new tube. The oil phase remaining in the collection tube was re-extracted by adding 100 µL nuclease-free water, 400 ng salmon sperm DNA, 100 µL PFO, vortexing for 1 min and subsequent centrifugation for 1 min at 14000 g. The top aqueous layer was completely removed and united with the previously removed aqueous layer. The united aqueous fractions usually contain a small remaining bottom oil phase. This remaining oil phase was then extracted by addition of 100 µL PFO, 1 min of vortexing, and 1 min of centrifugation at 1400 g. The top aqueous phase was then removed without any traces of oil phase and the plasmid DNA was column-purified using a DNA Clean & Concentrator Kit (Zymo Research, USA) according to the manufacturer's instructions, with the following modifications: The ratio of binding buffer to sample volume was 6:1. The wash buffer was carefully rinsed along the walls. After the second wash step, the flow-through was removed and the column was centrifuged once more for 1 min to dry. For elution, 6 µL of pre-warmed (50 ˚C) elution buffer were used. The eluted DNA was then electroporated into highly electrocompetent E. coli cells (E. cloni 10G Elite, Lucigen, USA) according to the manufacturer's instructions. Throughout the DNA extraction procedure, low-DNA-binding tubes (1.5 mL DNA LoBind tube; Eppendorf, Germany) and low-retention tips (Axygen Maxymum Recovery Filter Tips; Corning, USA) were used.

#### Microtiter plate screening

To quantify the lysate activity of P91 variants, individual colonies were picked and grown in 96-deep-well plates in 500 µL Luria-Bertani (LB) medium with 100 µg/mL carbenicillin at 37 ˚C/1050 rpm for ≈ 14 h. Subsequently, 10 µL of overnight cultures were used to inoculate 490 µL of medium for expression cultures in 96-well deep-well plates which were grown at 37 ˚C/1050 rpm for ≈ 2 h until $OD_{600}$ ≈ 0.5. Expression was then induced with anhydrotetracycline (final concentration 200 ng/mL; IBA Life Sciences, Germany) and carried out for 14 h at 20 ˚C and 1050 rpm shaking. Cells were pelleted by centrifugation at 3320 g for 60 min, the supernatant was then discarded, and cells were lysed with 100 µL lysis buffer (50 mM HEPES-NaOH, 150 mM NaCl, pH 8.0, 60 kU/mL rLysozyme, 1X BugBuster) for 20 min at 20 ˚C and 1050 rpm shaking. Cell lysates were diluted 1:20 or 1:400 in assay buffer (50 mM HEPES-NaOH, 150 mM NaCl, pH 8.0). For the reaction, 190 µL of the phosphotriester substrate paraoxon-ethyl (100 µM) or FDDEP in assay buffer (3 µM) were added to 10 µL aliquots of the diluted lysate in microtiter plates and the formation of fluorescein or *p*-nitrophenol was recorded in a plate reader (Infinite M200, Tecan, Switzerland) for 15 min at a wavelength of 405 nm for *p*-nitrophenol and at an excitation wavelength of 480 nm and an emission wavelength of 520 nm for fluorescein.

#### Protein expression and purification

Plasmids isolated from single colonies were used to transform BL21(DE3) cells. Expression cultures were then inoculated by a similar 'plating' method as previously described[8,9]. In brief, a dense lawn of freshly transformed BL21(DE3) cells was directly scraped into 500 mL TB medium containing 100 µg/mL carbenicillin. The cells were grown for ≈ 60 min at 37 ˚C/200 rpm before being induced with anhydrotetracyclin (final concentration 200 ng/mL). Protein was then expressed at 20 ˚C/200 rpm for 18–20 h. Cells were harvested by centrifugation at 4000 rcf for 10 min, the supernatant was discarded, and the dry pellet was stored at -80 ˚C. Note that the P91 construct for library screening originally contained an N-terminal StrepII-tag. For high-yield purification (as required for transient-state kinetics), the StrepII-tag was exchanged for an N-terminal 6xHis-tag. For purification, the pellet was resuspended in lysis buffer (50 mM HEPES-NaOH, 150 mM NaCl, pH 8.0, 1 mM TCEP, 20 mM imidazole, 0.5–1 mg/mL lysozyme, 0.1 % Triton X-100, 0.01 % (= 25 units/mL) benzonase nuclease) and rolled for 30–60 min at room temperature. Afterwards, the lysate was cleared by centrifugation at 20000 rcf/4 ˚C for 20 min and the soluble fraction was loaded onto a Ni-NTA gravity flow column (Super Ni-NTA Agarose Resin, Neo Biotech, France). The protein on the column was washed with 5 x 2.5 column volumes of wash buffer (50 mM HEPES-NaOH, 150 mM NaCl, pH 8.0, 1 mM TCEP, 20 mM imidazole) and eluted with 5 x 0.5 column volumes of elution buffer (50 mM HEPES-NaOH, 150 mM NaCl, pH 8.0, 1 mM TCEP, 250 mM imidazole). Subsequently, the eluate was concentrated with a spin concentrator (Vivaspin 10 000 kDa MWCO, Sartorius, Germany) and subsequently exchanged into imidazole-free assay buffer (50 mM HEPES-NaOH, 150 mM NaCl, pH 8.0, 1 mM TCEP) using PD 10 desalting columns (Cytiva, USA). The typical yield was ≈ 200 mg enzyme per litre of culture. Enzyme purity was controlled by SDS-PAGE and concentrations were determined by measurement of absorption at 280 nm on a NanoDrop 2000 Spectrophotometer (Thermo Fisher Scientific, USA), using an extinction coefficient calculated with the ProtParam web tool[10] (https://web.expasy.org/protparam).

#### Kinetic measurements

For steady-state kinetic measurements, 6xHis-tagged P91 variants were used and enzyme concentrations were kept at least 10–100-fold lower than the lowest substrate concentration. Substrate concentrations were chosen to span ≈ 10-fold below and above $K_M$, as far as not limited by substrate solubility. Optimal starting enzyme concentration $E_0$ and substrate concentration ranges were determined for each variant and substrate combination by empirical sampling. Substrates were pre-dissolved in DMSO at stocks of 200-fold the final concentration, in order to ensure constant DMSO concentration (0.5 %) across all substrate concentrations, as DMSO content was found to influence the catalytic parameters while being required as a co-solvent for substrate stocks. Upon measurement, aliquots of these substrate stocks were diluted 1:100 in assay buffer (50 mM HEPES-NaOH, 150 mM NaCl, pH 8.0, 1 mM TCEP), of which 100 μL were subsequently mixed with 100 μL of 2-fold concentrated enzyme solution in microtiter plate wells. The progress of the reaction was monitored by absorbance (for wavelengths and extinction coefficients for each substrate leaving group, see Table S4) or fluorescence (at an excitation wavelength of 480 nm and an emission wavelength of 520 nm for fluorescein) in a spectrophotometric microplate reader (Tecan Infinite 200PRO, Tecan, Switzerland) at 25 ˚C. Absorbance measurements at wavelengths < 320 nm were taken using a quartz 96-well plate. Absorbance maxima and extinction coefficients were determined for the leaving group of each substrate by absorbance wavelength scan followed by a calibration curve (see **Table S4**).The initial rates were extracted by linear fit of the first measurements (at < 10 % progress of the reaction) for each substrate concentration and normalised with an extinction coefficient determined from a calibration curve. Fitting of the data was done with R using the non-linear fitting function `nls()`.[11]

#### Note regarding the determination of $k_2$ with the substrate FDDEP

For the double-substituted phosphotriester FDDEP, the apparent $k_2$ as measured from the burst phase represents the hydrolysis of one of the two phosphotriester groups on the substrate and thus the release of the mono-phosphorylated reaction product, fluorescein mono(diethylphosphate), which has an unknown extinction coefficient. However, the observed burst amplitude (0.4–1.0, thus corresponding to the value expected for fluorescein release) indicates that most of the substrate fluorescence is generated by the release of this first group, thus allowing the use of a fluorescein calibration curve for the approximate quantification of $k_2$.

#### Note on the presence of a covalent intermediate

In principle, the complete absence of an intermediate in the mechanism of P91 is also a possibility. However, the following arguments support the presence of a covalent intermediate:

1. In contrast to other known phosphotriesterases, which have a metal-assisted instead of a nucleophilic mechanism , P91 has no metal in the active site.[12]
2. Dienelactone hydrolase, the closest functionally characterised homologue of P91, shares the same active site triad (Cys-His-Asp) with P91 and forms a covalent intermediate via its cysteine nucleophile in the hydrolysis of esters and lactones.[13]
3. The phosphotriesterase and esterase activities of P91 compete for the same active site, suggesting that they use the same mechanism.[12]
4. Mutation of the active-site cysteine Cys118 to alanine abolishes both, esterase and phosphotriesterase activity in P91.[12]
5. Mutation of the active-site cysteine Cys118 to serine induces burst kinetics in P91 which are simplest explained by the presence of a covalent intermediate.

#### Structural modelling with AlphaFold2

The structure of P91-R2 was modelled using AlphaFold2[14] via the ColabFold implementation[15] (https://colab.research.google.com/github/deepmind/alphafold/blob/main/notebooks/AlphaFold.ipynb).

#### Synthesis of phosphotriesters for linear free energy relationship measurements.

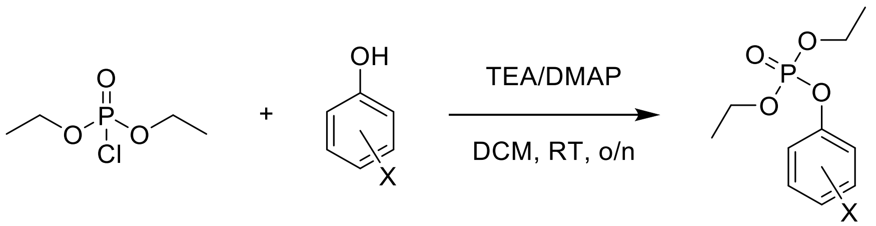

Synthesis procedures and p$K_a$ values were adapted from Khersonsky & Tawfik 2005.[16] In brief, the respective phenol derivative (1 g) and diethyl phosphochloridate (1.2 equivalents) were dissolved in dichloromethane. Base (1 mol eq., triethylamine for all but 3-fluoro-4-nitrophenyl, where dimethylaminopyridine (DMAP) was used) was added dropwise and the reaction mixture stirred at room temperature (≈ 23 ˚C) overnight (≈ 16 h). Reaction progress was followed by thin layer chromatography (silica gel 60, EtOAc:Hex, 2:1). The reaction mixture was washed with HCl (100 mM, 50 mL), saturated $Na_2HCO_3$ (50 mL) and NaCl brine (50 mL, pH ≈ 7). Products contaminated by starting material were purified by flash chromatography (silica gel 60, EtOAc:Hex 2:1). Products were characterized by $^1$H-NMR. NMR spectra can be found in section 5 of the Supplementary Information. All NMR data were collected at 298 K using Bruker Avance spectrometers with $^1$H resonance frequencies of 400 MHz. Chemical shifts (δH) are reported in parts per million (ppm), to the nearest 0.01 ppm and are referenced to the residual non-deuterated solvent peak. Coupling constants (*J*) are reported in Hertz (Hz) to the nearest 0.1 Hz. Data are reported in the order: (i) chemical shift, (ii) multiplicity (s = singlet; d = doublet; t = triplet; q = quartet; m = multiplet; or as a combination of these, e.g. dd, dt etc.), (iii) coupling constant(s) and (iv) integration. Peak integrals were used to produce correction factors for residual solvent contamination.

**3-fluoro-4-nitrophenyl diethyl phosphate (5)**

**$^1$H NMR** (399.6 MHz, $CDCl_3$): δ (ppm) 8.14 (t, $^3J_{HH}$/$^4J_{HF}$ = 8.8 Hz, 1H), 7.24 (dd, $^3J_{HF}$ = 11.5, $^4J_{HH}$ = 2.4 Hz, 1H), 7.20 (m, 1H), 4.29 (m, 4H), 1.41 (td, $^3J_{HH}$ = 7.1 Hz, $^4J_{HP}$ 1.1 Hz, 6H).

**4-formylphenyl diethyl phosphate (6)**

**$^1$H NMR** (399.6 MHz, $CDCl_3$): δ (ppm) 10.00 (s, 1H), 7.92 (d, $^3J_{HH}$ = 8.5 Hz, 2H), 7.41 (d, $^3J_{HH}$ = 8.5 Hz, 2H), 4.27 (m, 4H), 1.39 (td, $^3J_{HH}$ = 7.1 Hz, $^4J_{HP}$ 1.0 Hz, 6H).

**4-cyanophenyl diethyl phosphate (7)**

**$^1$H NMR** (399.6 MHz, $CDCl_3$): δ (ppm) 7.68 (dd, $^3J_{HH}$ = 8.9, $^4J_{HP}$ = 0.5 Hz, 2H), 7.37 (dd, $^3J_{HH}$ = 8.9 Hz, $^4J_{HP}$ 0.9 Hz, 2H), 4.26 (m, 4H), 1.39 (td, $^3J_{HH}$ = 7.1 Hz, $^4J_{HP}$ 1.1 Hz, 6H).

**4-acetylphenyl diethyl phosphate (8)**

**$^1$H NMR** (399.6 MHz, $CDCl_3$): δ (ppm) 7.99 (d, $^3J_{HH}$ = 8.4 Hz, 2H), 7.34 (dd, $^3J_{HH}$ = 8.9, $^4J_{HP}$ 0.9 Hz, 2H), 4.26 (m, 4H), 2.61 (s, 3H), 1.39 (td, $^3J_{HH}$ = 7.1 Hz, $^4J_{HP}$ 1.1 Hz, 6H).

**3-cyanophenyl diethyl phosphate (9)**

**$^1$H NMR** (399.6 MHz, $CDCl_3$): δ (ppm) 7.56-7.45 (m, 4H), 4.26 (m, 4H), 1.40 (td, $^3J_{HH}$ = 7.1 Hz, $^4J_{HP}$ 1.1 Hz, 6H).

**3-chlorophenyl diethyl phosphate (10)**

**$^1$H NMR** (399.6 MHz, $CDCl_3$): δ (ppm) 7.30 (d, $^3J_{HH}$ = 8.2 Hz, 1H), 7.27 (m, 1H), 7.20-7.14 (m, 2H), 4.25 (m, 4H), 2.61 (s, 3H), 1.38 (td, $^3J_{HH}$ = 7.1 Hz, $^4J_{HP}$ 1.0 Hz 6H).

#### 2. Supplementary Figures

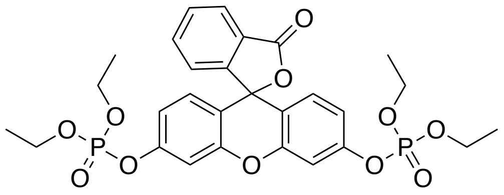

**1**

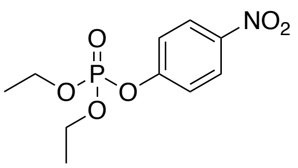

**2**

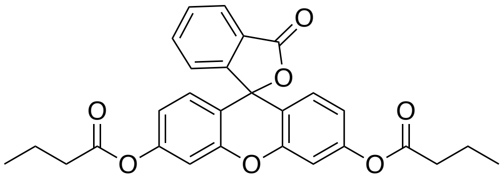

**3**

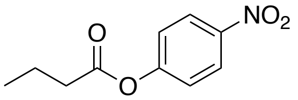

**4**

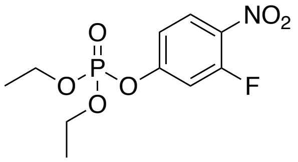

**5**

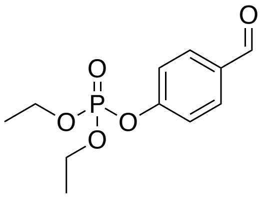

**6**

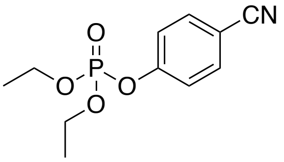

**7**

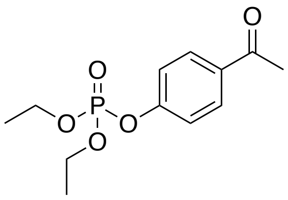

**8**

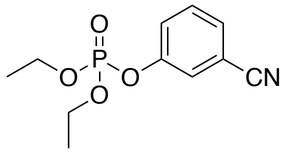

**9**

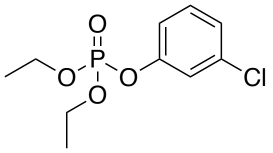

**10**

**Figure S1: Structures of the substrates used in this study. 1**: Fluorescein di(diethylphosphate) (FDDEP); **2:** Paraoxon-ethyl (PXN); **3:** Fluorescein dibutyrate; **4:** *p*-Nitrophenyl butyrate; **5**: 3-Fluoro-4-nitrophenyl diethylphosphate; **6**: 4-Formylphenyl diethylphosphate, **7**: 4-Cyanophenyl diethylphosphate, **8**: 4-Acetylphenyl diethylphosphate; **9**: 3-Cyanophenyl diethylphosphate; **10**: 3-Chlorophenyl diethylphosphate

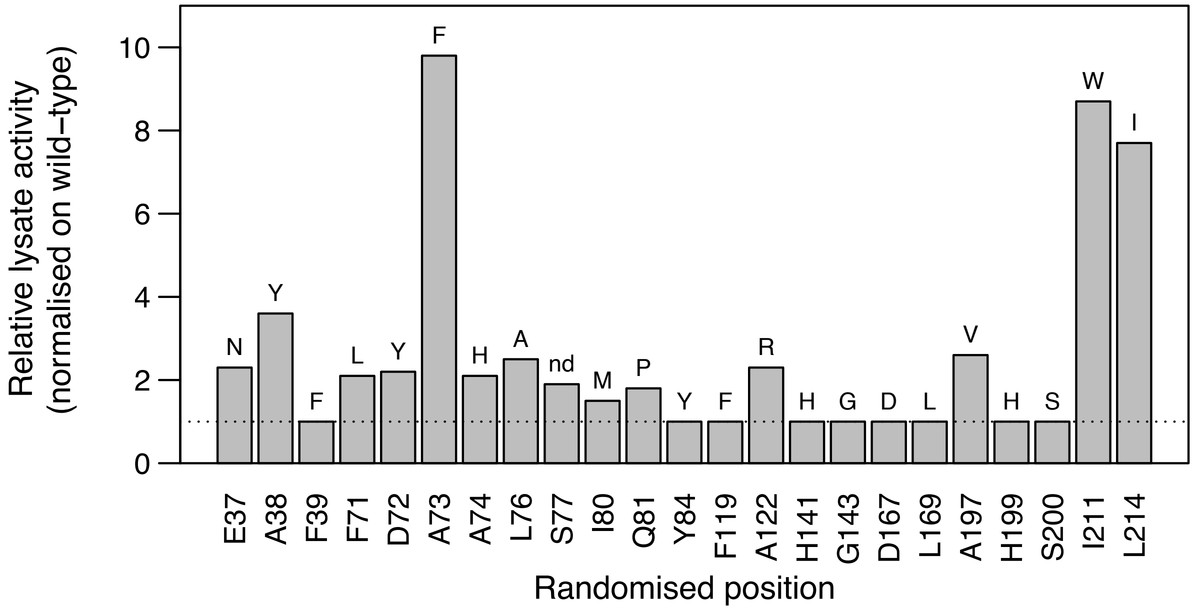

**Figure S2: Structure-guided mutational active-site scanning.** Positions lining the active site of P91 were individually randomised and screened for phosphotriesterase activity in microtiter plates using 1 μM FDDEP. Over four-fold oversampling of the theoretical diversity at each position ensures that with high probability every single amino acid substitution appears in the screen. The bars show the lysate activity (rate of product formation) of the respective best-performing clone at each position (indicated in single-letter amino acid code), relative to the wild type. The dotted line indicates wild-type level activity. The substituting amino acid of the respective most active clone is indicated on top of the bar; nd: not determined. Note that the triad residues Asp167 and His199 were also included but did not show tolerance to mutation.

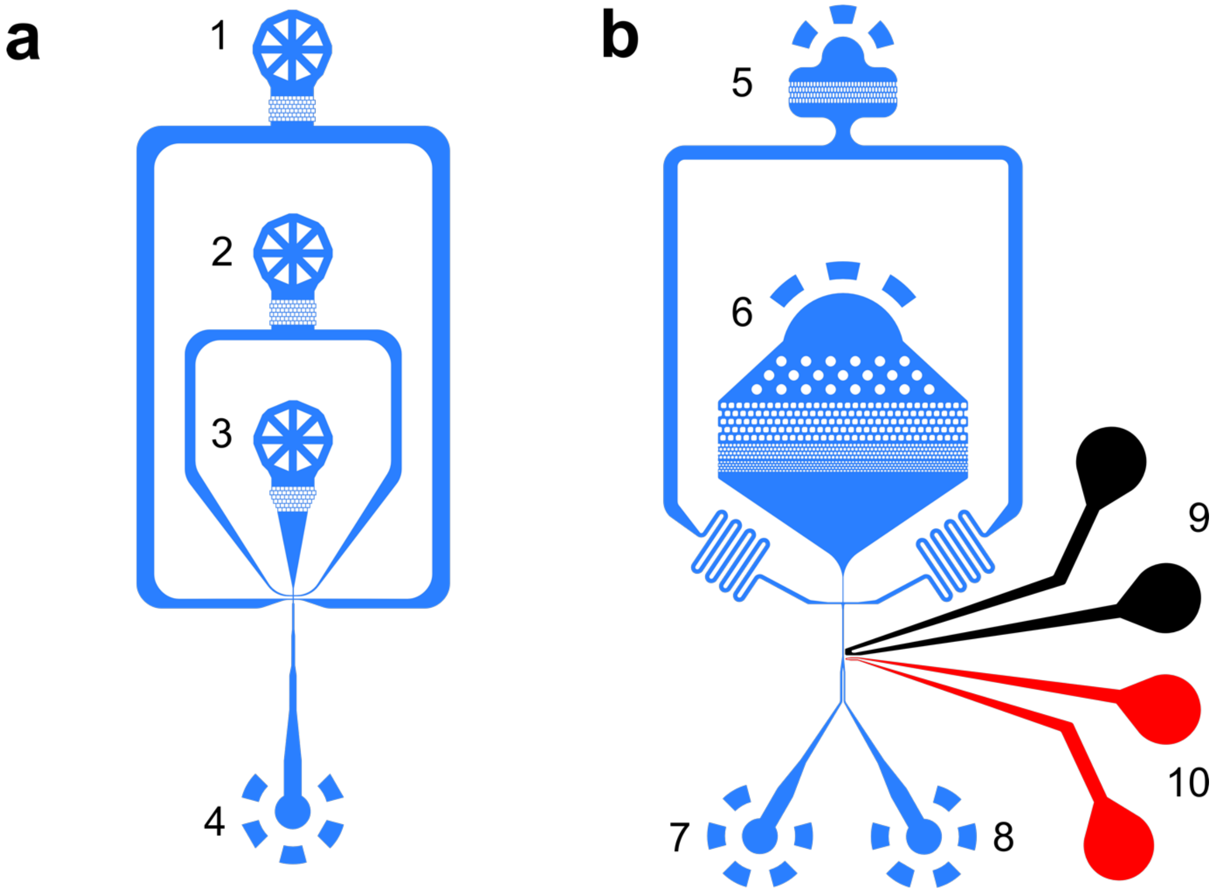

**Figure S3: Design of microfluidic chips for droplet generation and sorting (off-line droplet incubation). (a)** Flow-focussing chip (depth: 12 µm) for droplet generation with (1) oil/surfactant mixture inlet, (2) inlet for substrate/lysis agent mixture, (3) inlet for cell suspension, and (4) outlet for droplet collection. **(b)** Droplet sorting chip (depth: 21.5–23 µm) for fluorescence-activated droplet sorting with (5) inlet for spacing oil, (6) inlet for droplets, (7) waste outlet, (8) hit outlet, (9) ground electrode (+, black), and (10) signal electrode (–, red). This figure is adapted from Neun *et al.* 2019.[6]

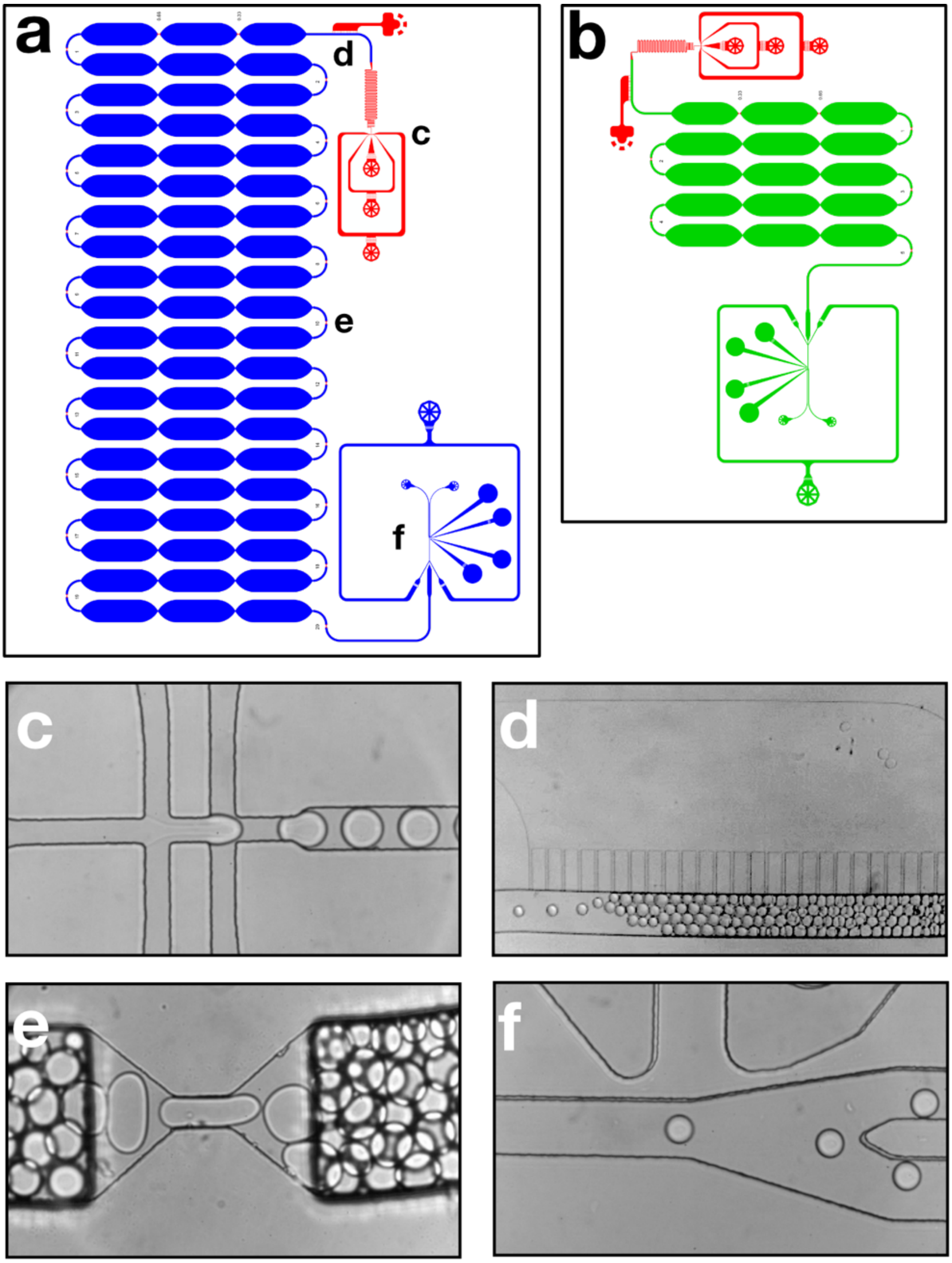

**Figure S4: Design of microfluidic chips for on-chip droplet incubation.** For short incubation times in evolution round 2, requiring on-chip incubation, an integrated chip was used, combining a flow-focussing module, a delay line, and a sorting module on a single device. The chips are two-layered such that the deeper delay line reduces backpressure. Red areas are 15 µm deep, green areas 28–29 µm, and blue areas 46–48 µm. **(a)** For initial stringency adjustments, a chip with a long delay line of 20 loops was used. **(b)** For library sorting, a shorter delay line consisting of five loops was used. **(c)** Monodisperse water-in-oil droplets are generated in a **flow-focusing nozzle**, co-encapsulating bacteria, the fluorogenic substrate and lysis agent. **(d)** An **oil extractor** ensures dense packing and equal incubation time for all droplets in the **(e) delay line chamber.** The delay line chambers possess mixing constrictions and shallow windows to monitor droplet fluorescence during incubation. **(f)** In the **sorting junction** droplets are electrophoretically sorted according to their fluorescence with up to kHz frequencies.

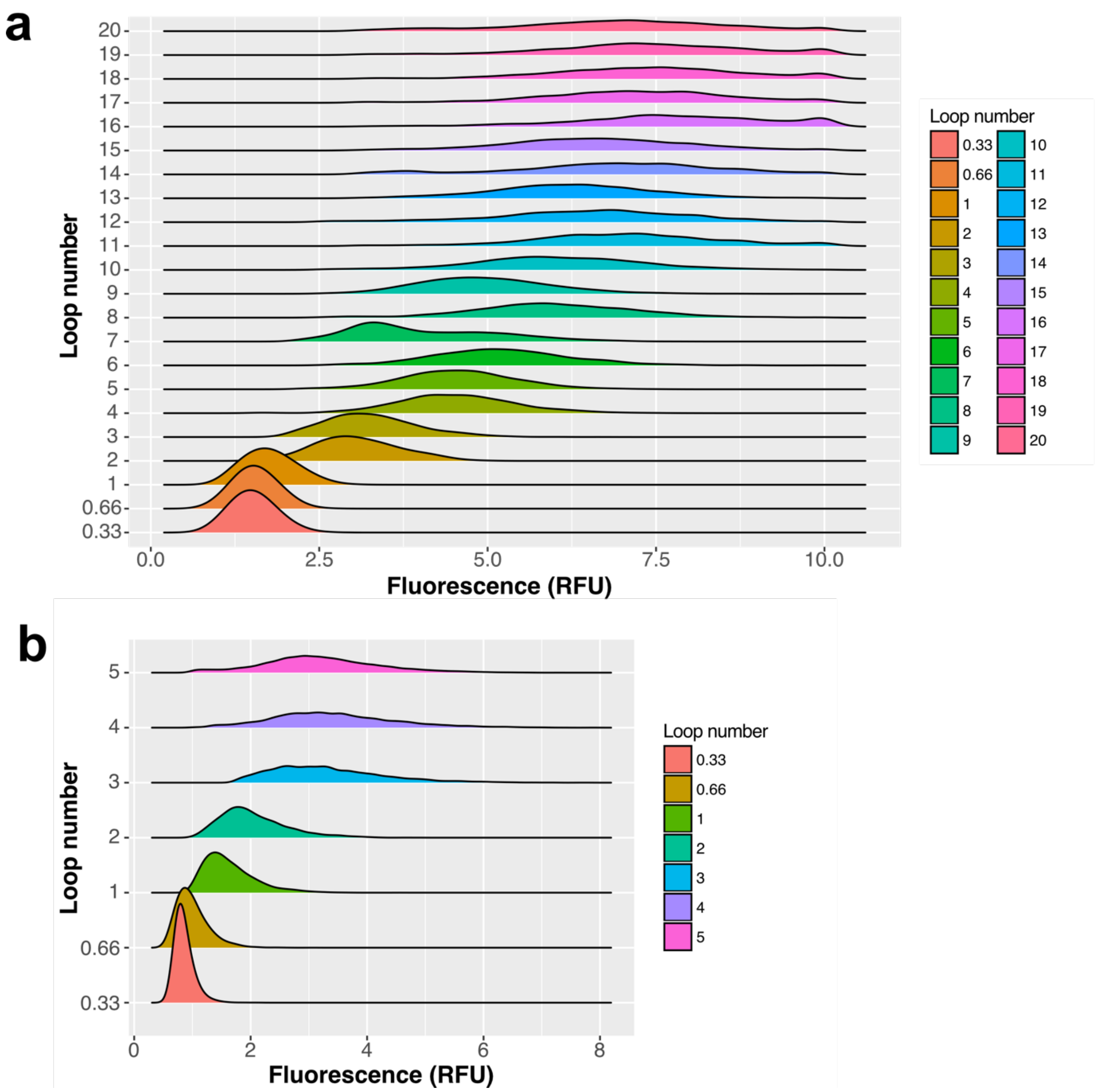

**Figure S5: On-chip fluorescence measurements for the adjustment of reaction time and sorting stringency.** Fluorescence distribution of 10 000 droplets in relative fluorescence units (RFU) at different points in the delay line. Fluorescence measurements were taken at the outer constrictions of the delay line loops (**Figure S4e**) in order to on follow the progress of the reaction in droplets. The loop number indicates the place of the constriction along the delay line where the laser was placed for measurements and is a proxy of reaction time. 0.33 and 0.66 refer to measurements at constrictions within the first loop of the delay line (after a third and two thirds of the first loop length, respectively). With the chosen flow rates, 20 loops correspond to ≈ 28 min in the long chip and five loops correspond to ≈ 4.5 min incubation time in the shorter and shallower library sorting chip. For initial stringency adjustments, a chip with a long delay line of 20 loops **(a)** was designed and cells expressing the parent variant for round 2, P91-R1, were encapsulated and incubated in the delay line. Fluorescence distribution increases linearly in the early loops and the reaction saturates in the later loops. With the aim to sort the library within the early linear phase of the reaction, this extent of reaction progress provided the basis for choosing five loops for the library sorting chip **(b).**

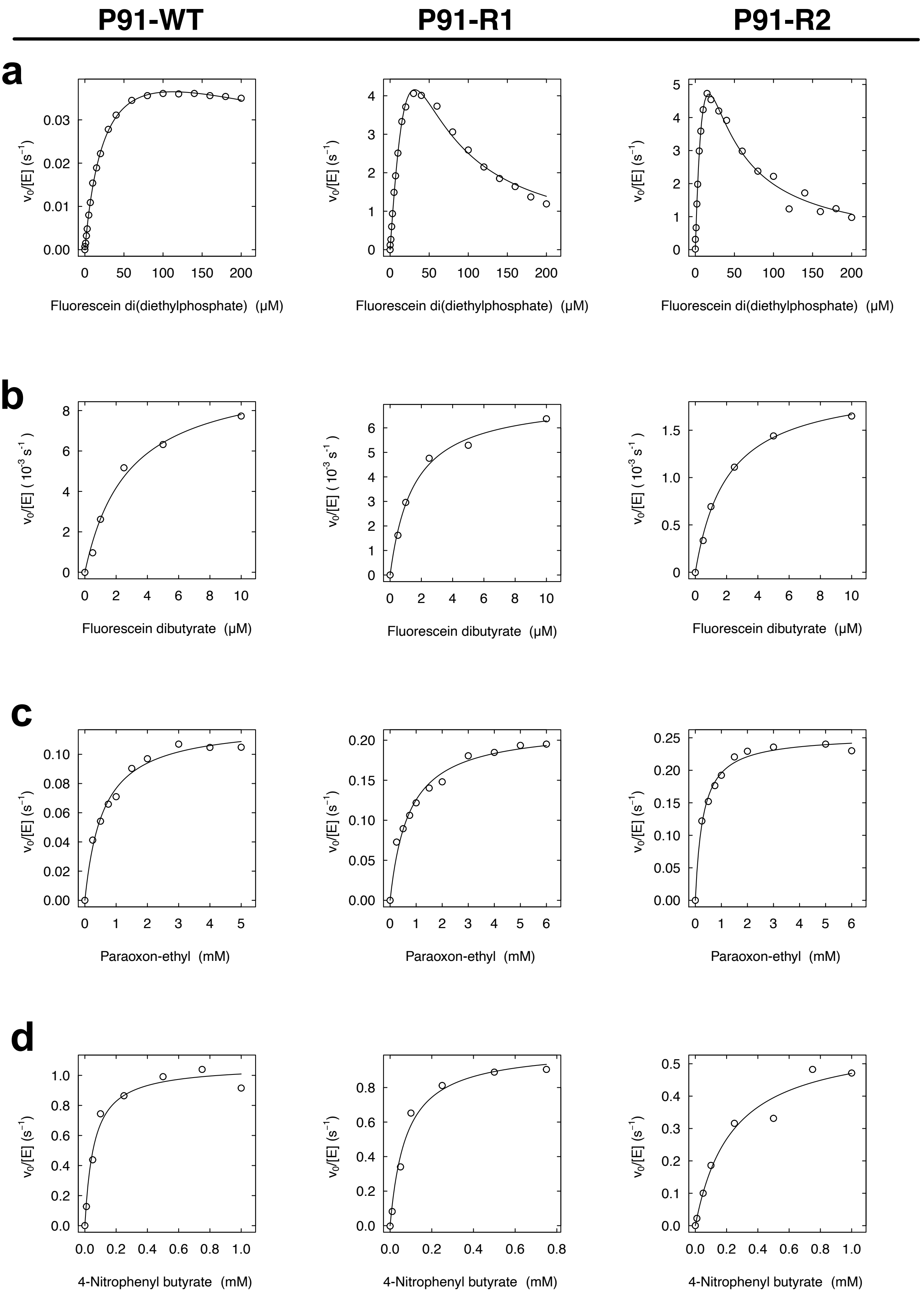

**Figure S6: Michaelis-Menten plots of steady-state kinetics** of P91-WT, P91-R1, and P91-R2 with **(a)** fluorescein di(diethylphosphate) **1, (b)** fluorescein dibutyrate **3**, **(c)** paraoxon-ethyl **2**, and **(d)** *p*-nitrophenyl butyrate **4**. Measured in 50 mM HEPES-NaOH, 150 mM NaCl, 1 mM TCEP, pH 8.0 at 25 ˚C.

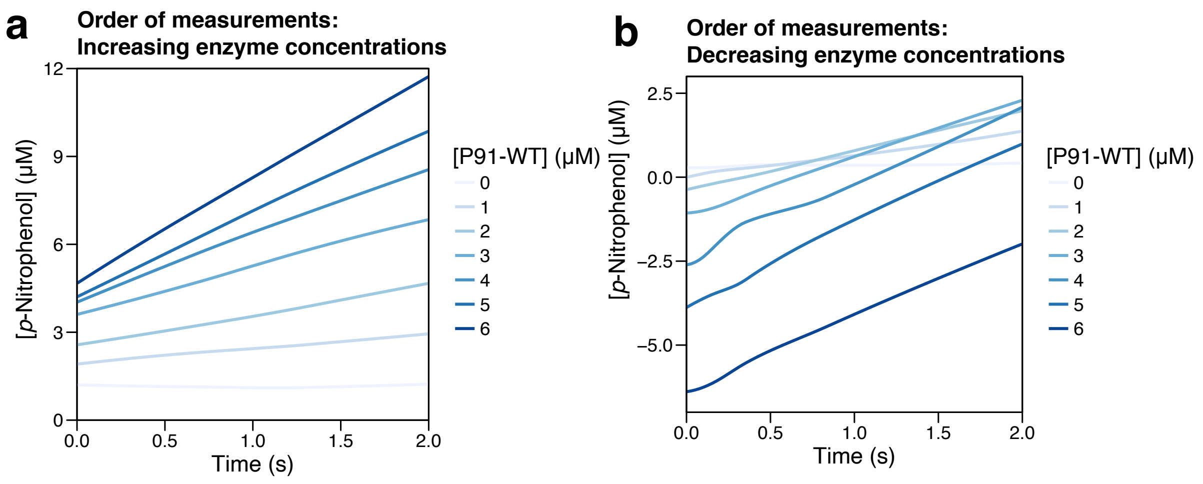

**Figure S7: Wild-type P91 (containing a cysteine triad) does not show biphasic burst kinetics**. Stopped-flow kinetic traces of P91-WT reacting with 2 mM paraoxon-ethyl, measured in 50 mM HEPES-NaOH, 150 mM NaCl, 1 mM TCEP, pH 8.0 at 25 ˚C at varying enzyme concentrations (0–6 µM, blue shades). Absorbance values are shown in units of corresponding concentration of released *p*-nitrophenol (PNP). **(a)** Time course measurements taken in order of *increasing* enzyme concentrations. **(b)** Time course measurements taken in order of *decreasing* enzyme concentrations. Although an absorbance offset is observable its amplitude does not depend on the enzyme concentration used but rather on the order of measurements (time-dependent). The absorbance offset is therefore probably consequence of chemical background hydrolysis, refuting the hypothesis that a very fast burst could be occurring in the dead time of the stopped-flow instrument (≈ 1–5 ms).

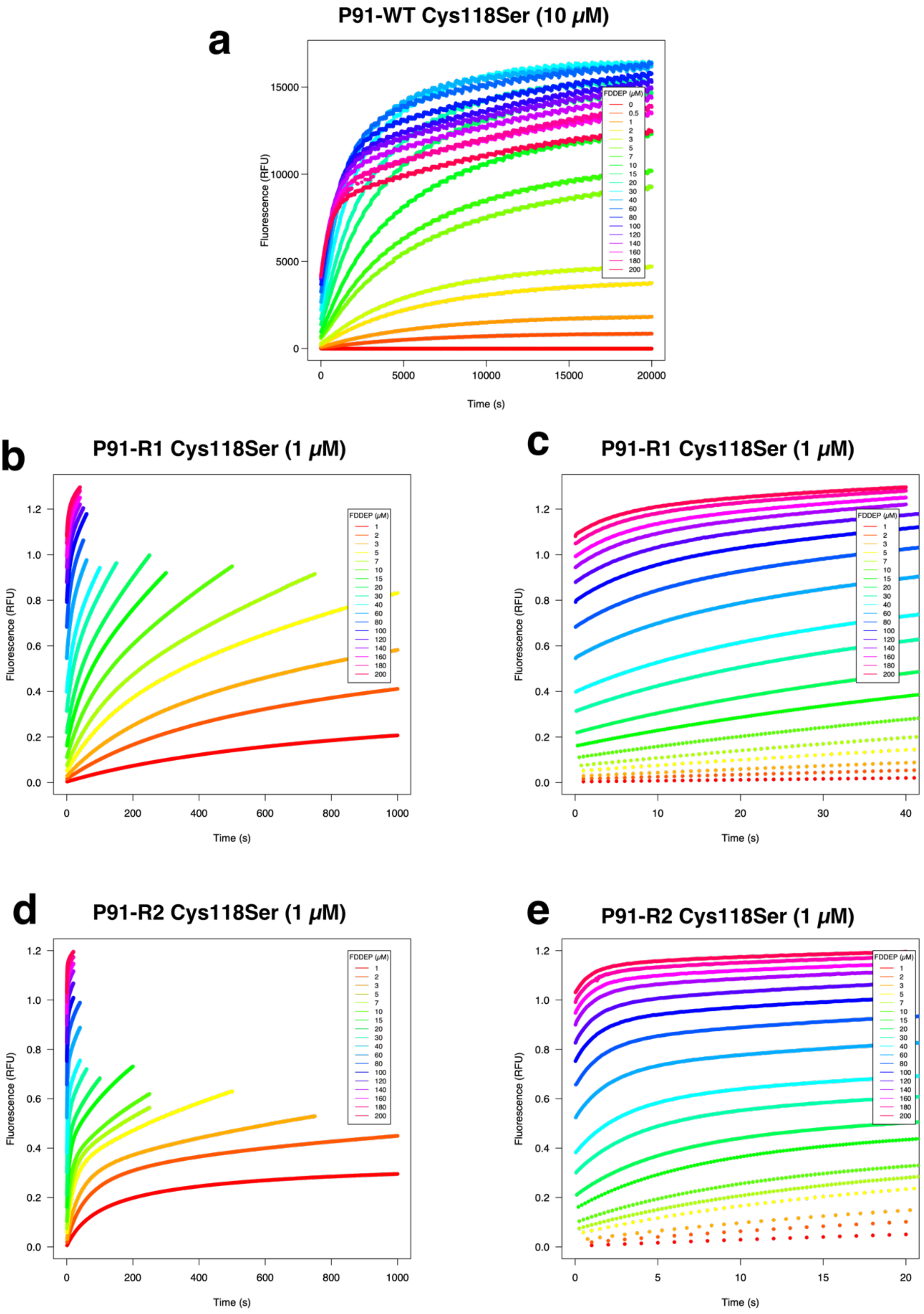

**Figure S8: Stopped-flow reaction traces** of all characterised P91 Cys118Ser variants with a concentration range of FDDEP (1–200 μM), measured in 50 mM HEPES-NaOH, 150 mM NaCl, 1 mM TCEP, pH 8.0 at 25 °C. Measurement time was varied with different substrate concentrations and was increased at low substrate concentrations according to reaction rate. **a)** P91-WT Cys118Ser (10 μM). **b)** P91-R1 Cys118Ser (1 μM), full time range. **c)** P91-R1 Cys118Ser (1 μM), close-up of the initial 40 s. **d)** P91-R2 Cys118Ser (1 μM), full time range. **e)** P91-R1 Cys118Ser (1 μM), close-up of the initial 20 s.

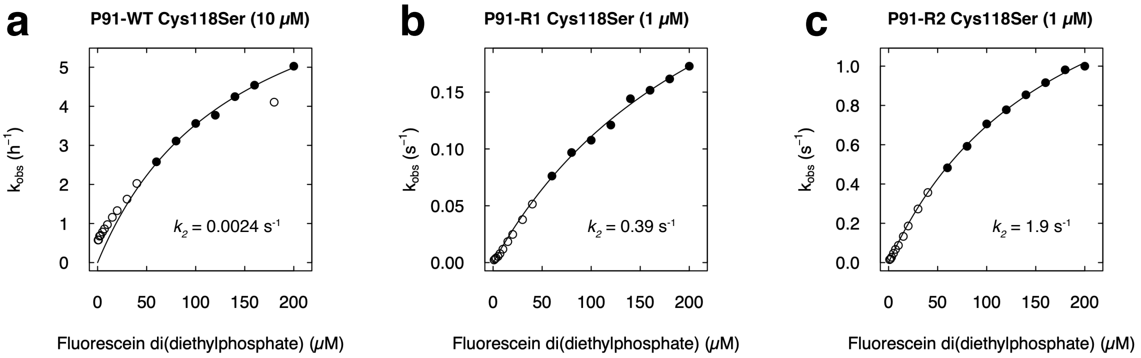

**Figure S9: Determination of the phosphorylation rate k2.** The parameter $k_{obs}$, determined from exponential fit of the kinetic burst traces measured with the substrate FDDEP, was plotted against substrate concentration and fitted to a modified Michaelis-Menten equation as described in the Methods section. Only substrate concentrations of ≥ 60 µM FDDEP (where substrate concentration >> enzyme concentration) were considered for the fit (filled dots). Phosphorylation rates and standard errors of the fit were determined as **a)** P91-WT Cys118Ser: (2.4 ± 0.10) · $10^{-3}$ $s^{-1}$. **b)** P91-R1 Cys118Ser: (3.9 ± 0.32) · $10^{-1}$ $s^{-1}$. **c)** P91-R2 Cys118Ser: (1.9 ± 0.067) $s^{-1}$. As the curves do not reach saturation due to low substrate solubilities, the values determined for $k_2$ may contain an error larger than that of the fit.

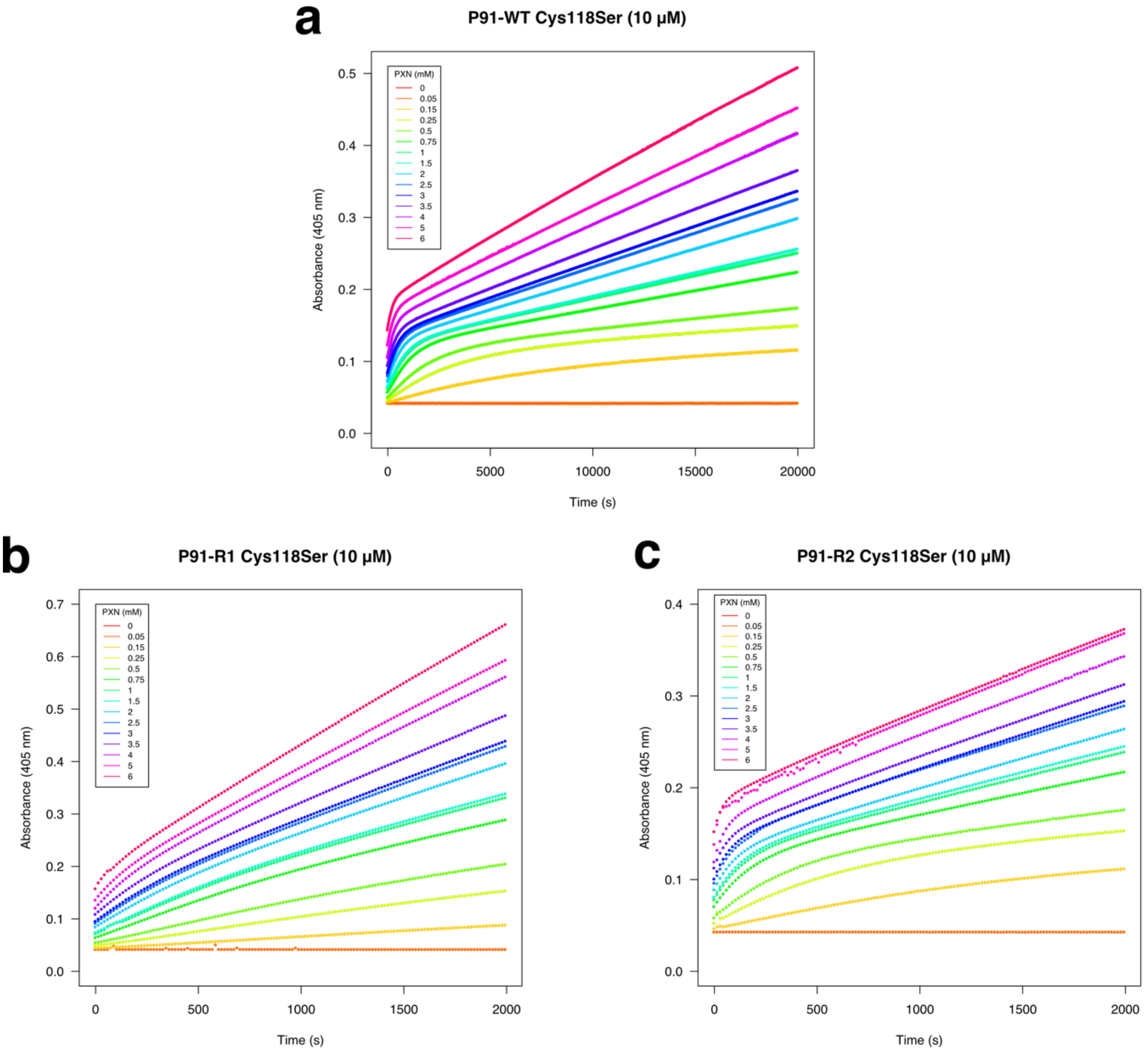

**Figure S10: Burst traces of all characterised P91 Cys118Ser variants** with a concentration range of paraoxon-ethyl (0.05–6 mM), measured in 50 mM HEPES-NaOH, 150 mM NaCl, 1 mM TCEP, pH 8.0 at 25 ˚C. **a)** P91-WT Cys118Ser (10 µM). **b)** P91-R1 Cys118Ser (10 µM). **c)** P91-R2 Cys118Ser (10 µM).

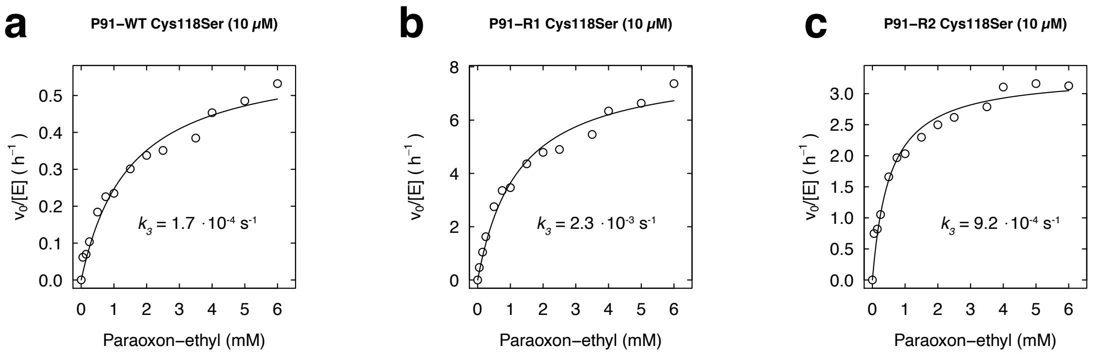

**Figure S11: Determination of the de-phosphorylation rate $k_3$.** The initial rate $v_0$ from the second phase of the burst kinetics with the substrate paraoxon-ethyl was fitted to the Michaelis-Menten equation to determine $k_{cat}$ and thus $k_3$, as described in the Methods section. In the case of P91-R1 Cys118Ser, where the burst is less pronounced (probably due to a lower k2 with the substate paraoxon), $k_{cat}$ gives an upper boundary and thus $k_3$ might be lower. De-phosphorylation rates were determined as **a)** P91-WT Cys118Ser: $(1.7 \pm 0.10) \times 10^{-4}$ $s^{-1}$. **b)** P91-R1 Cys118Ser: $(2.3 \pm 0.11) \times 10^{-3}$ $s^{-1}$. **c)** P91-R2 Cys118Ser: $(9.2 \pm 0.34) \times 10^{-4}$ $s^{-1}$.

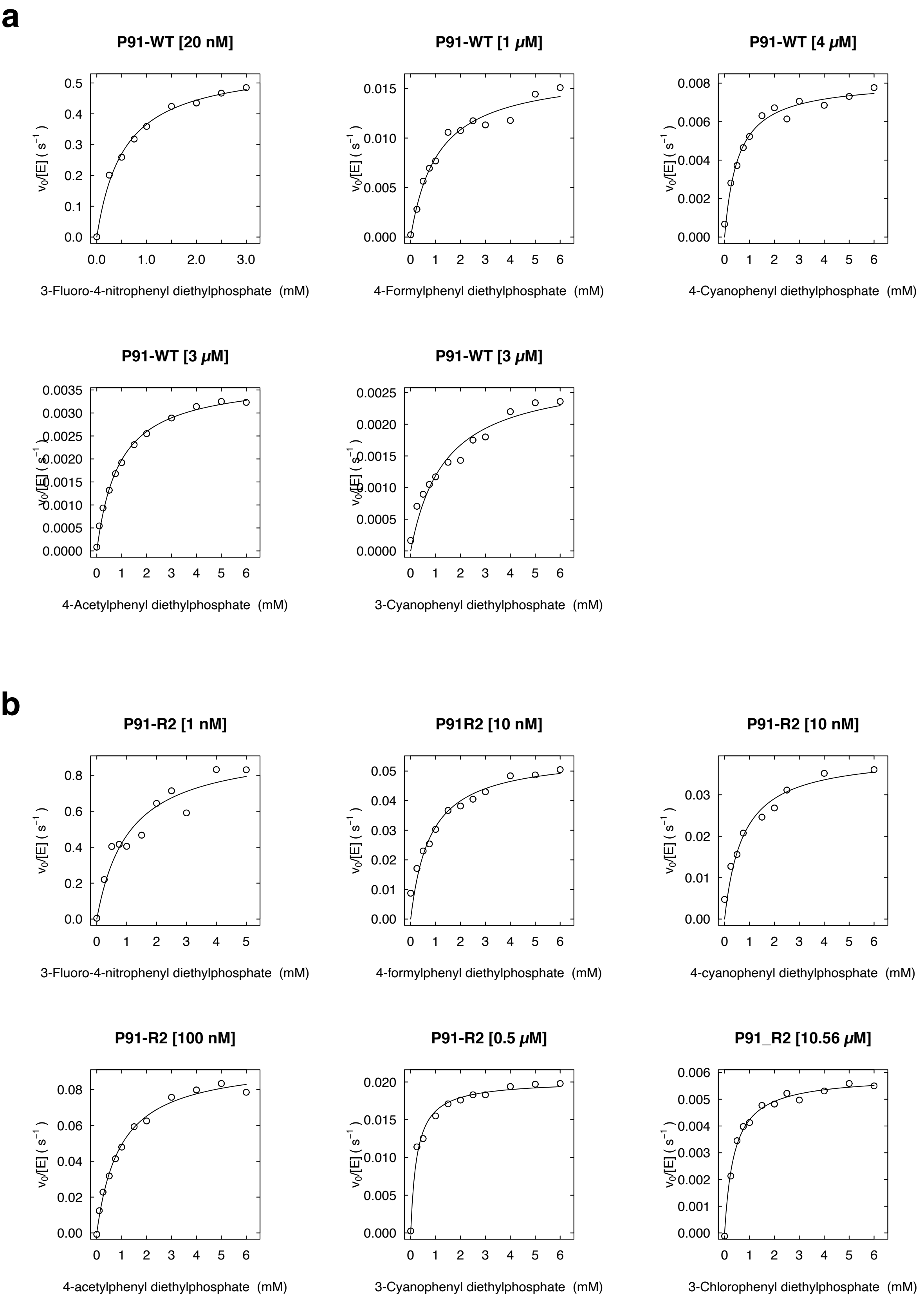

**Figure S12: Michaelis-Menten plots for steady-state kinetics of (a) P91-WT and (b) P91-R2 with linear-free energy relationship substrates** 5–10 (see axis labels), measured in 50 mM HEPES-NaOH, 150 mM NaCl, pH 8.0 at 25 °C.

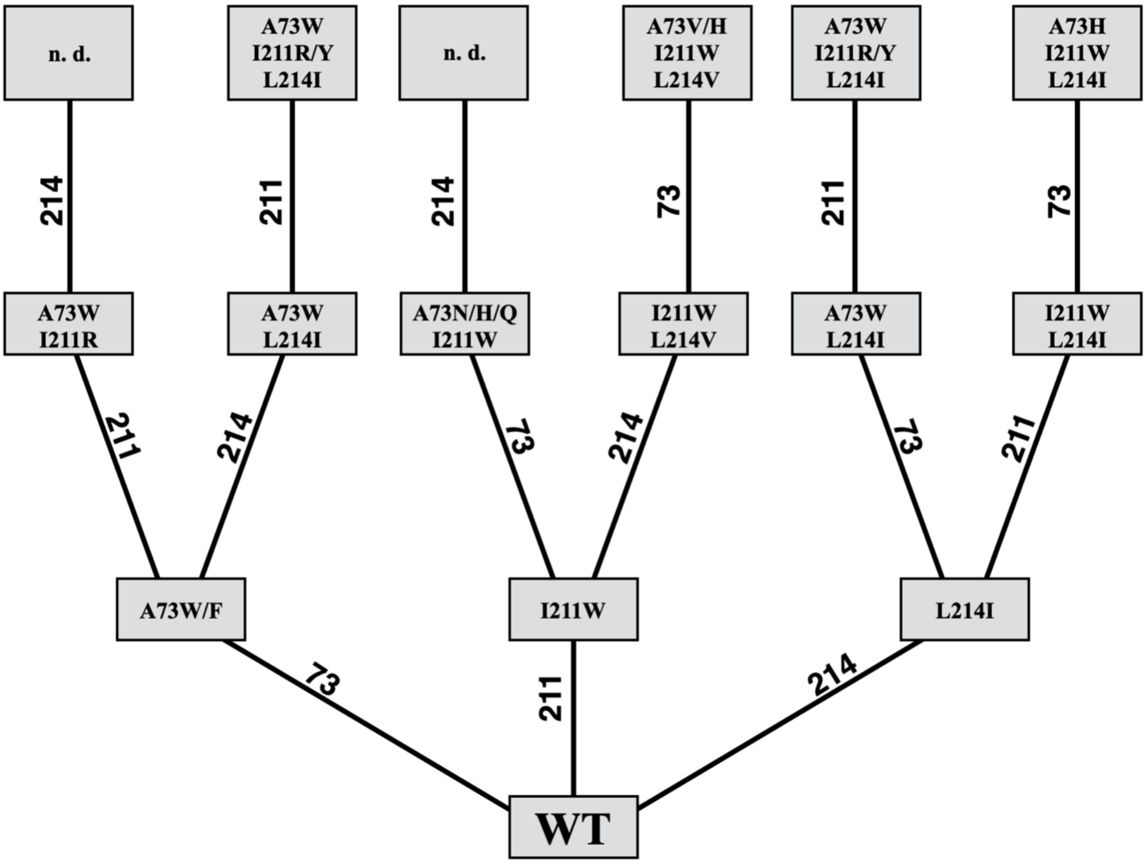

**Figure S13: Iterative saturation mutagenesis (ISM) of P91 at the three positions A73, I211 and L214.** Each position was randomised and screened for activity towards substrate **1** (FDDEP) in 96-well plates. Edges represent a screening campaign with the respective randomised residue indicated on the line, boxes represent the best identified mutant which served as starting variant for the next randomisation. In some cases, results were slightly ambiguous due to approximately equally improved mutants. n.d.; not determined.

### 3. Kinetic data and comparisons

**Table S1: Steady-state catalytic parameters** for P91-WT, P91-R1, and P91-R2, measured in 50 mM HEPES-NaOH, 150 mM NaCl, 1 mM TCEP, pH 8.0 at 25 ˚C.

| | P91-WT | | | | P91-R1 | | | | P91-R2 | | | |
|---|---|---|---|---|---|---|---|---|---|---|---|---|
| **Substrate** | $k_{cat}$ ($s^{-1}$) | $K_M$ (μM) | $K_i$ (μM) | $k_{cat}/K_M$ ($M^{-1}\cdot s^{-1}$) | $k_{cat}$ ($s^{-1}$) | $K_M$ (μM) | $K_i$ (μM) | $k_{cat}/K_M$ ($M^{-1}\cdot s^{-1}$) | $k_{cat}$ ($s^{-1}$) | $K_M$ (μM) | $K_i$ (μM) | $k_{cat}/K_M$ ($M^{-1}\cdot s^{-1}$) |
| **FDDEP** | 0.054 | 27 | 470 | $2.0 \cdot 10^3$ | 37[a] | 120[a] | 7.9 | $3.0 \cdot 10^5$ | 15[a] | 20[a] | 14 | $7.8 \cdot 10^5$ |
| **Fluorescein dibutyrate** | 0.010 | 2.8 | -- | $3.6 \cdot 10^3$ | 0.0072 | 1.5 | -- | $4.8 \cdot 10^3$ | 0.0020 | 2.0 | -- | $1.0 \cdot 10^3$ |
| **Paraoxon-ethyl** | 0.12 | 580 | -- | $2.1 \cdot 10^2$ | 0.22 | 720 | -- | $3.0 \cdot 10^2$ | 0.25 | 290 | -- | $7.5 \cdot 10^2$ |
| ***p*-Nitrophenyl butyrate** | 1.1 | 58 | -- | $1.9 \cdot 10^4$ | 1.0 | 77 | -- | $1.3 \cdot 10^4$ | 0.58 | 230 | -- | $2.5 \cdot 10^3$ |

*a) Due to strong substrate inhibition, estimates of $k_{cat}$ and $K_M$ are strong extrapolations, only $k_{cat}/K_M$ can be regarded as precise (see Michaelis-Menten plot, Figure **S6a**)*

*b) Values differ slightly from previously published values[12], presumably due to differences in affinity tag and buffer conditions.*

**Table S2: P92-R2 rivals the efficiencies of engineered and naturally evolved metal-dependent phosphotriesterases.** Comparison of kinetic parameters of promiscuous, engineered, or naturally evolved phosphotriesterases from different protein superfamilies: ABH, α/β-hydrolases; BP, β-propellers; MBL, metallo-β-lactamases; AH, amidohydrolases. Substrates between enzymes differ in leaving group but are all diethyl-substituted phosphotriesters. n.d., not determined; n.a., not applicable.

| **Label in figure 4** | **Enzyme** | **Protein superfamily** | **Catalytic mechanism** | **Organism of origin** | **Substrate** | $k_{cat}/K_M$ **($10^3\cdot M^{-1}\cdot s^{-1}$)** | **Relative increase in $k_{cat}/K_M$** | **Rounds[a]** | **Reference** |
|---|---|---|---|---|---|---|---|---|---|
| a | P91-WT | ABH | nucleophilic | metagenomic | Fluorescein di(diethylphosphate) | 2.0 | n.a. | n.a. | this work |
| b | P91-R2 | ABH | nucleophilic | metagenomic | Fluorescein di(diethylphosphate) | 780 | 400 | 2 | this work |
| c | BChE G117H | ABH | nucleophilic | *Homo sapiens* | Echothiophate | 0.45 | n.d.[b] | 1[b] | [17] |
| d | *Lc*αE7 G137D | ABH | nucleophilic | *Lucilia cuprina* | Diethylumbelliferyl phosphate | 0.16 | n.a. | n.a. | [18] |
| e | AiiA-WT | MBL | metal | *Bacillus thuringiensis* | Paraoxon-ethyl | 0.51 | n.a. | n.a. | [19] |
| | AiiA-WT | MBL | metal | *Bacillus thuringiensis* | Parathion-methyl[c] | 70 | n.a. | n.a. | [19] |
| f | AiiA-R6 | MBL | metal | *Bacillus thuringiensis* | Paraoxon-ethyl | 540 | 1100 | 6 | [19] |
| g | MPH | MBL | metal | *Pseudomonas sp. WBC-3* | Paraoxon-ethyl | 16 | n.a. | n.a. | [19] |
| h | rePON1-G3C9 | BP | metal | *Oryctolagus cuniculus*[d] | DEPCyc | 11 | n.a. | n.a. | [20] |
| i | rePON1-G3C9-3.2PC | BP | metal | *Oryctolagus cuniculus*[d] | DEPCyc | 1,400 | 130 | 6 | [21] |
| j | *Dr*PLL-WT | AH | metal | *Deinococcus radiodurans* | Paraoxon-ethyl | 0.029 | n.a. | n.a. | [22] |
| k | *Dr*PLL.10 | AH | metal | *Deinococcus radiodurans* | Paraoxon-ethyl | 20 | 690 | 10 | [22] |
| | *Dr*PLL.10 | AH | metal | *Deinococcus radiodurans* | O-isopbutyl-O-4-nitrophenyl-$R_P$-methylphosphonate[e] | 1100 | 37000 | 10 | [22] |
| l | *Bd*PTE | AH | metal | *Brevundimonas diminuta* | Paraoxon-ethyl | 120000 | n.a. | n.a. | [22] |
| | *Bd*PTE | AH | metal | *Brevundimonas diminuta* | Paraoxon-ethyl | 180000 | 1.8[f] | 1 | [23] |
| | *Dr*-OPH w.t. | AH | metal | *Deinococcus radiodurans* | Paraoxon-ethyl | 0.0014 | n.a. | n.a. | [24] |

| | | | | | | | | | |
|---|---|---|---|---|---|---|---|---|---|
| | *Dr*-OPH D71G/ E101G/ V235L | AH | metal | *Deinococcus radiodurans* | Paraoxon-ethyl | 0.77 | 560 | 3 | [24] |
| | *Gka*P-PLL WT | AH | metal | *Geobacillus kaustophilus* | Paraoxon-ethyl | 0.13 | n.a. | n.a. | [25,26] |
| | *Gka*P-PLL ML7-B6 | AH | metal | *Geobacillus kaustophilus* | Paraoxon-ethyl | 79 | 610 | 6 | [25,26] |
| | OpdA WT | AH | metal | *Agrobacterium radiobacter* | Z-Chlorfenvinphos | 0.0096 | n.a. | n.a. | [27] |
| | OpdA W131H/ F132A | AH | metal | *Agrobacterium radiobacter* | Z-Chlorfenvinphos | 4.6 | 480 | 2[g] | [27] |

a) Number of laboratory evolution rounds, meaning successive iterations of diversification and screening.
b) A single substitution was rationally engineered into the wild-type precursor which has no detectable activity.
c) Substrate with highest final activity in this evolution campaign.
d) Engineered rabbit gene with contributions from the respective human, mouse, and rat genes through DNA shuffling during directed evolution.
e) Substrate with highest final activity and highest relative activity increase in this evolution campaign.
f) 1.8-fold increase in $k_{cat}/K_M$ as compared to a wild-type $k_{cat}/K_M$ of $9.9 \cdot 10^7$ $M^{-1} \cdot s^{-1}$, as measured in this study. In addition, the selected *Bd*PTE variant displayed a 63-fold increase in $k_{cat}$.
g) Two steps of rational design

**Table S3: Microscopic rate constants** for the formation ($k_2$) and the breakdown ($k_3$) of the covalent intermediate with nucleophile-exchanged variants. $k_2$ was determined with the substrate FDDEP, $k_3$ was determined with the substrate paraoxon-ethyl (PXN).

| Enzyme variant | $k_2$ ($s^{-1}$) | $k_3$ ($s^{-1}$) | $k_3/K_M$ (*PXN*, $M^{-1}$ $s^{-1}$) |
|---|---|---|---|
| P91-WT Cys118Ser | $2.4 \cdot 10^{-3}$ | $1.7 \cdot 10^{-4}$ | 0.11 |
| P91-R1 Cys118Ser | $3.9 \cdot 10^{-1}$ | $2.3 \cdot 10^{-3}$ | 1.8 |
| P91-R2 Cys118Ser | $1.9 \cdot 10^{0}$ | $9.2 \cdot 10^{-4}$ | 1.7 |

**Table S4: Properties of phosphotriester substrates (paraoxon-ethyl derivatives)** used for the linear free-energy relationships.

| Substrate | Leaving group | p$K_a$ | Detection wavelength (nm) | Extinction coefficient ($M^{-1}$) |
|---|---|---|---|---|
| 5 | 3-fluoro-4-nitrophenol | 5.94 | 390 | 10473.2 |
| 2 | 4-nitrophenol | 7.14 | 405 | 10038.1 |
| 6 | 4-hydroxybenzaldehyde | 7.66 | 330 | 12483.8 |
| 7 | 4-cyanophenol | 7.95 | 275 | 6721.7 |
| 8 | 4-hydroxyacetophenone | 8.05 | 320 | 7242.1 |
| 9 | 3-cyanophenol | 8.61 | 295 | 1246.2 |
| 10 | 3-chlorophenol | 9.12 | 276 | 796.0 |

**Table S5: Steady-state catalytic parameters for linear free-energy relationship of P91-WT**, measured in 50 mM HEPES-NaOH, 150 mM NaCl, 1 mM TCEP, pH 8.0.

| Substrate | p$K_a$ of leaving group | Enzyme concentration in the measurement | $k_{cat}$ ($s^{-1}$) | $K_M$ (mM) | $k_{cat}/K_M$ ($M^{-1} \cdot s^{-1}$) |
|---|---|---|---|---|---|
| 5 | 5.94 | 20 nM | 0.57 | 0.55 | 1000 |
| 2 | 7.14 | 0.5 µM | 0.12 | 0.58 | 210 |

| | | | | | |
|---|---|---|---|---|---|
| 6 | 7.66 | 1 μM | 0.017 | 1.1 | 15 |
| 7 | 7.95 | 4 μM | 0.0081 | 0.54 | 15 |
| 8 | 8.05 | 3 μM | 0.0038 | 0.9 | 4.2 |
| 9 | 8.61 | 3 μM | 0.0028 | 1.4 | 2.1 |
| 10 | 9.12 | 12.56 μM | *no turnover could be detected* | | |

**Table S6: Steady-state catalytic parameters for linear free-energy relationship of P91-R2**, measured in 50 mM HEPES-NaOH, 150 mM NaCl, 1 mM TCEP, pH 8.0.

| Substrate | p$K_a$ of leaving group | Enzyme concentration in the measurement | $k_{cat}$ ($s^{-1}$) | $K_M$ (mM) | $k_{cat}/K_M$ ($M^{-1}\cdot s^{-1}$) |
|---|---|---|---|---|---|
| 5 | 5.94 | 1 nM | 0.96 | 1.10 | 900 |
| 2 | 7.14 | 500 nM | 0.25 | 0.29 | 750 |
| 6 | 7.66 | 10 nM | 0.056 | 0.79 | 71 |
| 7 | 7.95 | 10 nM | 0.04 | 0.74 | 54 |
| 8 | 8.05 | 100 nM | 0.096 | 0.94 | 100 |
| 9 | 8.61 | 500 nM | 0.02 | 0.25 | 80 |
| 10 | 9.12 | 10.56 μM | 0.0059 | 0.39 | 15 |

**Table S7: Steady-state catalytic parameters of other P91 variants identified in round 2 for phosphotriester hydrolysis**, measured with the substrate FDDEP in 50 mM HEPES-NaOH, 150 mM NaCl, 1 mM TCEP, pH 8.0 at 25 ˚C. Enzyme concentrations were: 20 nM for P91-YYIA and 0.5 nM for P91-SKSL. Due to strong substrate inhibition, estimates of $k_{cat}$ and $K_M$ for both variants are extrapolations and only $k_{cat}/K_M$ can be regarded as precise.

| Enzyme variant | Mutations | $k_{cat}$ ($s^{-1}$) [a] | $K_M$ (μM) [a] | $K_i$ (μM) | $k_{cat}/K_M$ ($M^{-1}s^{-1}$) [b] |
|---|---|---|---|---|---|
| P91-YYIA | Ala38Tyr, Ala73Tyr, Leu76Ile, Ile211Trp, Leu214Val | 0.15 | 5.1 | 3.6 | $3.0 \times 10^4$ |
| P91-SKLS | Ala38Ser, Ala73Lys, Leu76Leu, Ile211Trp, Leu214Val | 1.1 | 7.9 | 5.7 | $1.4 \times 10^5$ |

### 4. Sequences

#### 4.1 Sequences of P91 variants and plasmid constructs

Notable codons are highlighted in bold, the coding region for the respective P91 variant is highlighted in green. XXX; randomised with degenerate codons NDT, VHG, TGG (22-codon trick).

```
>6xHis-P91-WT
MHHHHHHGGSMTARKVDYTDGATRCIGEFHWDEGKSGPRPGVVVFPEAFGLNDHAKERARRL
ADLGFAALAADMHGDAQVFDAASLSSTIQGYYGDRAHWRRRAQAALDALTAQPEVDGSKVAA
IGFCFGGATCLELARTGAPLTAIVTFHGGLLPEMAGDAGRIQSSVLVCHGADDPLVQDETMK
AVMDEFRRDKVDWQVLYLGNAVHSFTDPLAGSHGIPGLAYDATAEARSWTAMCNLFSELFG

>6xHis-P91-R1
MHHHHHHGGSMTARKVDYTDGATRCIGEFHWDEGKSGPRPGVVVFPEAFGLNDHAKERARRL
ADLGFAALAADMHGDAQVFDAASLSSTIQGYYGDRAHWRRRAQAALDALTAQPEVDGSKVAA
IGFCFGGATCLELARTGAPLTAIVTFHGGLLPEMAGDAGRIQSSVLVCHGADDPLVQDETMK
AVMDEFRRDKVDWQVLYLGNAVHSFTDPLAGSHGWPGVAYDATAEARSWTAMCNLFSELFG

>6xHis-P91-R2
MHHHHHHGGSMTARKVDYTDGATRCIGEFHWDEGKSGPRPGVVVFPELFGLNDHAKERARRL
ADLGFAALAADMHGDAQVFDEASVSSTIQGYYGDRAHWRRRAQAALDALTAQPEVDGSKVAA
IGFCFGGATCLELARTGAPLTAIVTFHGGLLPEMAGDAGRIQSSVLVCHGADDPLVQDETMK
AVMDEFRRDKVDWQVLYLGNAVHSFTDPLAGSHGWPGVAYDATAEARSWTAMCNLFSELFG

>pASK-IBA5plus_P91-A (library, round 1)
CTGGCAAGATTTTTTACGTAATAACGCTAAAAGTTTTAGATGTGCTTTACTAAGTCATCGCG
ATGGAGCAAAAGTACATTTAGGTACACGGCCTACAGAAAAACAGTATGAAACTCTCGAAAAT
CAATTAGCCTTTTTATGCCAACAAGGTTTTTCACTAGAGAATGCATTATATGCACTCAGCGC
AGTGGGGCATTTTACTTTAGGTTGCGTATTGGAAGATCAAGAGCATCAAGTCGCTAAAGAAG
AAAGGGAAACACCTACTACTGATAGTATGCCGCCATTATTACGACAAGCTATCGAATTATTT
GATCACCAAGGTGCAGAGCCAGCCTTCTTATTCGGCCTTGAATTGATCATATGCGGATTAGA
AAAACAACTTAAATGTGAAAGTGGGTCTTAAAAGCAGCATAACCTTTTTCCGTGATGGTAAC
TTCACTAGTTTAAAAGGATCTAGGTGAAGATCCTTTTTGATAATCTCATGACCAAAATCCCT
TAACGTGAGTTTTCGTTCCACTGAGCGTCAGACCCCGTAGAAAAGATCAAAGGATCTTCTTG
AGATCCTTTTTTTCTGCGCGTAATCTGCTGCTTGCAAACAAAAAAACCACCGCTACCAGCGG
TGGTTTGTTTGCCGGATCAAGAGCTACCAACTCTTTTTCCGAAGGTAACTGGCTTCAGCAGA
GCGCAGATACCAAATACTGTCCTTCTAGTGTAGCCGTAGTTAGGCCACCACTTCAAGAACTC
TGTAGCACCGCCTACATACCTCGCTCTGCTAATCCTGTTACCAGTGGCTGCTGCCAGTGGCG
ATAAGTCGTGTCTTACCGGGTTGGACTCAAGACGATAGTTACCGGATAAGGCGCAGCGGTCG
GGCTGAACGGGGGGTTCGTGCACACAGCCCAGCTTGGAGCGAACGACCTACACCGAACTGAG
ATACCTACAGCGTGAGCTATGAGAAAGCGCCACGCTTCCCGAAGGGAGAAAGGCGGACAGGT
ATCCGGTAAGCGGCAGGGTCGGAACAGGAGAGCGCACGAGGGAGCTTCCAGGGGGAAACGCC
TGGTATCTTTATAGTCCTGTCGGGTTTCGCCACCTCTGACTTGAGCGTCGATTTTTGTGATG
CTCGTCAGGGGGGCGGAGCCTATGGAAAAACGCCAGCAACGCGGCCTTTTTACGGTTCCTGG
CCTTTTGCTGGCCTTTTGCTCACATGACCCGACACCATCGAATGGCCAGATGATTAATTCCT
AATTTTTGTTGACACTCTATCATTGATAGAGTTATTTTACCACTCCCTATCAGTGATAGAGA
```

AAAGTGAAATGAATAGTTCGACAAAAATCTAGAAATAATTTTGTTTAACTTTAAGAAGGAGA
TATACAAATGGCTAGCTGGAGCCACCCGCAGTTCGAAAAAGGCGCCATGACAGCAAGAAAAG
TCGACTACACAGACGGTGCAACCCGCTGTATCGGTGAGTTTCATTGGGATGAAGGCAAGTCG
GGCCCGCGTCCCGGCGTGGTGGTCTTTCCCGAGGCTTTCGGCCTCAACGACCATGCCAAGGA
GCGCGCGCGGCGCCTTGCCGACCTCGGCTTTGCAGCCCTGGCGGCGGATATGCACGGAGACG
CCCAGGTTTTCGAT**NNK**GCGAGTCTCTCATCAACCATACAGGGCTACTACGGCGACCGCGCC
CACTGGCGACGTCGTGCGCAGGCAGCGCTCGATGCACTGACGGCACAGCCAGAGGTGGACGG
CAGCAAGGTGGCGGCCATCGGCTTTTGTTTCGGCGGTGCCACCTGCCTTGAACTGGCCCGCA
CAGGTGCGCCGCTGACCGCCATTGTCACCTTCCACGGCGGTTTGCTGCCGGAGATGGCAGGC
GATGCCGGACGGATCCAGTCCAGTGTTCTGGTGTGCCATGGCGCTGATGATCCGCTCGTACA
GGACGAAACCATGAAGGCCGTCATGGACGAGTTTCGTCGCGACAAGGTGGATTGGCAGGTGC
TCTACCTCGGAAATGCGGTACACAGTTTCACCGATCCACTCGCTGGCAGTCACGGC**NNK**CCC
GGG**NNK**GCCTATGACGCCACTGCCGAAGCCCGGTCGTGGACGGCCATGTGCAATCTGTTCAG
TGAACTGTTCGGCTGATGATATCTAACTAAGCTTGACCTGTGAAGTGAAAAATGGCGCACAT
TGTGCGACATTTTTTTTGTCTGCCGTTTACCGCTACTGCGTCACGGATCTCCACGCGCCCTG
TAGCGGCGCATTAAGCGCGGCGGGTGTGGTGGTTACGCGCAGCGTGACCGCTACACTTGCCA
GCGCCCTAGCGCCCGCTCCTTTCGCTTTCTTCCCTTCCTTTCTCGCCACGTTCGCCGGCTTT
CCCCGTCAAGCTCTAAATCGGGGGCTCCCTTTAGGGTTCCGATTTAGTGCTTTACGGCACCT
CGACCCCAAAAAACTTGATTAGGGTGATGGTTCACGTAGTGGGCCATCGCCCTGATAGACGG
TTTTTCGCCCTTTGACGTTGGAGTCCACGTTCTTTAATAGTGGACTCTTGTTCCAAACTGGA
ACAACACTCAACCCTATCTCGGTCTATTCTTTTGATTTATAAGGGATTTTGCCGATTTCGGC
CTATTGGTTAAAAAATGAGCTGATTTAACAAAAATTTAACGCGAATTTTAACAAAATATTAA
CGCTTACAATTTCAGGTGGCACTTTTCGGGGAAATGTGCGCGGAACCCCTATTTGTTTATTT
TTCTAAATACATTCAAATATGTATCCGCTCATGAGACAATAACCCTGATAAATGCTTCAATA
ATATTGAAAAAGGAAGAGTATGAGTATTCAACATTTCCGTGTCGCCCTTATTCCCTTTTTTG
CGGCATTTTGCCTTCCTGTTTTTGCTCACCCAGAAACGCTGGTGAAAGTAAAAGATGCTGAA
GATCAGTTGGGTGCACGAGTGGGTTACATCGAACTGGATCTCAACAGCGGTAAGATCCTTGA
GAGTTTTCGCCCCGAAGAACGTTTTCCAATGATGAGCACTTTTAAAGTTCTGCTATGTGGCG
CGGTATTATCCCGTATTGACGCCGGGCAAGAGCAACTCGGTCGCCGCATACACTATTCTCAG
AATGACTTGGTTGAGTACTCACCAGTCACAGAAAAGCATCTTACGGATGGCATGACAGTAAG
AGAATTATGCAGTGCTGCCATAACCATGAGTGATAACACTGCGGCCAACTTACTTCTGACAA
CGATCGGAGGACCGAAGGAGCTAACCGCTTTTTTGCACAACATGGGGGATCATGTAACTCGC
CTTGATCGTTGGGAACCGGAGCTGAATGAAGCCATACCAAACGACGAGCGTGACACCACGAT
GCCTGTAGCAATGGCAACAACGTTGCGCAAACTATTAACTGGCGAACTACTTACTCTAGCTT
CCCGGCAACAATTGATAGACTGGATGGAGGCGGATAAAGTTGCAGGACCACTTCTGCGCTCG
GCCCTTCCGGCTGGCTGGTTTATTGCTGATAAATCTGGAGCCGGTGAGCGTGGCTCTCGCGG
TATCATTGCAGCACTGGGGCCAGATGGTAAGCCCTCCCGTATCGTAGTTATCTACACGACGG
GGAGTCAGGCAACTATGGATGAACGAAATAGACAGATCGCTGAGATAGGTGCCTCACTGATT
AAGCATTGGTAGGAATTAATGATGTCTCGTTTAGATAAAAGTAAAGTGATTAACAGCGCATT
AGAGCTGCTTAATGAGGTCGGAATCGAAGGTTTAACAACCCGTAAACTCGCCCAGAAGCTAG
GTGTAGAGCAGCCTACATTGTATTGGCATGTAAAAAATAAGCGGGCTTTGCTCGACGCCTTA
GCCATTGAGATGTTAGATAGGCACCATACTCACTTTTGCCCTTTAGAAGGGGAAAG

>pASK-IBA5plus_P91-B (library, round 2)
CTGGCAAGATTTTTTACGTAATAACGCTAAAAGTTTTAGATGTGCTTTACTAAGTCATCGCG
ATGGAGCAAAAGTACATTTAGGTACACGGCCTACAGAAAAACAGTATGAAACTCTCGAAAAT
CAATTAGCCTTTTTATGCCAACAAGGTTTTTCACTAGAGAATGCATTATATGCACTCAGCGC
AGTGGGGCATTTTACTTTAGGTTGCGTATTGGAAGATCAAGAGCATCAAGTCGCTAAAGAAG
AAAGGGAAACACCTACTACTGATAGTATGCCGCCATTATTACGACAAGCTATCGAATTATTT
GATCACCAAGGTGCAGAGCCAGCCTTCTTATTCGGCCTTGAATTGATCATATGCGGATTAGA
AAAACAACTTAAATGTGAAAGTGGGTCTTAAAAGCAGCATAACCTTTTTCCGTGATGGTAAC
TTCACTAGTTTAAAAGGATCTAGGTGAAGATCCTTTTTGATAATCTCATGACCAAAATCCCT

TAACGTGAGTTTTCGTTCCACTGAGCGTCAGACCCCGTAGAAAAGATCAAAGGATCTTCTTG
AGATCCTTTTTTTCTGCGCGTAATCTGCTGCTTGCAAACAAAAAAACCACCGCTACCAGCGG
TGGTTTGTTTGCCGGATCAAGAGCTACCAACTCTTTTTCCGAAGGTAACTGGCTTCAGCAGA
GCGCAGATACCAAATACTGTCCTTCTAGTGTAGCCGTAGTTAGGCCACCACTTCAAGAACTC
TGTAGCACCGCCTACATACCTCGCTCTGCTAATCCTGTTACCAGTGGCTGCTGCCAGTGGCG
ATAAGTCGTGTCTTACCGGGTTGGACTCAAGACGATAGTTACCGGATAAGGCGCAGCGGTCG
GGCTGAACGGGGGGTTCGTGCACACAGCCCAGCTTGGAGCGAACGACCTACACCGAACTGAG
ATACCTACAGCGTGAGCTATGAGAAAGCGCCACGCTTCCCGAAGGGAGAAAGGCGGACAGGT
ATCCGGTAAGCGGCAGGGTCGGAACAGGAGAGCGCACGAGGGAGCTTCCAGGGGGAAACGCC
TGGTATCTTTATAGTCCTGTCGGGTTTCGCCACCTCTGACTTGAGCGTCGATTTTTGTGATG
CTCGTCAGGGGGGCGGAGCCTATGGAAAAACGCCAGCAACGCGGCCTTTTTACGGTTCCTGG
CCTTTTGCTGGCCTTTTGCTCACATGACCCGACACCATCGAATGGCCAGATGATTAATTCCT
AATTTTTGTTGACACTCTATCATTGATAGAGTTATTTTACCACTCCCTATCAGTGATAGAGA
AAAGTGAAATGAATAGTTCGACAAAAATCTAGAAATAATTTTGTTTAACTTTAAGAAGGAGA
TATACAAATGGCTAGCTGGAGCCACCCGCAGTTCGAAAAAGGCGCCATGACAGCAAGAAAAG
TCGACTACACAGACGGTGCAACCCGCTGTATCGGTGAGTTTCATTGGGATGAAGGCAAGTCG
GGCCCGCGTCCCGGCGTGGTGGTCTTTCCCGAG**XXX**TTCGGCCTCAACGACCATGCCAAGGA
GCGCGCGCGGCGCCTTGCCGACCTCGGCTTTGCAGCCCTGGCGGCGGATATGCACGGAGACG
CCCAGGTTTTCGAT**XXX**GCGAGT**XXX**TCATCAACCATACAGGGCTACTACGGCGACCGCGCC
CACTGGCGACGTCGTGCGCAGGCAGCGCTCGATGCACTGACGGCACAGCCAGAGGTGGACGG
CAGCAAGGTGGCGGCCATCGGCTTTTGTTTCGGCGGT**XXX**ACCTGCCTTGAACTGGCCCGCA
CAGGTGCGCCGCTGACCGCCATTGTCACCTTCCACGGCGGTTTGCTGCCGGAGATGGCAGGC
GATGCCGGACGGATCCAGTCCAGTGTTCTGGTGTGCCATGGCGCTGATGATCCGCTCGTACA
GGACGAAACCATGAAGGCCGTCATGGACGAGTTTCGTCGCGACAAGGTGGATTGGCAGGTGC
TCTACCTCGGAAATGCGGTACACAGTTTCACCGATCCACTCGCTGGCAGTCACGGC**TGG**CCC
GGG**GTT**GCCTATGACGCCACTGCCGAAGCCCGGTCGTGGACGGCCATGTGCAATCTGTTCAG
TGAACTGTTCGGCTGATGATATCTAACTAAGCTTGACCTGTGAAGTGAAAAATGGCGCACAT
TGTGCGACATTTTTTTTGTCTGCCGTTTACCGCTACTGCGTCACGGATCTCCACGCGCCCTG
TAGCGGCGCATTAAGCGCGGCGGGTGTGGTGGTTACGCGCAGCGTGACCGCTACACTTGCCA
GCGCCCTAGCGCCCGCTCCTTTCGCTTTCTTCCCTTCCTTTCTCGCCACGTTCGCCGGCTTT
CCCCGTCAAGCTCTAAATCGGGGGCTCCCTTTAGGGTTCCGATTTAGTGCTTTACGGCACCT
CGACCCCAAAAAACTTGATTAGGGTGATGGTTCACGTAGTGGGCCATCGCCCTGATAGACGG
TTTTTCGCCCTTTGACGTTGGAGTCCACGTTCTTTAATAGTGGACTCTTGTTCCAAACTGGA
ACAACACTCAACCCTATCTCGGTCTATTCTTTTGATTTATAAGGGATTTTGCCGATTTCGGC
CTATTGGTTAAAAAATGAGCTGATTTAACAAAAATTTAACGCGAATTTTAACAAAATATTAA
CGCTTACAATTTCAGGTGGCACTTTTCGGGGAAATGTGCGCGGAACCCCTATTTGTTTATTT
TTCTAAATACATTCAAATATGTATCCGCTCATGAGACAATAACCCTGATAAATGCTTCAATA
ATATTGAAAAAGGAAGAGTATGAGTATTCAACATTTCCGTGTCGCCCTTATTCCCTTTTTTG
CGGCATTTTGCCTTCCTGTTTTTGCTCACCCAGAAACGCTGGTGAAAGTAAAAGATGCTGAA
GATCAGTTGGGTGCACGAGTGGGTTACATCGAACTGGATCTCAACAGCGGTAAGATCCTTGA
GAGTTTTCGCCCCGAAGAACGTTTTCCAATGATGAGCACTTTTAAAGTTCTGCTATGTGGCG
CGGTATTATCCCGTATTGACGCCGGGCAAGAGCAACTCGGTCGCCGCATACACTATTCTCAG
AATGACTTGGTTGAGTACTCACCAGTCACAGAAAAGCATCTTACGGATGGCATGACAGTAAG
AGAATTATGCAGTGCTGCCATAACCATGAGTGATAACACTGCGGCCAACTTACTTCTGACAA
CGATCGGAGGACCGAAGGAGCTAACCGCTTTTTTGCACAACATGGGGGATCATGTAACTCGC
CTTGATCGTTGGGAACCGGAGCTGAATGAAGCCATACCAAACGACGAGCGTGACACCACGAT
GCCTGTAGCAATGGCAACAACGTTGCGCAAACTATTAACTGGCGAACTACTTACTCTAGCTT
CCCGGCAACAATTGATAGACTGGATGGAGGCGGATAAAGTTGCAGGACCACTTCTGCGCTCG
GCCCTTCCGGCTGGCTGGTTTATTGCTGATAAATCTGGAGCCGGTGAGCGTGGCTCTCGCGG
TATCATTGCAGCACTGGGGCCAGATGGTAAGCCCTCCCGTATCGTAGTTATCTACACGACGG
GGAGTCAGGCAACTATGGATGAACGAAATAGACAGATCGCTGAGATAGGTGCCTCACTGATT
AAGCATTGGTAGGAATTAATGATGTCTCGTTTAGATAAAAGTAAAGTGATTAACAGCGCATT

AGAGCTGCTTAATGAGGTCGGAATCGAAGGTTTAACAACCCGTAAACTCGCCCAGAAGCTAG
GTGTAGAGCAGCCTACATTGTATTGGCATGTAAAAAATAAGCGGGCTTTGCTCGACGCCTTA
GCCATTGAGATGTTAGATAGGCACCATACTCACTTTTGCCCTTTAGAAGGGGAAAG

>pASK-IBA5plus_6xHis-P91-WT
CTGGCAAGATTTTTTACGTAATAACGCTAAAAGTTTTAGATGTGCTTTACTAAGTCATCGCG
ATGGAGCAAAAGTACATTTAGGTACACGGCCTACAGAAAAACAGTATGAAACTCTCGAAAAT
CAATTAGCCTTTTTATGCCAACAAGGTTTTTCACTAGAGAATGCATTATATGCACTCAGCGC
AGTGGGGCATTTTACTTTAGGTTGCGTATTGGAAGATCAAGAGCATCAAGTCGCTAAAGAAG
AAAGGGAAACACCTACTACTGATAGTATGCCGCCATTATTACGACAAGCTATCGAATTATTT
GATCACCAAGGTGCAGAGCCAGCCTTCTTATTCGGCCTTGAATTGATCATATGCGGATTAGA
AAAACAACTTAAATGTGAAAGTGGGTCTTAAAAGCAGCATAACCTTTTTCCGTGATGGTAAC
TTCACTAGTTTAAAAGGATCTAGGTGAAGATCCTTTTTGATAATCTCATGACCAAAATCCCT
TAACGTGAGTTTTCGTTCCACTGAGCGTCAGACCCCGTAGAAAAGATCAAAGGATCTTCTTG
AGATCCTTTTTTTCTGCGCGTAATCTGCTGCTTGCAAACAAAAAAACCACCGCTACCAGCGG
TGGTTTGTTTGCCGGATCAAGAGCTACCAACTCTTTTTCCGAAGGTAACTGGCTTCAGCAGA
GCGCAGATACCAAATACTGTCCTTCTAGTGTAGCCGTAGTTAGGCCACCACTTCAAGAACTC
TGTAGCACCGCCTACATACCTCGCTCTGCTAATCCTGTTACCAGTGGCTGCTGCCAGTGGCG
ATAAGTCGTGTCTTACCGGGTTGGACTCAAGACGATAGTTACCGGATAAGGCGCAGCGGTCG
GGCTGAACGGGGGGTTCGTGCACACAGCCCAGCTTGGAGCGAACGACCTACACCGAACTGAG
ATACCTACAGCGTGAGCTATGAGAAAGCGCCACGCTTCCCGAAGGGAGAAAGGCGGACAGGT
ATCCGGTAAGCGGCAGGGTCGGAACAGGAGAGCGCACGAGGGAGCTTCCAGGGGGAAACGCC
TGGTATCTTTATAGTCCTGTCGGGTTTCGCCACCTCTGACTTGAGCGTCGATTTTTGTGATG
CTCGTCAGGGGGGCGGAGCCTATGGAAAAACGCCAGCAACGCGGCCTTTTTACGGTTCCTGG
CCTTTTGCTGGCCTTTTGCTCACATGACCCGACACCATCGAATGGCCAGATGATTAATTCCT
AATTTTTGTTGACACTCTATCATTGATAGAGTTATTTTACCACTCCCTATCAGTGATAGAGA
AAAGTGAAATGAATAGTTCGACAAAAATCTAGAAATAATTTTGTTTAACTTTAAGAAGGAGA
TATACAAATGCATCACCATCATCACCACGGTGGAAGTATGACAGCAAGAAAAGTCGACTACA
CAGACGGTGCAACCCGCTGTATCGGTGAGTTTCATTGGGATGAAGGCAAGTCGGGCCCGCGT
CCCGGCGTGGTGGTCTTTCCCGAGGCTTTCGGCCTCAACGACCATGCCAAGGAGCGCGCGCG
GCGCCTTGCCGACCTCGGCTTTGCAGCCCTGGCGGCGGATATGCACGGAGACGCCCAGGTTT
TCGATGCGGCGAGTCTCTCATCAACCATACAGGGCTACTACGGCGACCGCGCCCACTGGCGA
CGTCGTGCGCAGGCAGCGCTCGATGCACTGACGGCACAGCCAGAGGTGGACGGCAGCAAGGT
GGCGGCCATCGGCTTTTGTTTCGGCGGTGCCACCTGCCTTGAACTGGCCCGCACAGGTGCGC
CGCTGACCGCCATTGTCACCTTCCACGGCGGTTTGCTGCCGGAGATGGCAGGCGATGCCGGA
CGGATCCAGTCCAGTGTTCTGGTGTGCCATGGCGCTGATGATCCGCTCGTACAGGACGAAAC
CATGAAGGCCGTCATGGACGAGTTTCGTCGCGACAAGGTGGATTGGCAGGTGCTCTACCTCG
GAAATGCGGTACACAGTTTCACCGATCCACTCGCTGGCAGTCACGGCATACCCGGGCTGGCC
TATGACGCCACTGCCGAAGCCCGGTCGTGGACGGCCATGTGCAATCTGTTCAGTGAACTGTT
CGGCTGATGATATCTAACTAAGCTTGACCTGTGAAGTGAAAAATGGCGCACATTGTGCGACA
TTTTTTTTGTCTGCCGTTTACCGCTACTGCGTCACGGATCTCCACGCGCCCTGTAGCGGCGC
ATTAAGCGCGGCGGGTGTGGTGGTTACGCGCAGCGTGACCGCTACACTTGCCAGCGCCCTAG
CGCCCGCTCCTTTCGCTTTCTTCCCTTCCTTTCTCGCCACGTTCGCCGGCTTTCCCCGTCAA
GCTCTAAATCGGGGGCTCCCTTTAGGGTTCCGATTTAGTGCTTTACGGCACCTCGACCCCAA
AAAACTTGATTAGGGTGATGGTTCACGTAGTGGGCCATCGCCCTGATAGACGGTTTTTCGCC
CTTTGACGTTGGAGTCCACGTTCTTTAATAGTGGACTCTTGTTCCAAACTGGAACAACACTC
AACCCTATCTCGGTCTATTCTTTTGATTTATAAGGGATTTTGCCGATTTCGGCCTATTGGTT
AAAAAATGAGCTGATTTAACAAAAATTTAACGCGAATTTTAACAAAATATTAACGCTTACAA
TTTCAGGTGGCACTTTTCGGGGAAATGTGCGCGGAACCCCTATTTGTTTATTTTTCTAAATA
CATTCAAATATGTATCCGCTCATGAGACAATAACCCTGATAAATGCTTCAATAATATTGAAA
AAGGAAGAGTATGAGTATTCAACATTTCCGTGTCGCCCTTATTCCCTTTTTTGCGGCATTTT
GCCTTCCTGTTTTTGCTCACCCAGAAACGCTGGTGAAAGTAAAAGATGCTGAAGATCAGTTG

GGTGCACGAGTGGGTTACATCGAACTGGATCTCAACAGCGGTAAGATCCTTGAGAGTTTTCG
CCCCGAAGAACGTTTTCCAATGATGAGCACTTTTAAAGTTCTGCTATGTGGCGCGGTATTAT
CCCGTATTGACGCCGGGCAAGAGCAACTCGGTCGCCGCATACACTATTCTCAGAATGACTTG
GTTGAGTACTCACCAGTCACAGAAAAGCATCTTACGGATGGCATGACAGTAAGAGAATTATG
CAGTGCTGCCATAACCATGAGTGATAACACTGCGGCCAACTTACTTCTGACAACGATCGGAG
GACCGAAGGAGCTAACCGCTTTTTTGCACAACATGGGGGATCATGTAACTCGCCTTGATCGT
TGGGAACCGGAGCTGAATGAAGCCATACCAAACGACGAGCGTGACACCACGATGCCTGTAGC
AATGGCAACAACGTTGCGCAAACTATTAACTGGCGAACTACTTACTCTAGCTTCCCGGCAAC
AATTGATAGACTGGATGGAGGCGGATAAAGTTGCAGGACCACTTCTGCGCTCGGCCCTTCCG
GCTGGCTGGTTTATTGCTGATAAATCTGGAGCCGGTGAGCGTGGCTCTCGCGGTATCATTGC
AGCACTGGGGCCAGATGGTAAGCCCTCCCGTATCGTAGTTATCTACACGACGGGGAGTCAGG
CAACTATGGATGAACGAAATAGACAGATCGCTGAGATAGGTGCCTCACTGATTAAGCATTGG
TAGGAATTAATGATGTCTCGTTTAGATAAAAGTAAAGTGATTAACAGCGCATTAGAGCTGCT
TAATGAGGTCGGAATCGAAGGTTTAACAACCCGTAAACTCGCCCAGAAGCTAGGTGTAGAGC
AGCCTACATTGTATTGGCATGTAAAAAATAAGCGGGCTTTGCTCGACGCCTTAGCCATTGAG
ATGTTAGATAGGCACCATACTCACTTTTGCCCTTTAGAAGGGGAAAG

> pASK-IBA5plus_6xHis-P91-R1
CTGGCAAGATTTTTTACGTAATAACGCTAAAAGTTTTAGATGTGCTTTACTAAGTCATCGCG
ATGGAGCAAAAGTACATTTAGGTACACGGCCTACAGAAAAACAGTATGAAACTCTCGAAAAT
CAATTAGCCTTTTTATGCCAACAAGGTTTTTCACTAGAGAATGCATTATATGCACTCAGCGC
AGTGGGGCATTTTACTTTAGGTTGCGTATTGGAAGATCAAGAGCATCAAGTCGCTAAAGAAG
AAAGGGAAACACCTACTACTGATAGTATGCCGCCATTATTACGACAAGCTATCGAATTATTT
GATCACCAAGGTGCAGAGCCAGCCTTCTTATTCGGCCTTGAATTGATCATATGCGGATTAGA
AAAACAACTTAAATGTGAAAGTGGGTCTTAAAAGCAGCATAACCTTTTTCCGTGATGGTAAC
TTCACTAGTTTAAAAGGATCTAGGTGAAGATCCTTTTTGATAATCTCATGACCAAAATCCCT
TAACGTGAGTTTTCGTTCCACTGAGCGTCAGACCCCGTAGAAAAGATCAAAGGATCTTCTTG
AGATCCTTTTTTTCTGCGCGTAATCTGCTGCTTGCAAACAAAAAAACCACCGCTACCAGCGG
TGGTTTGTTTGCCGGATCAAGAGCTACCAACTCTTTTTCCGAAGGTAACTGGCTTCAGCAGA
GCGCAGATACCAAATACTGTCCTTCTAGTGTAGCCGTAGTTAGGCCACCACTTCAAGAACTC
TGTAGCACCGCCTACATACCTCGCTCTGCTAATCCTGTTACCAGTGGCTGCTGCCAGTGGCG
ATAAGTCGTGTCTTACCGGGTTGGACTCAAGACGATAGTTACCGGATAAGGCGCAGCGGTCG
GGCTGAACGGGGGGTTCGTGCACACAGCCCAGCTTGGAGCGAACGACCTACACCGAACTGAG
ATACCTACAGCGTGAGCTATGAGAAAGCGCCACGCTTCCCGAAGGGAGAAAGGCGGACAGGT
ATCCGGTAAGCGGCAGGGTCGGAACAGGAGAGCGCACGAGGGAGCTTCCAGGGGGAAACGCC
TGGTATCTTTATAGTCCTGTCGGGTTTCGCCACCTCTGACTTGAGCGTCGATTTTTGTGATG
CTCGTCAGGGGGGCGGAGCCTATGGAAAAACGCCAGCAACGCGGCCTTTTTACGGTTCCTGG
CCTTTTGCTGGCCTTTTGCTCACATGACCCGACACCATCGAATGGCCAGATGATTAATTCCT
AATTTTTGTTGACACTCTATCATTGATAGAGTTATTTTACCACTCCCTATCAGTGATAGAGA
AAAGTGAAATGAATAGTTCGACAAAAATCTAGAAATAATTTTGTTTAACTTTAAGAAGGAGA
TATACAAATGCATCACCATCATCACCACGGTGGAAGTATGACAGCAAGAAAAGTCGACTACA
CAGACGGTGCAACCCGCTGTATCGGTGAGTTTCATTGGATGAAGGCAAGTCGGCCCGCGT
CCCGGCGTGGTGGTCTTTCCCGAGGCTTTCGGCCTCAACGACCATGCCAAGGAGCGCGCGCG
GCGCCTTGCCGACCTCGGCTTTGCAGCCCTGGCGGCGGATATGCACGGAGACGCCCAGGTTT
TCGATGCGGCGAGTCTCTCATCAACCATACAGGGCTACTACGGCGACCGCGCCCACTGGCGA
CGTCGTGCGCAGGCAGCGCTCGATGCACTGACGGCACAGCCAGAGGTGGACGGCAGCAAGGT
GGCGGCCATCGGCTTTTGTTTCGGCGGTGCCACCTGCCTTGAACTGGCCCGCACAGGTGCGC
CGCTGACCGCCATTGTCACCTTCCACGGCGGTTTGCTGCCGGAGATGGCAGGCGATGCCGGA
CGGATCCAGTCCAGTGTTCTGGTGTGCCATGGCGCTGATGATCCGCTCGTACAGGACGAAAC
CATGAAGGCCGTCATGGACGAGTTTCGTCGCGACAAGGTGGATTGGCAGGTGCTCTACCTCG
GAAATGCGGTACACAGTTTCACCGATCCACTCGCTGGCAGTCACGGC**TGG**CCCGGG**GTT**GCC

TATGACGCCACTGCCGAAGCCCGGTCGTGGACGGCCATGTGCAATCTGTTCAGTGAACTGTT
CGGCTGATGATATCTAACTAAGCTTGACCTGTGAAGTGAAAAATGGCGCACATTGTGCGACA
TTTTTTTTGTCTGCCGTTTACCGCTACTGCGTCACGGATCTCCACGCGCCCTGTAGCGGCGC
ATTAAGCGCGGCGGGTGTGGTGGTTACGCGCAGCGTGACCGCTACACTTGCCAGCGCCCTAG
CGCCCGCTCCTTTCGCTTTCTTCCCTTCCTTTCTCGCCACGTTCGCCGGCTTTCCCCGTCAA
GCTCTAAATCGGGGGCTCCCTTTAGGGTTCCGATTTAGTGCTTTACGGCACCTCGACCCCAA
AAAACTTGATTAGGGTGATGGTTCACGTAGTGGGCCATCGCCCTGATAGACGGTTTTTCGCC
CTTTGACGTTGGAGTCCACGTTCTTTAATAGTGGACTCTTGTTCCAAACTGGAACAACACTC
AACCCTATCTCGGTCTATTCTTTTGATTTATAAGGGATTTTGCCGATTTCGGCCTATTGGTT
AAAAAATGAGCTGATTTAACAAAAATTTAACGCGAATTTTAACAAAATATTAACGCTTACAA
TTTCAGGTGGCACTTTTCGGGGAAATGTGCGCGGAACCCCTATTTGTTTATTTTTCTAAATA
CATTCAAATATGTATCCGCTCATGAGACAATAACCCTGATAAATGCTTCAATAATATTGAAA
AAGGAAGAGTATGAGTATTCAACATTTCCGTGTCGCCCTTATTCCCTTTTTTGCGGCATTTT
GCCTTCCTGTTTTTGCTCACCCAGAAACGCTGGTGAAAGTAAAAGATGCTGAAGATCAGTTG
GGTGCACGAGTGGGTTACATCGAACTGGATCTCAACAGCGGTAAGATCCTTGAGAGTTTTCG
CCCCGAAGAACGTTTTCCAATGATGAGCACTTTTAAAGTTCTGCTATGTGGCGCGGTATTAT
CCCGTATTGACGCCGGGCAAGAGCAACTCGGTCGCCGCATACACTATTCTCAGAATGACTTG
GTTGAGTACTCACCAGTCACAGAAAAGCATCTTACGGATGGCATGACAGTAAGAGAATTATG
CAGTGCTGCCATAACCATGAGTGATAACACTGCGGCCAACTTACTTCTGACAACGATCGGAG
GACCGAAGGAGCTAACCGCTTTTTTGCACAACATGGGGGATCATGTAACTCGCCTTGATCGT
TGGGAACCGGAGCTGAATGAAGCCATACCAAACGACGAGCGTGACACCACGATGCCTGTAGC
AATGGCAACAACGTTGCGCAAACTATTAACTGGCGAACTACTTACTCTAGCTTCCCGGCAAC
AATTGATAGACTGGATGGAGGCGGATAAAGTTGCAGGACCACTTCTGCGCTCGGCCCTTCCG
GCTGGCTGGTTTATTGCTGATAAATCTGGAGCCGGTGAGCGTGGCTCTCGCGGTATCATTGC
AGCACTGGGGCCAGATGGTAAGCCCTCCCGTATCGTAGTTATCTACACGACGGGGAGTCAGG
CAACTATGGATGAACGAAATAGACAGATCGCTGAGATAGGTGCCTCACTGATTAAGCATTGG
TAGGAATTAATGATGTCTCGTTTAGATAAAAGTAAAGTGATTAACAGCGCATTAGAGCTGCT
TAATGAGGTCGGAATCGAAGGTTTAACAACCCGTAAACTCGCCCAGAAGCTAGGTGTAGAGC
AGCCTACATTGTATTGGCATGTAAAAAATAAGCGGGCTTTGCTCGACGCCTTAGCCATTGAG
ATGTTAGATAGGCACCATACTCACTTTTGCCCTTTAGAAGGGGAAAG

> pASK-IBA5plus_6xHis-P91-R2
CTGGCAAGATTTTTTACGTAATAACGCTAAAAGTTTTAGATGTGCTTTACTAAGTCATCGCG
ATGGAGCAAAAGTACATTTAGGTACACGGCCTACAGAAAAACAGTATGAAACTCTCGAAAAT
CAATTAGCCTTTTTATGCCAACAAGGTTTTTCACTAGAGAATGCATTATATGCACTCAGCGC
AGTGGGGCATTTTACTTTAGGTTGCGTATTGGAAGATCAAGAGCATCAAGTCGCTAAAGAAG
AAAGGGAAACACCTACTACTGATAGTATGCCGCCATTATTACGACAAGCTATCGAATTATTT
GATCACCAAGGTGCAGAGCCAGCCTTCTTATTCGGCCTTGAATTGATCATATGCGGATTAGA
AAAACAACTTAAATGTGAAAGTGGGTCTTAAAAGCAGCATAACCTTTTTCCGTGATGGTAAC
TTCACTAGTTTAAAAGGATCTAGGTGAAGATCCTTTTTGATAATCTCATGACCAAAATCCCT
TAACGTGAGTTTTCGTTCCACTGAGCGTCAGACCCCGTAGAAAAGATCAAAGGATCTTCTTG
AGATCCTTTTTTTCTGCGCGTAATCTGCTGCTTGCAAACAAAAAAACCACCGCTACCAGCGG
TGGTTTGTTTGCCGGATCAAGAGCTACCAACTCTTTTTCCGAAGGTAACTGGCTTCAGCAGA
GCGCAGATACCAAATACTGTCCTTCTAGTGTAGCCGTAGTTAGGCCACCACTTCAAGAACTC
TGTAGCACCGCCTACATACCTCGCTCTGCTAATCCTGTTACCAGTGGCTGCTGCCAGTGGCG
ATAAGTCGTGTCTTACCGGGTTGGACTCAAGACGATAGTTACCGGATAAGGCGCAGCGGTCG
GGCTGAACGGGGGGTTCGTGCACACAGCCCAGCTTGGAGCGAACGACCTACACCGAACTGAG
ATACCTACAGCGTGAGCTATGAGAAAGCGCCACGCTTCCCGAAGGGAGAAAGGCGGACAGGT
ATCCGGTAAGCGGCAGGGTCGGAACAGGAGAGCGCACGAGGGAGCTTCCAGGGGGAAACGCC
TGGTATCTTTATAGTCCTGTCGGGTTTCGCCACCTCTGACTTGAGCGTCGATTTTTGTGATG
CTCGTCAGGGGGGCGGAGCCTATGGAAAAACGCCAGCAACGCGGCCTTTTTACGGTTCCTGG
CCTTTTGCTGGCCTTTTGCTCACATGACCCGACACCATCGAATGGCCAGATGATTAATTCCT

AATTTTTGTTGACACTCTATCATTGATAGAGTTATTTTACCACTCCCTATCAGTGATAGAGA
AAAGTGAAATGAATAGTTCGACAAAAATCTAGAAATAATTTTGTTTAACTTTAAGAAGGAGA
TATACAAATGCATCACCATCATCACCACGGTGGAAGTATGACAGCAAGAAAAGTCGACTACA
CAGACGGTGCAACCCGCTGTATCGGTGAGTTTCATTGGGATGAAGGCAAGTCGGGCCCGCGT
CCCGGCGTGGTGGTCTTTCCCGAG**CTG**TTCGGCCTCAACGACCATGCCAAGGAGCGCGCGCG
GCGCCTTGCCGACCTCGGCTTTGCAGCCCTGGCGGCGGATATGCACGGAGACGCCCAGGTTT
TCGAT**GAG**GCGAGT**GTG**TCATCAACCATACAGGGCTACTACGGCGACCGCGCCCACTGGCGA
CGTCGTGCGCAGGCAGCGCTCGATGCACTGACGGCACAGCCAGAGGTGGACGGCAGCAAGGT
GGCGGCCATCGGCTTTTGTTTCGGCGGTGCGACCTGCCTTGAACTGGCCCGCACAGGTGCGC
CGCTGACCGCCATTGTCACCTTCCACGGCGGTTTGCTGCCGGAGATGGCAGGCGATGCCGGA
CGGATCCAGTCCAGTGTTCTGGTGTGCCATGGCGCTGATGATCCGCTCGTACAGGACGAAAC
CATGAAGGCCGTCATGGACGAGTTTCGTCGCGACAAGGTGGATTGGCAGGTGCTCTACCTCG
GAAATGCGGTACACAGTTTCACCGATCCACTCGCTGGCAGTCACGGC**TGG**CCCGGG**GTT**GCC
TATGACGCCACTGCCGAAGCCCGGTCGTGGACGGCCATGTGCAATCTGTTCAGTGAACTGTT
CGGCTGATGATATCTAACTAAGCTTGACCTGTGAAGTGAAAAATGGCGCACATTGTGCGACA
TTTTTTTTGTCTGCCGTTTACCGCTACTGCGTCACGGATCTCCACGCGCCCTGTAGCGGCGC
ATTAAGCGCGGCGGGTGTGGTGGTTACGCGCAGCGTGACCGCTACACTTGCCAGCGCCCTAG
CGCCCGCTCCTTTCGCTTTCTTCCCTTCCTTTCTCGCCACGTTCGCCGGCTTTCCCCGTCAA
GCTCTAAATCGGGGGCTCCCTTTAGGGTTCCGATTTAGTGCTTTACGGCACCTCGACCCCAA
AAAACTTGATTAGGGTGATGGTTCACGTAGTGGGCCATCGCCCTGATAGACGGTTTTTCGCC
CTTTGACGTTGGAGTCCACGTTCTTTAATAGTGGACTCTTGTTCCAAACTGGAACAACACTC
AACCCTATCTCGGTCTATTCTTTTGATTTATAAGGGATTTTGCCGATTTCGGCCTATTGGTT
AAAAAATGAGCTGATTTAACAAAAATTTAACGCGAATTTTAACAAAATATTAACGCTTACAA
TTTCAGGTGGCACTTTTCGGGGAAATGTGCGCGGAACCCCTATTTGTTTATTTTTCTAAATA
CATTCAAATATGTATCCGCTCATGAGACAATAACCCTGATAAATGCTTCAATAATATTGAAA
AAGGAAGAGTATGAGTATTCAACATTTCCGTGTCGCCCTTATTCCCTTTTTTGCGGCATTTT
GCCTTCCTGTTTTTGCTCACCCAGAAACGCTGGTGAAAGTAAAAGATGCTGAAGATCAGTTG
GGTGCACGAGTGGGTTACATCGAACTGGATCTCAACAGCGGTAAGATCCTTGAGAGTTTTCG
CCCCGAAGAACGTTTTCCAATGATGAGCACTTTTAAAGTTCTGCTATGTGGCGCGGTATTAT
CCCGTATTGACGCCGGGCAAGAGCAACTCGGTCGCCGCATACACTATTCTCAGAATGACTTG
GTTGAGTACTCACCAGTCACAGAAAAGCATCTTACGGATGGCATGACAGTAAGAGAATTATG
CAGTGCTGCCATAACCATGAGTGATAACACTGCGGCCAACTTACTTCTGACAACGATCGGAG
GACCGAAGGAGCTAACCGCTTTTTTGCACAACATGGGGGATCATGTAACTCGCCTTGATCGT
TGGGAACCGGAGCTGAATGAAGCCATACCAAACGACGAGCGTGACACCACGATGCCTGTAGC
AATGGCAACAACGTTGCGCAAACTATTAACTGGCGAACTACTTACTCTAGCTTCCCGGCAAC
AATTGATAGACTGGATGGAGGCGGATAAAGTTGCAGGACCACTTCTGCGCTCGGCCCTTCCG
GCTGGCTGGTTTATTGCTGATAAATCTGGAGCCGGTGAGCGTGGCTCTCGCGGTATCATTGC
AGCACTGGGGCCAGATGGTAAGCCCTCCCGTATCGTAGTTATCTACACGACGGGGAGTCAGG
CAACTATGGATGAACGAAATAGACAGATCGCTGAGATAGGTGCCTCACTGATTAAGCATTGG
TAGGAATTAATGATGTCTCGTTTAGATAAAAGTAAAGTGATTAACAGCGCATTAGAGCTGCT
TAATGAGGTCGGAATCGAAGGTTTAACAACCCGTAAACTCGCCCAGAAGCTAGGTGTAGAGC
AGCCTACATTGTATTGGCATGTAAAAAATAAGCGGGCTTTGCTCGACGCCTTAGCCATTGAG
ATGTTAGATAGGCACCATACTCACTTTTGCCCTTTAGAAGGGGAAAG

#### 4.2 Primer sequences

Annealing parts are shown in uppercase, overhangs in lowercase. Mutagenic or degenerate codons are highlighted in bold.

##### Sequencing primers for pASK-IBA5+ constructs:

| Label | Sequence (5'→3') |
|---|---|
| IF | GAGTTATTTTACCACTCCCT |
| IR | CGCAGTAGCGGTAAACG |

##### Saturation mutagenesis for mutational scanning:

| Label | Sequence (5'→3') |
|---|---|
| P91 SM 37 NDT fwd | gattacaggtctcgTCCC**NDT**GCTTTCGGCCTCAACG |
| P91 SM 37 VHG fwd | gattacaggtctcgTCCC**VHG**GCTTTCGGCCTCAACG |
| P91 SM 37 TGG fwd | gattacaggtctcgTCCC**TGG**GCTTTCGGCCTCAACG |
| P91 SM 37 rev | gattacaggtctcgGGGAAAGACCACCACGC |
| P91 SM 38 NDT fwd | gattacaggtctcgCGAG**NDT**TTCGGCCTCAACGACC |
| P91 SM 38 VHG fwd | gattacaggtctcgCGAG**VHG**TTCGGCCTCAACGACC |
| P91 SM 38 TGG fwd | gattacaggtctcgCGAG**TGG**TTCGGCCTCAACGACC |
| P91 SM 38 rev | gattacaggtctcgCTCGGGAAAGACCACCAC |
| P91 SM 39 NDT fwd | gattacaggtctcgGGCT**NDT**GGCCTCAACGACCATGC |
| P91 SM 39 VHG fwd | gattacaggtctcgGGCT**VHG**GGCCTCAACGACCATGC |
| P91 SM 39 TGG fwd | gattacaggtctcgGGCT**TGG**GGCCTCAACGACCATGC |
| P91 SM 39 rev | gattacaggtctcgAGCCTCGGGAAAGACCAC |
| P91 SM 71 NDT fwd | ggtctcgGGTT**NDT**GATGCGGCGAGTCTCTC |
| P91 SM 71 VHG fwd | ggtctcgGGTT**VHG**GATGCGGCGAGTCTCTC |
| P91 SM 71 TGG fwd | ggtctcgGGTT**TGG**GATGCGGCGAGTCTCTC |
| P91 SM 71 rev | ggtctcgAACCTGGGCGTCTCCG |
| P91 SM 72 NDT fwd | ggtctcgTTTC**NDT**GCGGCGAGTCTCTCATC |
| P91 SM 72 VHG fwd | ggtctcgTTTC**VHG**GCGGCGAGTCTCTCATC |
| P91 SM 72 TGG fwd | ggtctcgTTTC**TGG**GCGGCGAGTCTCTCATC |
| P91 SM 72 rev | ggtctcgGAAAACCTGGGCGTCTCC |
| P91 SM 73 NDT fwd | ggtctcgCGAT**NDT**GCGAGTCTCTCATCAACCATAC |
| P91 SM 73 VHG fwd | ggtctcgCGAT**VHG**GCGAGTCTCTCATCAACCATAC |
| P91 SM 73 TGG fwd | ggtctcgCGAT**TGG**GCGAGTCTCTCATCAACCATAC |
| P91 SM 71-74 rev | ggtctcgATCGAAAACCTGGGCG |
| P91 SM 74 NDT fwd | ggtctcgTGCG**NDT**AGTCTCTCATCAACCATACAGGG |
| P91 SM 74 VHG fwd | ggtctcgTGCG**VHG**AGTCTCTCATCAACCATACAGGG |
| P91 SM 74 TGG fwd | ggtctcgTGCG**TGG**AGTCTCTCATCAACCATACAGGG |
| P91 SM 71-74 rev | ggtctcgCGCATCGAAAACCTGG |
| P91 SM 76 NDT fwd | gattacaggtctcgGAGT**NDT**TCATCAACCATACAGGGCTACTAC |
| P91 SM 76 VHG fwd | gattacaggtctcgGAGT**VHG**TCATCAACCATACAGGGCTACTAC |
| P91 SM 76 TGG fwd | gattacaggtctcgGAGT**TGG**TCATCAACCATACAGGGCTACTAC |
| P91 SM 76 rev | gattacaggtctccACTCGCCGCATCGAA |
| P91 SM 77 NDT fwd | gattacaggtctctTCTC**NDT**TCAACCATACAGGGCTACTACG |
| P91 SM 77 VHG fwd | gattacaggtctctTCTC**VHG**TCAACCATACAGGGCTACTACG |
| P91 SM 77 TGG fwd | gattacaggtctctTCTC**TGG**TCAACCATACAGGGCTACTACG |
| P91 SM 77 rev | gattacaggtctcgGAGACTCGCCGCATCG |
| P91 SM 80 NDT fwd | ggtctcgAACC**NDT**CAGGGCTACTACGGCGAC |
| P91 SM 80 VHG fwd | ggtctcgAACC**VHG**CAGGGCTACTACGGCGAC |
| P91 SM 80 TGG fwd | ggtctcgAACC**TGG**CAGGGCTACTACGGCGAC |
| P91 SM 80 rev | ggtctcgGGTTGATGAGAGACTCGCC |
| P91 SM 81 NDT fwd | gattacaggtctcgCATA**NDT**GGCTACTACGGCGACCG |
| P91 SM 81 VHG fwd | gattacaggtctcgCATA**VHG**GGCTACTACGGCGACCG |
| P91 SM 81 TGG fwd | gattacaggtctcgCATA**TGG**GGCTACTACGGCGACCG |
| P91 SM 81 rev | gattacaggtctccTATGGTTGATGAGAGACTCGCC |
| P91 SM 84 NDT fwd | gattacaggtctcaCTAC**NDT**GGCGACCGCGC |

| | |
|---|---|
| P91 SM 84 VHG fwd | gattacaggtctcaCTAC**VHG**GGCGACCGCGC |
| P91 SM 84 TGG fwd | gattacaggtctcaCTAC**TGG**GGCGACCGCGC |
| P91 SM 84 rev | gattacaggtctcgGTAGCCCTGTATGGTTGATGAG |
| P91 SM 119 NDT fwd | gattacaggtctcgTTGT**NDT**GGCGGTGCCACCT |
| P91 SM 119 VHG fwd | gattacaggtctcgTTGT**VHG**GGCGGTGCCACCT |
| P91 SM 119 TGG fwd | gattacaggtctcgTTGT**TGG**GGCGGTGCCACCT |
| P91 SM 119 rev | gattacaggtctcgACAAAAGCCGATGGCC |
| P91 SM 122 NDT fwd | gattacaggtctcaCGGT**NDT**ACCTGCCTTGAACTGGC |
| P91 SM 122 VHG fwd | gattacaggtctcaCGGT**VHG**ACCTGCCTTGAACTGGC |
| P91 SM 122 TGG fwd | gattacaggtctcaCGGT**TGG**ACCTGCCTTGAACTGGC |
| P91 SM 122 rev | gattacaggtctcgACCGCCGAAACAAAAGC |
| P91 SM 141 NDT fwd | gattacaggtctcgCTTC**NDT**GGCGGTTTGCTGCC |
| P91 SM 141 VHG fwd | gattacaggtctcgCTTC**VHG**GGCGGTTTGCTGCC |
| P91 SM 141 TGG fwd | gattacaggtctcgCTTC**TGG**GGCGGTTTGCTGCC |
| P91 SM 141 rev | gattacaggtctcaGAAGGTGACAATGGCGG |
| P91 SM 143 NDT fwd | gattacaggtctcgCGGC**NDT**TTGCTGCCGGAGATGG |
| P91 SM 143 VHG fwd | gattacaggtctcgCGGC**VHG**TTGCTGCCGGAGATGG |
| P91 SM 143 TGG fwd | gattacaggtctcgCGGC**TGG**TTGCTGCCGGAGATGG |
| P91 SM 143 rev | gattacaggtctcgGCCGTGGAAGGTGACAAT |
| P91 SM 167 NDT fwd | gattacaggtctcgTGAT**NDT**CCGCTCGTACAGGACG |
| P91 SM 167 VHG fwd | gattacaggtctcgTGAT**VHG**CCGCTCGTACAGGACG |
| P91 SM 167 TGG fwd | gattacaggtctcgTGAT**TGG**CCGCTCGTACAGGACG |
| P91 SM 167 rev | gattacaggtctcgATCAGCGCCATGGCA |
| P91 SM 169 NDT fwd | gattacaggtctcgTCCG**NDT**GTACAGGACGAAACCATGAAGG |
| P91 SM 169 VHG fwd | gattacaggtctcgTCCG**VHG**GTACAGGACGAAACCATGAAGG |
| P91 SM 169 TGG fwd | gattacaggtctcgTCCG**TGG**GTACAGGACGAAACCATGAAGG |
| P91 SM 169 rev | gattacaggtctcaCGGATCATCAGCGCC |
| P91 SM 197 NDT fwd | GGTCTCCAAAT**NDT**GTACACAGTTTCACCGATCCAC |
| P91 SM 197 VHG fwd | GGTCTCCAAAT**VHG**GTACACAGTTTCACCGATCCAC |
| P91 SM 197 TGG fwd | GGTCTCCAAAT**TGG**GTACACAGTTTCACCGATCCAC |
| P91 SM 197 rev | ggtctcgATTTCCGAGGTAGAGCACCTG |
| P91 SM 199 NDT fwd | gattacaggtctcgGGTA**NDT**AGTTTCACCGATCCACTCG |
| P91 SM 199 VHG fwd | gattacaggtctcgGGTA**VHG**AGTTTCACCGATCCACTCG |
| P91 SM 199 TGG fwd | gattacaggtctcgGGTA**TGG**AGTTTCACCGATCCACTCG |
| P91 SM 199 rev | GATTACAGGTCTCCTACCGCATTTCCGAGGTAG |
| P91 SM 200 NDT fwd | gattacaggtctcgACAC**NDT**TTCACCGATCCACTCGC |
| P91 SM 200 VHG fwd | gattacaggtctcgACAC**VHG**TTCACCGATCCACTCGC |
| P91 SM 200 TGG fwd | gattacaggtctcgACAC**TGG**TTCACCGATCCACTCGC |
| P91 SM 200 rev | gattacaggtctcgGTGTACCGCATTTCCGAG |
| P91 SM 211 NDT fwd | gattacaggtctcgCGGC**NDT**CCCGGGCTGGCC |
| P91 SM 211 VHG fwd | gattacaggtctcgCGGC**VHG**CCCGGGCTGGCC |
| P91 SM 211 TGG fwd | gattacaggtctcgCGGC**TGG**CCCGGGCTGGCC |
| P91 SM 211 rev | gattacaggtctcgGCCGTGACTGCCAGC |
| P91 SM 214 NDT fwd | gattacaggtctcgCGGG**NDT**GCCTATGACGCCACTGC |
| P91 SM 214 VHG fwd | gattacaggtctcgCGGG**VHG**GCCTATGACGCCACTGC |
| P91 SM 214 TGG fwd | gattacaggtctcgCGGG**TGG**GCCTATGACGCCACTGC |
| P91 SM 214 rev | gattacaggtctcaCCCGGGTATGCCGT |

#### Nucleophile exchange:

| Label | Sequence (5'→3') |
|---|---|
| P91 WT/R1 C118S fwd | gattacaggtctccCTTT**AGC**TTCGGCGGTGCCAC |
| P91 R2 C118S fwd | gattacaggtctccCTTT**AGC**TTCGGCGGTGCG |
| P91 C118S rev | gattacaggtctcgAAAGCCGATGGCCG |

#### Construction of library P91-A (round 1)

| Fragment | Label | Sequence (5'→3') |
|---|---|---|
| 1 | P91-A Frag 1 A73NNK fwd | ACGCCCAGGTTTTCGAT**NNK**GCGAGTCTCTCATCAACCATAC |

| | | |
|---|---|---|
| | P91-A Frag1 rev | ATCGAAAACCTGGGCGT |
| 2 | P91-A Frag2 211NNK 214NNK fwd | GCTGGCAGTCACGGC**NNK**CCCGGG**NNK**GCCTATGACGCCACTGC |
| | P91-A Frag2 rev | GCCGTGACTGCCAGC |

#### Construction of library P91-B (round 2)

| Fragment | Label | Sequence (5'→3') |
|---|---|---|
| 1 | P91 SM 38 NDT fwd | gattacaggtctcgCGAG**NDT**TTCGGCCTCAACGACC |
| | P91 SM 38 VHG fwd | gattacaggtctcgCGAG**VHG**TTCGGCCTCAACGACC |
| | P91 SM 71-74 rev | ggtctcgATCGAAAACCTGGGCG |
| | P91 SM 38 TGG fwd | gattacaggtctcgCGAG**TGG**TTCGGCCTCAACGACC |
| 2 | P91 A73NDT L76NDT fwd | gattacaggtctcgCGAT**NDT**GCGAGT**NDT**TCATCAACCATACAGGGCTACTAC |
| | P91 A73NDT L76VHG fwd | gattacaggtctcgCGAT**NDT**GCGAGT**VHG**TCATCAACCATACAGGGCTACTAC |
| | P91 A73NDT L76TGG fwd | gattacaggtctcgCGAT**NDT**GCGAGT**TGG**TCATCAACCATACAGGGCTACTAC |
| | P91 A73VHG L76NDT fwd | gattacaggtctcgCGAT**VHG**GCGAGT**NDT**TCATCAACCATACAGGGCTACTAC |
| | P91 A73VHG L76VHG fwd | gattacaggtctcgCGAT**VHG**GCGAGT**VHG**TCATCAACCATACAGGGCTACTAC |
| | P91 A73VHG L76TGG fwd | gattacaggtctcgCGAT**VHG**GCGAGT**TGG**TCATCAACCATACAGGGCTACTAC |
| | P91 A73TGG L76NDT fwd | gattacaggtctcgCGAT**TGG**GCGAGT**NDT**TCATCAACCATACAGGGCTACTAC |
| | P91 A73TGG L76VHG fwd | gattacaggtctcgCGAT**TGG**GCGAGT**VHG**TCATCAACCATACAGGGCTACTAC |
| | P91 A73TGG L76TGG fwd | gattacaggtctcgCGAT**TGG**GCGAGT**TGG**TCATCAACCATACAGGGCTACTAC |
| | P91 SM 122 rev | gattacaggtctcgACCGCCGAAACAAAAGC |
| 3 | P91 SM 122 NDT fwd | gattacaggtctcaCGGT**NDT**ACCTGCCTTGAACTGGC |
| | P91 SM 122 VHG fwd | gattacaggtctcaCGGT**VHG**ACCTGCCTTGAACTGGC |
| | P91 SM 122 TGG fwd | gattacaggtctcaCGGT**TGG**ACCTGCCTTGAACTGGC |
| | P91 SM 38 rev | gattacaggtctcgCTCGGAAAGACCACCAC |

#### Introduction of an N-terminal 6xHis-tag:

| Label | Sequence (5'→3') |
|---|---|
| pASK_strepless_rev | gattacaggtctcgTTGTATATCTCCTTCTTAAAGTTAAACAAAATTATTTCTAG |
| 6xHis_P91_fwd | gattacaggtctcgACAA**atgcatcaccatcatcaccacggtggaagt**ATGACAGCAAGAAAAGTCGAC |

### 5. NMR spectra

NMR data were collected at 298 K using Bruker Avance spectrometers with $^{1}H$ resonance frequencies of 400 MHz.

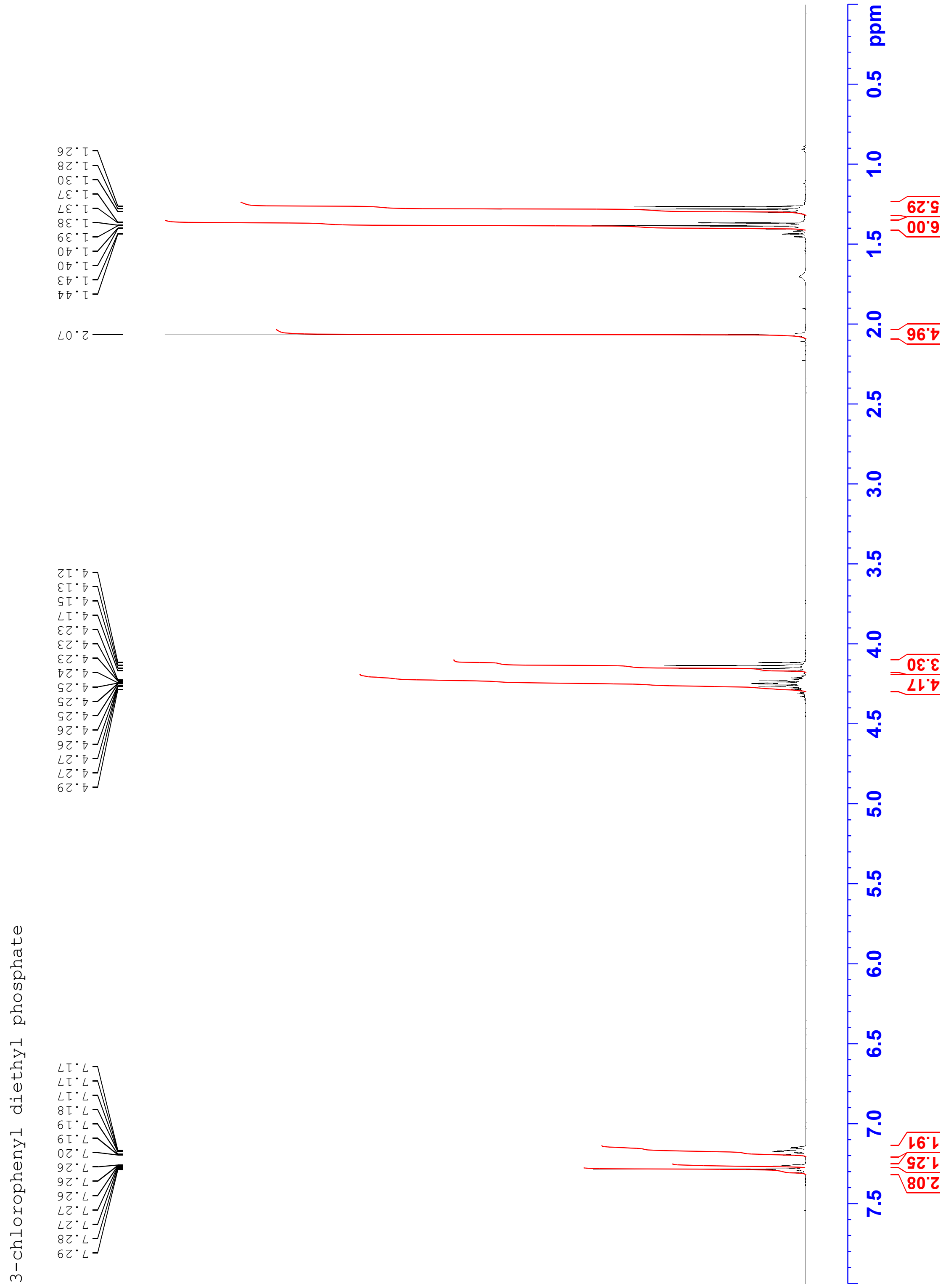

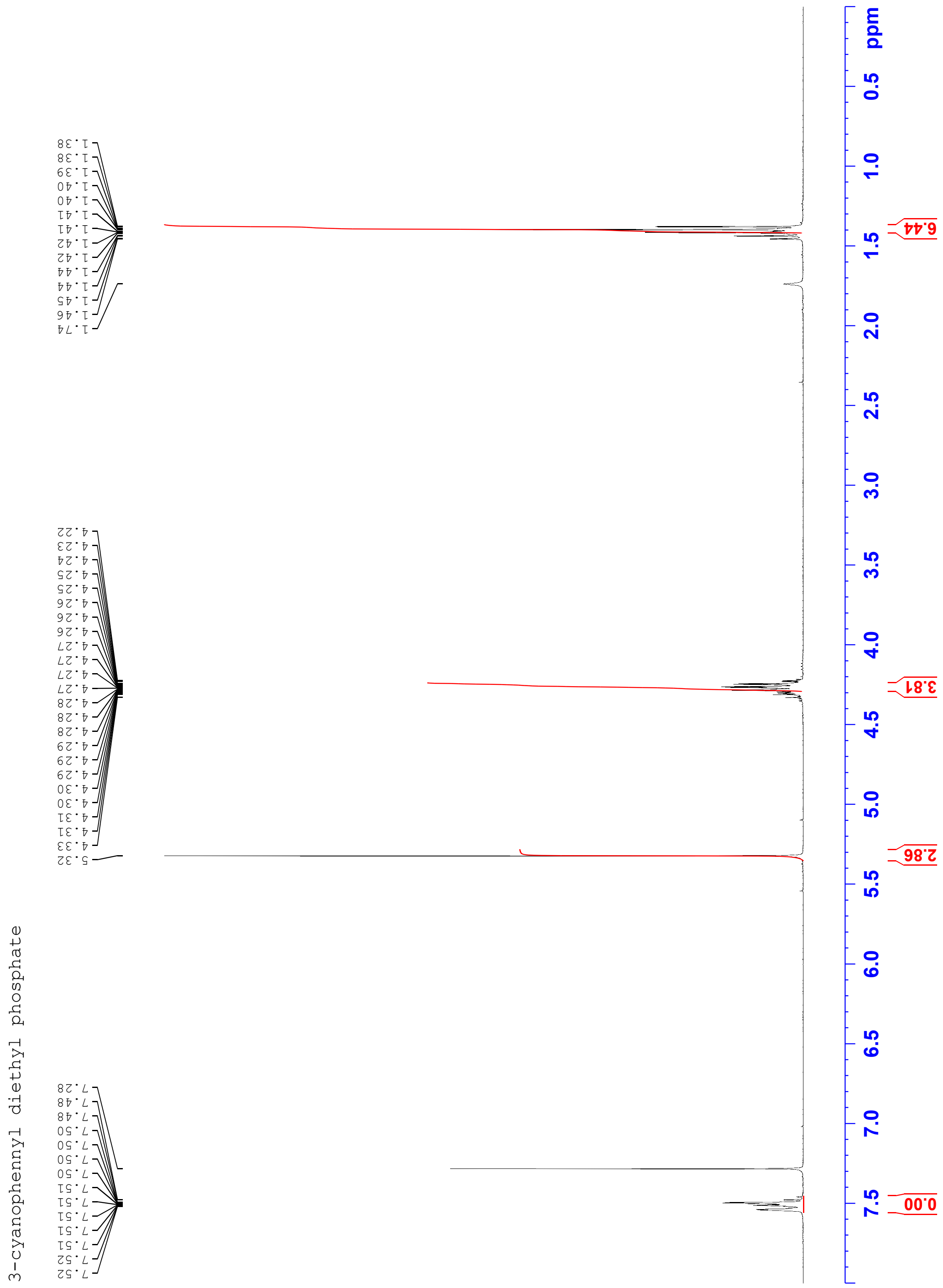
3-cyanophenyl diethyl phosphate
7.52
7.52
7.51
7.51
7.51
7.51
7.51
7.50
7.50
7.50
7.50
7.48
7.48
7.28
5.32
4.33
4.31
4.31
4.30
4.30
4.29
4.29
4.29
4.28
4.28
4.28
4.27
4.27
4.27
4.27
4.26
4.26
4.26
4.25
4.25
4.24
4.23
4.22
1.74
1.46
1.45
1.44
1.44
1.42
1.42
1.41
1.41
1.40
1.40
1.39
1.38
1.38
0.00
2.86
3.81
6.44
ppm

3-fluoro-4-nitrophenyl diethyl phosphate

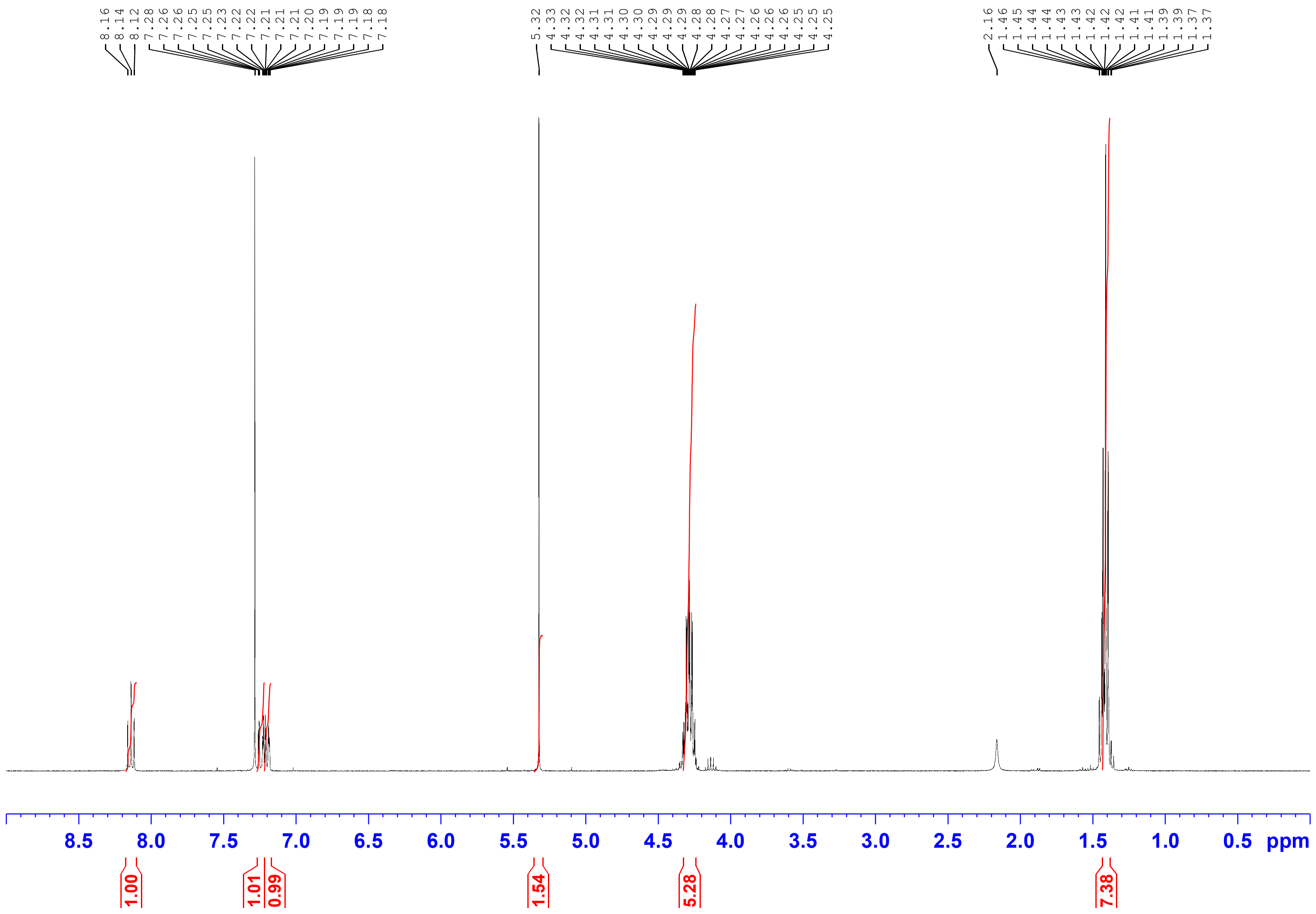

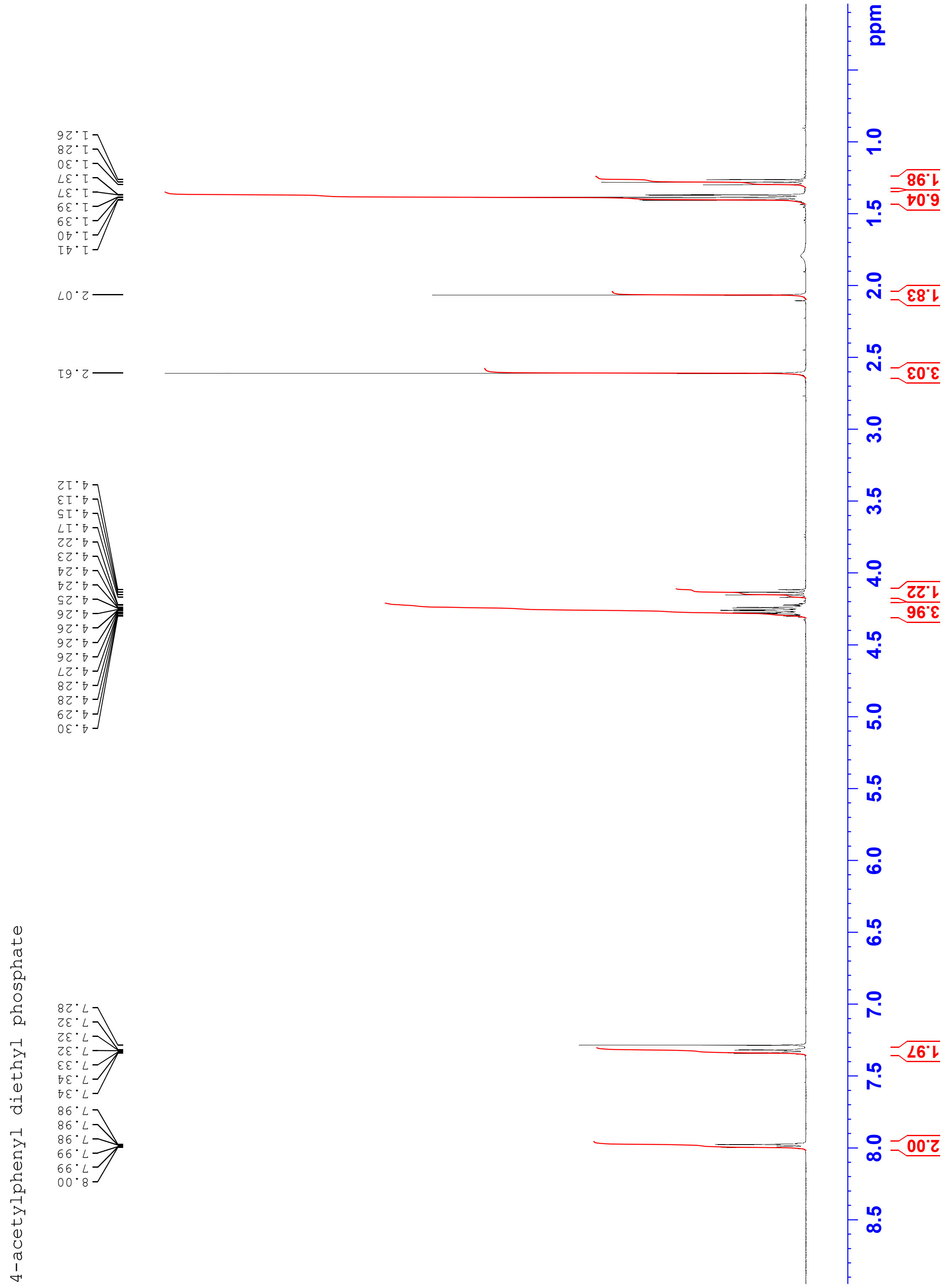
4-acetylphenyl diethyl phosphate
8.00
7.99
7.99
7.98
7.98
7.98
7.34
7.34
7.33
7.32
7.32
7.32
7.28
4.30
4.29
4.28
4.28
4.27
4.26
4.26
4.26
4.26
4.25
4.24
4.24
4.23
4.22
4.17
4.15
4.13
4.12
2.61
2.07
1.41
1.40
1.39
1.39
1.37
1.37
1.30
1.28
1.26
2.00
1.97
3.96
1.22
3.03
1.83
6.04
1.98
ppm

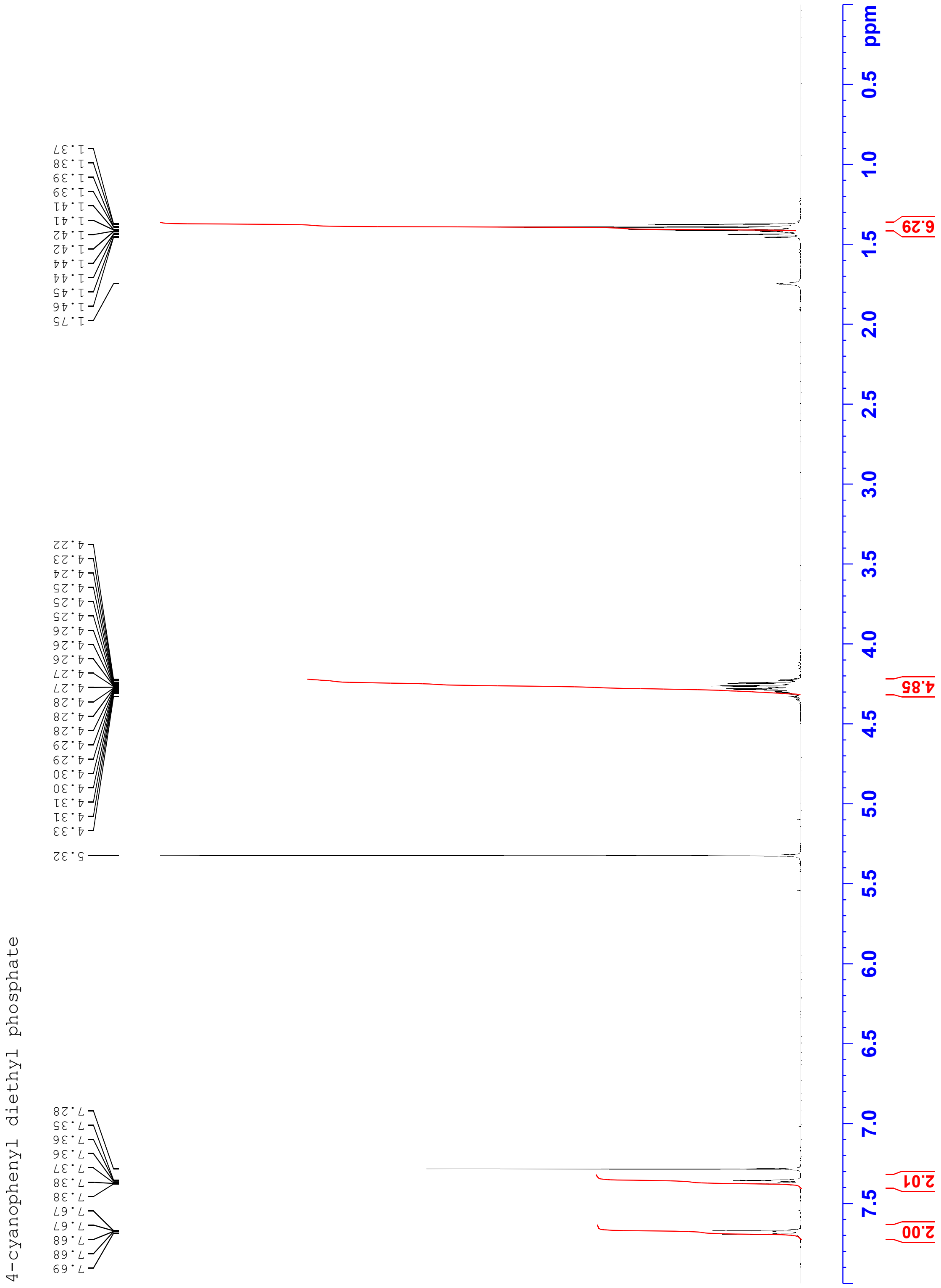
4-cyanophenyl diethyl phosphate
1.37
1.38
1.39
1.39
1.41
1.41
1.42
1.42
1.44
1.44
1.45
1.46
1.75
4.22
4.23
4.24
4.25
4.25
4.25
4.26
4.26
4.26
4.27
4.27
4.28
4.28
4.28
4.29
4.29
4.30
4.30
4.31
4.31
4.33
5.32
7.28
7.35
7.36
7.36
7.37
7.38
7.38
7.67
7.67
7.68
7.68
7.69
ppm
0.5
1.0
1.5
2.0
2.5
3.0
3.5
4.0
4.5
5.0
5.5
6.0
6.5
7.0
7.5
6.29
4.85
2.01
2.00

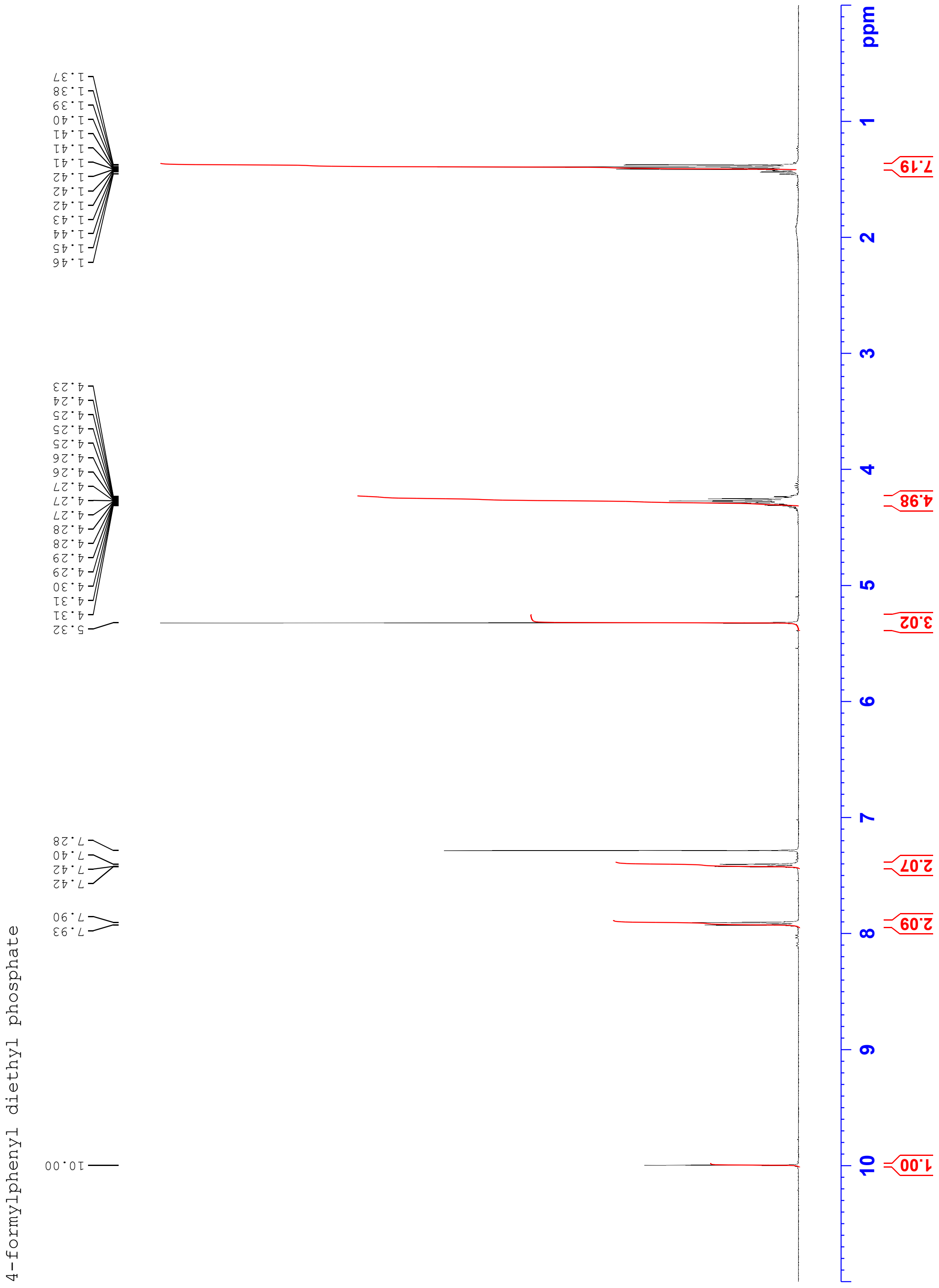
4-formylphenyl diethyl phosphate
10.00
7.93
7.90
7.42
7.42
7.40
7.28
5.32
4.31
4.31
4.30
4.29
4.29
4.28
4.28
4.27
4.27
4.27
4.26
4.26
4.25
4.25
4.25
4.24
4.23
1.46
1.45
1.44
1.43
1.42
1.42
1.42
1.41
1.41
1.41
1.40
1.39
1.38
1.37
1.00
2.09
2.07
3.02
4.98
7.19
ppm

(10) Gasteiger, E. Protein Identification and Analysis Tools on the ExPASy Server.

(11) R Core Team. *R: A Language and Environment for Statistical Computing*; R Foundation for Statistical Computing: Vienna, Austria, 2017.
